# Recurrent loss of an immunity gene that protects *Drosophila* against a major natural parasite

**DOI:** 10.1101/2022.05.27.493757

**Authors:** Ramesh Arunkumar, Shuyu Olivia Zhou, Jonathan P. Day, Sherifat Bakare, Simone Pitton, Chi-Yun Hsing, Sinead O’Boyle, Juan Pascual-Gil, Belinda Clark, Rachael J. Chandler, Alexandre B. Leitão, Francis M. Jiggins

## Abstract

Polymorphisms in immunity genes can have large effects on susceptibility to infection. To understand the origins of this variation, we investigated the genetic basis of resistance to the parasitoid wasp *Leptopilina boulardi* in *Drosophila melanogaster*. A *cis*-regulatory polymorphism in the gene *Lectin-24A* abolishes expression after infection, strongly reducing survival. Other null mutations have arisen repeatedly in this gene, with additional loss-of-expression and premature stop codons segregating in nature. The frequency of these alleles varies greatly, and in some populations natural selection has driven them near to fixation. We conclude that there is a pleiotropic cost to *Lectin-24A* expression, and in some populations this outweighs the benefit of resistance, resulting in natural selection causing the repeated loss of this important immune defense.

**Significance Statement:** Genetic differences between individuals can have a large effect on susceptibility to infectious disease. We have identified a gene called *Lectin-24A* that is important in the immune response that protects fruit flies against one of their main natural enemies—parasitic wasps. However, in nature many flies carry mutated copies of this gene that are no longer functional. We found that the high frequency of these loss-of-function mutations can only be explained if they have a selective advantage in some populations. Therefore, we can conclude that this immune defiance is costly, and genetic variation in susceptibility is maintained because in some locations susceptible flies are fitter than resistant flies.

## Introduction

Parasites can impose strong selection on host populations, driving resistance alleles up in frequency when infection is common. In Africa and Southeast Asia, where malaria-causing *Plasmodium* parasites have high transmission, multiple resistance alleles have undergone selective sweeps and become fixed (*1*). Similarly, genes that protect against Lassa hemorrhagic fever virus are at high frequency in Nigeria where the disease is found (*2*).

Despite the advantages of resistance, genetic variability in susceptibility to infection is abundant in humans (*3,4*), plants (*5,6*) and insects (*7*-*10*). The polymorphisms underlying this variation may be transient as resistant alleles are spread through populations, or they can be maintained by processes such as temporal and spatial differences in selection pressures (*11,12*) or negative frequency dependent selection (*13*).

Variability in susceptibility can be maintained in populations when resistance trades-off with other fitness-related traits. These costs may occur in the absence of infection (*14*) due to the diversion of resources from growth and reproduction (*15,16*), autoimmune damage (*17*), or when resistance to one pathogen increases susceptibility to a different pathogen (*18*).

However, not all resistance alleles are costly (*19*), and overtime compensatory mutations that reduce or negate fitness costs may spread (*20,21*). Alternatively, fitness costs could be avoided by reverting to susceptibility when the pathogen pressure is low (*22*).

One of the first demonstrations of the costs of evolving resistance came from *Drosophila* and parasitoid wasps, where populations selected for increased resistance had reduced competitive ability (*23,24*) and lower feeding rates (*25*). Female parasitoid wasps lay their eggs inside the larvae of *Drosophila*, and if the host is unable to mount a successful immune response, the parasitoid larva feeds on the host tissue and eventually kills it. Flies can kill parasitoid wasps through a cellular immune response known as encapsulation, in which hemocytes (blood cells) are recruited to the parasitoid egg and surround it (*26*). The capsule is then melanized, killing the parasitoid egg. Despite parasitoids being common in nature, there is considerable variation within and between populations in susceptibility to the parasitoids *Asobara tabida* (Braconidae) (*27*) and *Leptopilina boulardi* (Figitidae) (*28*). Early work found polymorphisms in two regions of the genome that are involved in resisting parasitoid infection: one for resistance against *L. boulardi* and the other against *A. tabida* (*29*-*32*). However, a recent study did not recover the same loci (*33*). In the light of these observations, we have attempted to identify the genetic basis of resistance against *L. boulardi* in a natural population of *Drosophila melanogaster*.

## Results

### Resistance to parasitoid infection is associated with a faster immune response

We identified two inbred lines, DGRP-437 and DGRP-892, that originated from the same wild population in the USA but had a marked difference in their ability to survive parasitoid infection (Figure 1A; binomial GLMM, *z*=-7.6, *p*<0.001). This difference is caused by varying anti-parasitoid defense response, which involves the wasp being encapsulated by immune cells called hemocytes and melanized. In the resistant line DGRP-437, the melanization phenotype becomes apparent at 24 hours and the wasp embryo is completely melanized at 26 hours (Figure 1B). In contrast, no melanization is seen in the susceptible line DGRP-892 in the first 26 hours post infection (Figure 1B).

**Figure 1.**
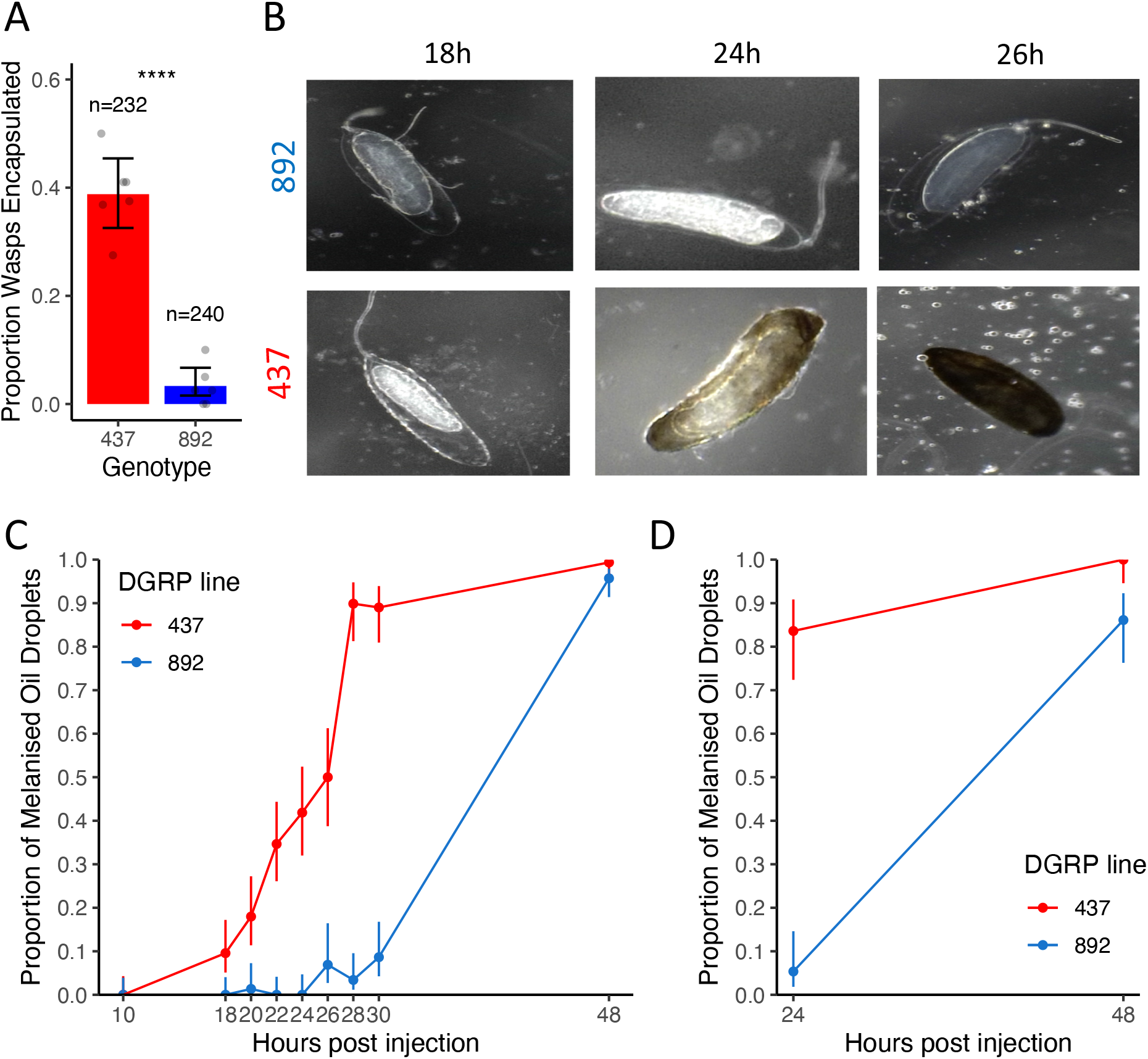
Genetic differences in the immune response to parasitoid infection. (**A**) The proportion of *Leptopilina boulardi* wasp embryos melanized in two inbred lines, DGRP-437 and DGRP-892. Each point is an independent replicate (6 experimental assay per line), the bars are 95% confidence intervals, and the total number of larvae across all assays is above the bar. (**B**) Wasp embryo encapsulation at 18, 24 and 26 hours after infection. (**C** and **D**) Larvae were injected with oil droplets containing homogenized *L. boulardi* (C, n_DGRP-437_ = 853, n_DGRP-892_ = 816) or *Asobara tabida* (D, n_DGRP-437_ = 128, n_DGRP-892_ = 128). Bars represent standard errors.

It has been proposed that this immune response must be fast to succeed, because once the wasp larva emerges from the egg chorion 24-48 hours post infection (hpi), it is mobile and better able to escape the cellular capsule (*34*). Parasitoids can suppress host immunity by injecting venoms along with the egg. To examine the speed of the immune response in the absence of this immune suppression, we triggered the immune response by injecting larvae with droplets of mineral oil containing homogenized parasitoid wasp (Figure 1B). Both *Drosophila* lines had melanized the oil droplets by 48 hours post injection (hpi). However, the resistant line mounted this immune response faster—89.8% of the oil droplets were melanized at 28 hpi compared to 3.4% in the susceptible line (binomial GLM, logistic regression χ^2^=432.1, *df*=1, *p*<0.001). This difference is not specific to the parasitoid *L. boulardi*, as we obtained similar results when injecting oil droplets containing homogenate of the parasitoid *A. tabida* (Figure 1C; binomial GLM, genotype: logistic regression χ^2^=97.3, *df*=1, *p*<0.001; time post injection: logistic regression χ^2^=111.5, *df*=1, *p*<0.001).

To investigate how hemocytes contribute to the different encapsulation responses of the resistant and susceptible lines, we crossed them to flies that express florescent proteins in hemocytes. Msn-mCherry is expressed specifically in lamellocytes (*35*), while HmlΔ-GAL4 UAS-GFP is expressed in nearly all hemocytes except mature lamellocytes, and therefore marks plasmatocytes. The F_1_ progeny of this cross differed in their ability to melanize wasp eggs and oil droplets depending on whether they had a resistant or susceptible parent (Figure S1A and S1B; Fisher’s exact test, wasp eggs: *p<*0.001; oil droplet: *p<*0.001). However, when the wasps or oil droplets were dissected, lamellocytes and plasmatocytes could be seen attached to both the oil droplet and wasp egg in both lines (Figure S1C). Therefore, the hemocytes of the susceptible line can form a cellular capsule, although it is possible that differences in the timing or magnitude of this response may cause differences in the melanization response.

### Resistance results from epistatic interactions between genes on different chromosomes

We next investigated the genetic basis of resistance. When we crossed the resistant and susceptible lines the F_1_ progeny were highly resistant, indicating that resistance was dominant (Figure 2A). By swapping whether the resistant parent was the mother or father, we generated male offspring that only differed in their X chromosome (Figure 2A). Neither the sex of progeny (Likelihood ratio test, χ^2^=3.46, *p*=0.06) nor the X chromosome genotype (Tukey’s HSD, *z*=0.45, *p*=0.65) had a significant impact on susceptibility (Figure 2A). Together, these results demonstrate that resistance is an autosomal dominant trait.

**Figure 2.**
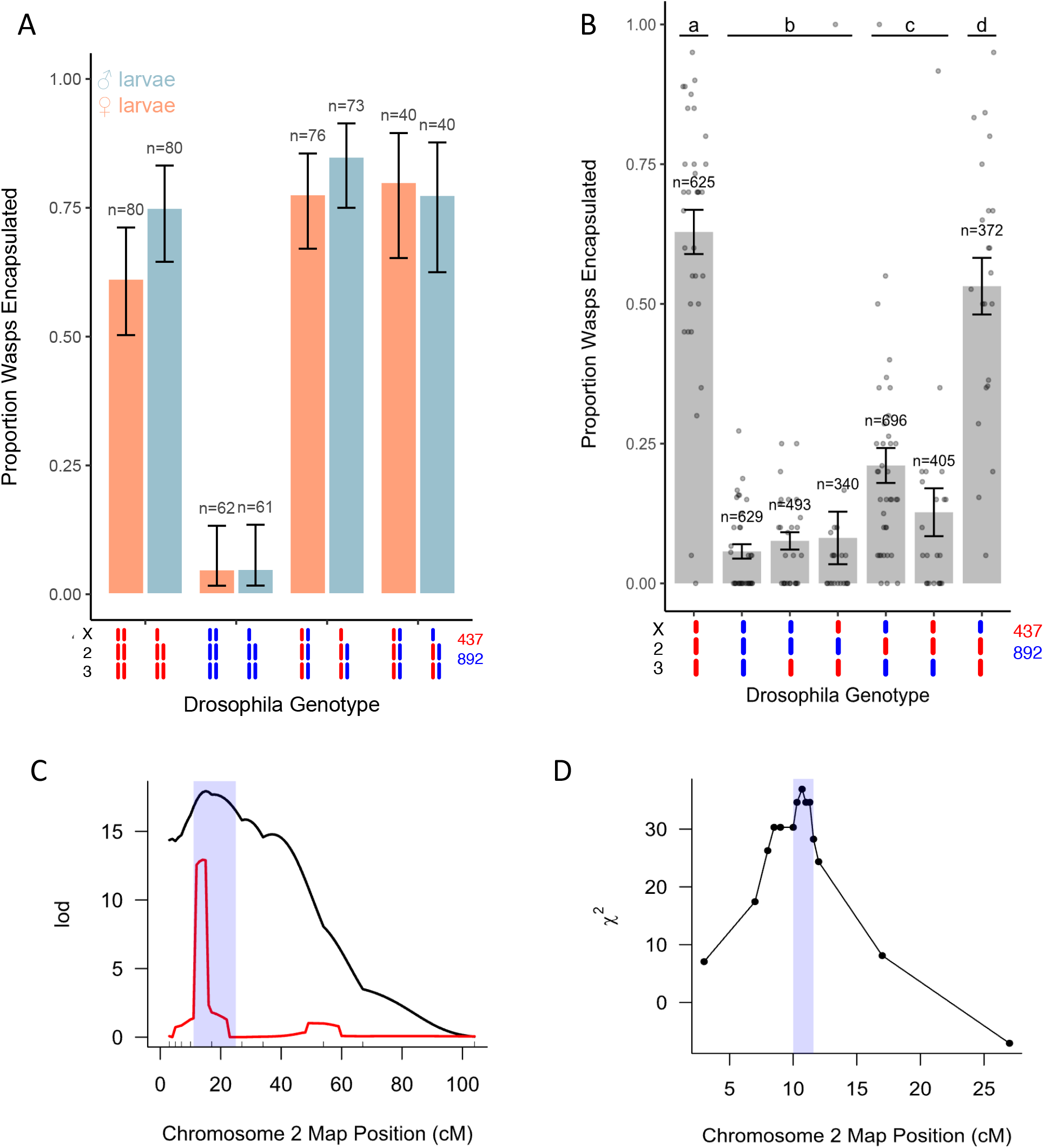
A single locus on chromosome II is associated with resistance to parasitoid wasp infection. (**A**) The mean proportion of wasps encapsulated in the F_1_ progeny of crosses between two *Drosophila* lines. Error bars are 95% confidence intervals. (**B**) The mean proportion of wasps encapsulated in isogenic lines with different chromosome combinations. Bars are standard errors and points are replicates of 20 larvae. Letters show significantly different groups (Tukey’s test with Holm correction, *p*<0.05 between groups) (**C**) QTL on chromosome II associated with the encapsulation rate measured by dissecting larvae. The black line is interval mapping and the red line composite interval mapping. The blue shaded region is the confidence interval on the location of the QTL (1.5 LOD drop). Ticks above the X axis are marker locations. (**D**) Fine-scale QTL map of region containing the gene. Only informative recombinants were genotyped. The χ^2^ statistic represents the deviation from the expected Mendelian frequency (0.5) of each marker among 152 adults that contained a capsule and tested positive for parasitoid wasp DNA. The shaded region indicates the 95% confidence interval (χ^2^ drop of 4.6).

To identify the chromosomes affecting susceptibility to infection, we generated lines carrying varying combinations of the X, II and III chromosomes (Figure 2B). When comparing lines that differ only in their second chromosome, having chromosome II from the resistant parent always resulted in greater levels of encapsulation (Figure 2B). However, the magnitude of this effect depends on genes elsewhere in the genome. When paired with a third chromosome from the resistant parent, swapping chromosome II could convert a fully resistant line into a fully susceptible line (Figure 2B). However, when paired with a thirdfalls at the peak of the QTL. As lectins are important chromosome from the susceptible parent, chromosome II only had a small effect. This is reflected in a statistical interaction between the two chromosomes (GLM with logit link, Wald Chi-square test: χ^*2*^=24.7, *p*<0.001), indicating that there is a multiplicative epistatic interaction between the second and third chromosome.

### A major effect locus on chromosome II affects resistance

We used quantitative trait locus (QTL) mapping to locate the region on the second chromosome affecting susceptibility to infection. We crossed fly lines that differed only in the second chromosome (X^892^; II^892^; III^437^ and X^892^; II^437^; III^437^), then backcrossed the F_1_ progeny to the susceptible parent. The resulting larvae were parasitized, and we then genotyped 386 individual larvae using 10 molecular markers spanning the second chromosome. We identified a single region where the chromosome II genotype was associated differences in susceptibility (Figure 2C, black line). Based on a 1.5 logarithm of the odds (LOD) drop, the QTL encompassed 11 to 25 cM (Figure 2C, blue box). Composite interval mapping, which searches for additional QTL while accounting for the main peak, indicated that there is a single locus on chromosome II affecting susceptibility (Figure 2C, red line).

As this QTL contained many genes, we conducted a second round of genetic mapping using only flies that were recombinant within the QTL. We repeated the genetic cross, parasitized the backcrossed larvae, and selected adults that contained visible capsules in their body and had therefore likely survived infection (the flies that did not resist infection were killed). We genotyped these individuals using molecular markers flanking the QTL (3 and 27 cM) to identify recombinants. Out of 1,486 adults, 298 had a recombination breakpoint between 3 and 27 cM—a recombination fraction of 0.20. We genotyped twelve molecular markers within this region for 152 individuals where we could amplify wasp DNA to confirm they had been infected. As we only genotyped resistant flies, we tested whether each marker allele frequency departed from the Mendelian expectation of 50:50 (we checked there was no segregation distortion by genotyping 348 uninfected flies).

Using this approach, we identified a single QTL at 10.3cM on the left arm of chromosome II (Figure 2D). By simulating 1000 replicate datasets we estimated that the 95% confidence interval on the location of the QTL contained 84 protein coding genes and 23 long non-coding RNAs (10.0-11.6 cM, genome version 6: 3.43-4.03 Mbp, Table S1). Among the 1,188 non-recombinant flies, 906 carried the resistant allele and 282 the susceptible allele, indicating a risk ratio of 3.21 (homozygous susceptible versus heterozygotes).

### A C-type lectin underlies resistance

Infection by parasitoid wasps induces a large transcriptional response in the two main immune tissues of *Drosophila—*the fat body and hemocytes. Using our unpublished RNA sequencing (RNA-Seq) data (*36*), we searched for genes within the QTL that were upregulated after anti-parasitoid immune induction in these tissues (log_2_ fold change>2). We found that 8 genes were differentially expressed in hemocytes and one gene, *Lectin-24A*, in the fat body (Figure 3A). *Lectin-24A* has previously been found to be massively upregulated following parasitoid wasp infection (*37*-*39*) and falls at the peak of the QTL. As lectins are important receptors in innate immune systems, we focused on this gene.

**Figure 3.**
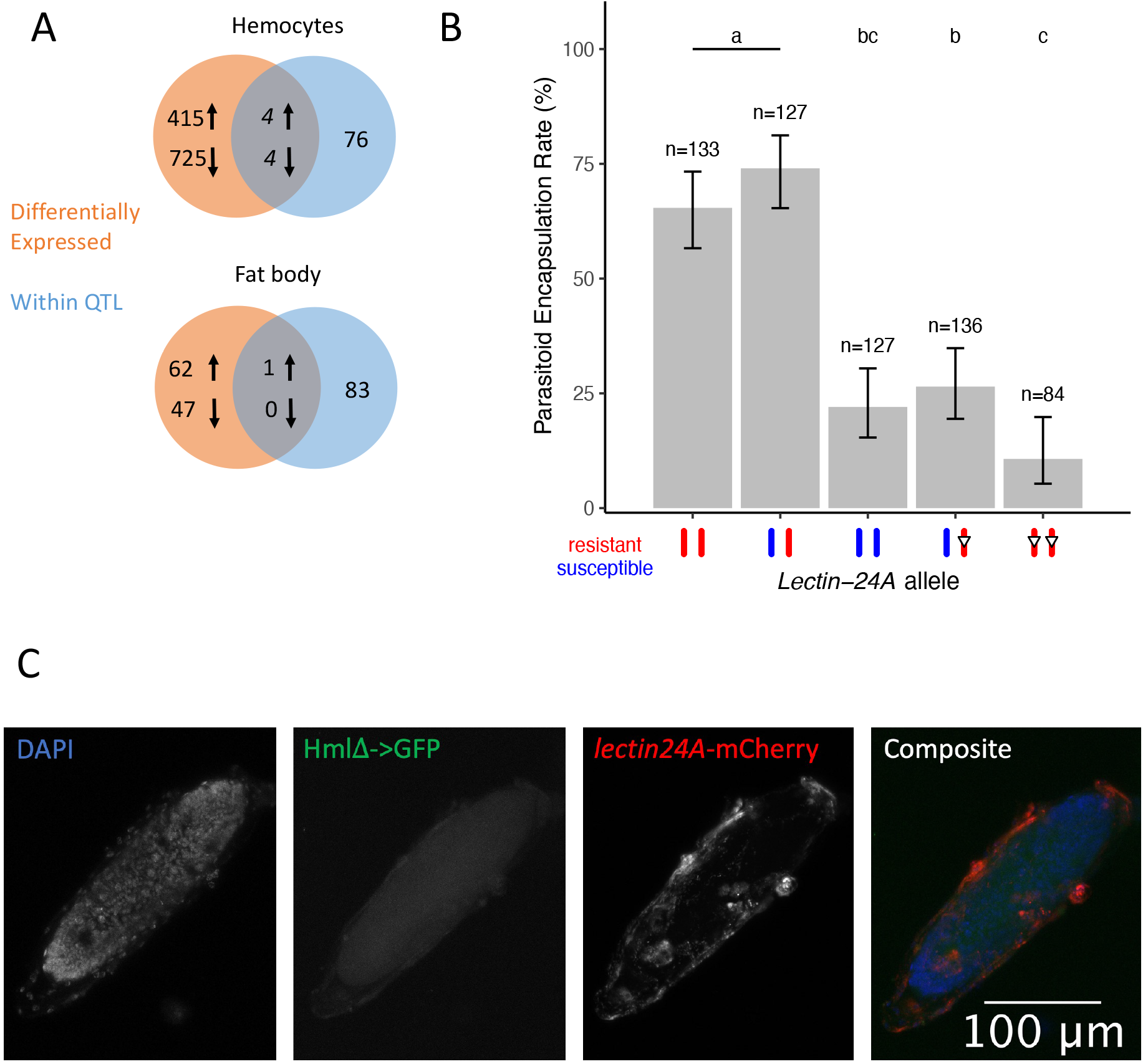
*Lectin-24A* is required for parasitoid resistance. (A) Overlap between protein-coding genes that are within the QTL and differentially expressed after immune induction (log_2_ fold change > 2). The gene in the intersect of the Venn diagram in the fat body is *Lectin-24A*. (B) The mean parasitoid encapsulation rate in lines with resistant, susceptible, and mutated (open inverted triangles, *Δ129*) *Lectin-24A* alleles. Error bars represent 95% confidence intervals. Tukey’s test, *p*<0.05 between groups. (C) Confocal microscopy imaging of wasp egg dissected 23 hours post-infection from a heterozygous F_1_ larva produced from the cross of virgins expressing *Lectin-24A* tagged with mCherry and males expressing HmlΔ-GAL4 UAS-GFP. At this time point, plasmatocytes have yet to spread on the wasp egg, while *Lectin-24A* can be observed to bind to the surface of the egg.

To test if *Lectin-24A* is necessary for resistance to parasitoid wasps, we mutated the resistant allele using CRISPR-Cas9 in the resistant X^892^; II^437^; III^437^ flies. This resulted in a mutated allele with a 4bp insertion that was 129bp downstream of the start codon, which we named *Lectin-24A*^*Δ129*^. The change in reading frame introduces a premature stop codon and abolishes the carbohydrate binding domain of the protein. This mutation made flies susceptible to infection (Figure 3B; Tukey’s HSD test: *p*<0.001), with heterozygote flies containing one mutated resistant allele and one susceptible allele having similar encapsulation rates as flies with two susceptible alleles.

Lectin-24A is a C-type lectin, a family of proteins which frequently act as pattern recognition receptors in the innate immune system due to their ability to bind specifically to ligands (*40*). To investigate the role of *Lectin-24A* in the immune response, we created transgenic flies that expressed *Lectin-24A* fused to a fluorescent protein under the control of the gene’s native promoter. When larvae from these lines were infected by a parasitoid, the protein localized to the surface of the wasp egg before it was encapsulated by hemocytes (Figure 3C). This is consistent with this molecule being an opsonin involved in the initial recognition of the parasite, guiding the subsequent cellular immune response.

C-type lectins are named due to their ability to bind to specific carbohydrates in a calcium-dependent manner (*40*). By aligning the peptide sequence of the Lectin-24A carbohydrate recognition domain with other members of the protein family, we found that residues required for the interaction with the calcium ions have been lost, suggesting it is not involved in calcium-dependent carbohydrate binding (Figure S2A). This is reminiscent of another well-characterized group of C-type lectins that have lost the ability to bind calcium—natural killer cells receptors—which bind ligands including proteins (*41*). Alongside conserved cysteines involved in forming the Ca^2+^ binding site, Lectin-24A contains an additional cysteine within the carbohydrate domain and four cysteines elsewhere in the protein (Figure S2A), suggesting it may form multimers. By expressing affinity tagged Lectin-24A in *Drosophila* cells, we confirmed the protein forms tetramers (Figure S2B).

### A *cis* regulatory polymorphism in *Lectin-24A* is associated with resistance

We used quantitative PCR (qPCR) to examine whether the resistant and susceptible alleles of *Lectin-24A* differed in their expression (Figure 4A). In uninfected larvae, the resistant allele is expressed at a higher level than the susceptible allele. Furthermore, after infection there was a 5-15 fold upregulation of the expression of the resistant allele, but the susceptible allele was not induced (6 and 18 hpi, Figure 4A; ANOVA, effect of *Drosophila* line: *F*=163.96, *df*=1, *p*<0.001).

**Figure 4.**
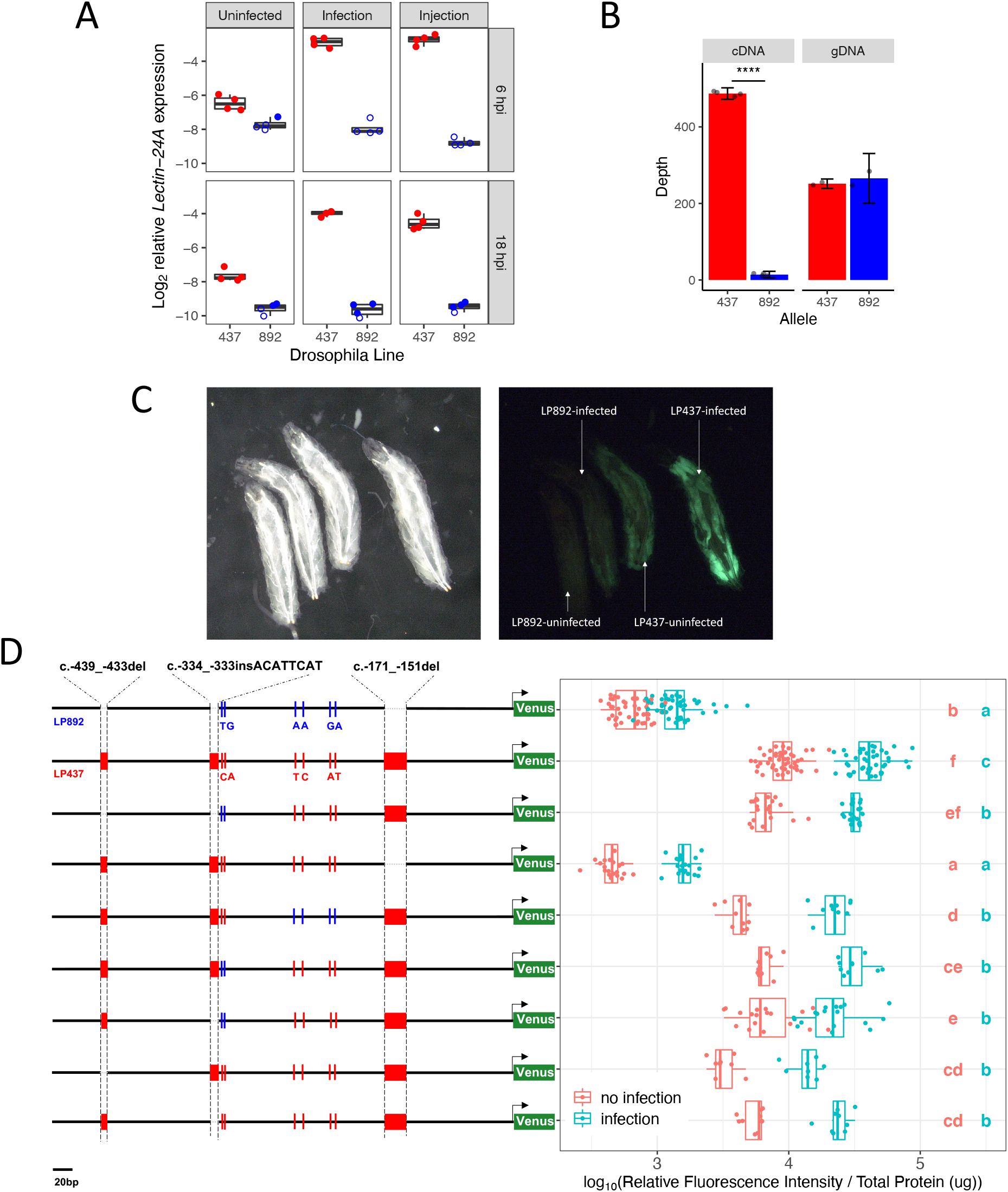
*Cis*-regulatory polymorphisms in *Lectin-24A*. (A) Expression of *Lectin-24A* 6 and 18 hours post infection with parasitoid wasp or injection with an oil droplet containing wasp. Open circles indicate gene expression was below the detection threshold of that sample. (B) Allele-specific expression of *Lectin-24A* in heterozygous flies. Read counts (depth) of the resistant (437) and susceptible (892) allele for complementary DNA (cDNA) and genomic DNA (gDNA) libraries. Asterisks indicate a Welch *t-*test, *p*<0.0001. (C) Expression of Venus driven by the sequence upstream of the susceptible allele of *Lectin-24A* (LP892) or the resistant allele (LP437). Infected or uninfected larvae were imaged 24 hours after infection under bright-field (left) or GFP filter (right). (D) Reporter constructs with different combinations of indels and SNPs, designed based on the upstream region of *Lectin-24A* in the susceptible (892, blue) and resistant (437, red) lines and expression of the Venus protein. Each point on the plot represents an independent sample of 15 larvae. Letters are Tukey’s test, *p*<0.05 between groups, *p*>0.05 within groups.

The differences in *Lectin-24A* expression could be controlled in *cis* or *trans*. To distinguish these two possibilities, we crossed the two lines, infected them, and Illumina-sequenced the *Lectin-24A* transcript in the heterozygous F_1_ progeny. The sequence reads were assigned to the resistant and susceptible alleles using SNPs that differ between the two alleles, allowing us to measure their relative expression. In these heterozygous flies, we found that the expression of the resistant allele was 34 times greater than the susceptible allele (Figure 4B; Welch two sample t-test: *t*=135.6, *df*=4.7737, *p*<0.001). As the two alleles are present in the same cells they share the same *trans*-regulatory environment, these differences in expression are controlled in *cis*.

Many *cis* regulatory elements are found a short distance upstream of the gene they control. The *Lectin-24A-mCherry* transgene described above included 489bp of sequence upstream of the start codon of the resistant allele, and this is strongly induced after infection (Figure S3). However, when the equivalent transgene was made from the susceptible allele there was no detectable expression (Figure S3). To confirm this result, we cloned these regulatory sequences in front of GFP to create fluorescent reporters that were inserted into the *Drosophila* genome. Recapitulating the results from the *Lectin-24A-mCherry* construct, the sequence upstream of resistant allele drove strong expression of the reporter in the fat body but the equivalent sequence upstream of the susceptible allele did not (Figure 4C). To accurately quantify expression, we measured fluorescence in protein extracted from larvae. Confirming the microscopy (Figure 4C), we observed a ∼30-fold difference in fluorescence between the reporter lines carrying the regulatory sequence of the resistant and susceptible alleles of *Lectin-24A* (Figure 4D, top two constructs). In contrast to the expression of *Lectin-24A* itself (Figure 4A), the *cis* regulatory sequence from both alleles upregulated expression after infection (Figure 4D), despite the resistant allele always having higher expression. Together, this demonstrates this region contains a *cis* regulatory polymorphism that differs between the resistant and susceptible alleles.

Comparing the resistant and susceptible alleles, the *cis* regulatory sequence used in the reporter constructs differed by three insertion-deletion polymorphisms (indels) and six single nucleotide polymorphisms (SNPs; Figure 4D, top two rows). To identify which of these causes the differences in expression, we created seven more transgenic fly lines, each carrying a reporter construct that had a different combination of alleles at these sites (Figure 4D). When we introduced a 21bp indel (c.-171_-151del) found upstream of the susceptible allele into the resistant allele reporter, it greatly reduced the expression levels (Figure 4D). In contrast, swapping alleles of the other polymorphic sites frequently resulted in minor but statistically significant changes in expression (∼3 fold). Therefore, we can conclude that the 21bp indel (c.-171_-151del) is primarily responsible for the differences in the expression of *Lectin-24A* in the resistant and susceptible lines.

### Loss-of-function alleles have arisen repeatedly in natural populations

We next investigated genetic variation in *Lectin-24A* expression at the population level. We first sequenced a 557bp region upstream of the gene the panel of inbred lines from North America (*42*) (Table S2) and selected 20 lines with different haplotypes at the three indels shown in Figure 4D. We crossed these to our resistant line (DGRP-437) and Illumina-sequenced the *Lectin-24A* transcript in the F_1_ progeny to look for evidence of allele-specific expression (ASE). This assay produced consistent results across replicates (Figure S4A, S4B) and when we sequenced genomic DNA (gDNA) the frequency of reads from the two alleles was close to 0.5 (Figure 5A). However, when we sequenced transcripts (cDNA) there was considerable variation in the expression of the different alleles, and this was perfectly associated with the indel haplotype (Figure 5A). As expected, the four lines carrying the 21bp deletion (c.-171_-151del) all had very low expression (Figure 5A, haplotype DDD).

**Figure 5.**
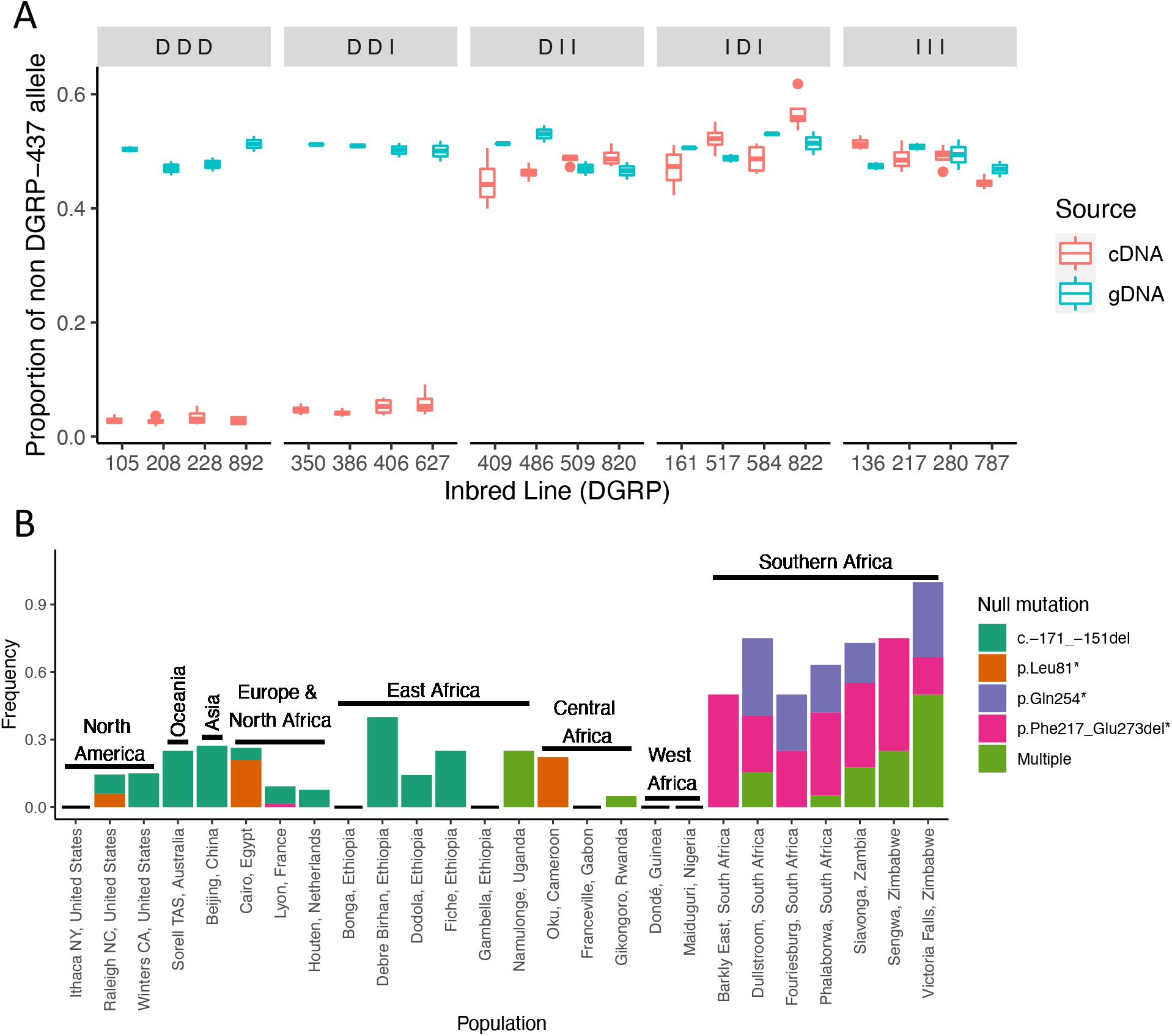
The frequency of predicted loss-of-function alleles of *Lectin-24A* in natural populations. (A) To identify variation in *cis* regulatory elements affecting gene expression, *Lectin-24A* complementary DNA (cDNA) and genomic DNA (gDNA) was sequenced in the F_1_ progeny of a cross between DGRP-437 and 20 different inbred DGRP lines. The lines are grouped by their indel haplotype, which is ordered as c.-439_-433del, c.-334_- 333insACATTCAT and the 21bp indel (c.-171_-151del) and the deletion ‘D’ or insertion ‘I’ state for each indel is depicted. For each of the 20 genotypes we estimated the frequency of reads from the test line as opposed to DGRP-437. We had two technical and two biological replicates for each cross and sampled larvae 24 hours post infection with parasitoid wasp.Data for line 892 is also presented in Figure 3. (B) The frequency of variants that either abolish expression (the 21bp indel c.-171_-151del) or introduce premature stop codons (other variants). Multiple refers to the cooccurrence of any two loss-of-function variants. Analysis used 672 genome sequences genotyped for >50% of the *Lectin-24A* gene region.

However, four lines with the insertion allele at this site also had strongly reduced expression (Figure 5A, haplotype DDI). While expression in these lines was low, it was nonetheless 1.7 times higher than lines with the 21bp deletion (quasibinomial GLM, DDD vs DDI: *t*=4.319, *p*<0.001). To confirm these results, we also measured *Lectin-24A* expression in a sample of inbred lines by qPCR. These results broadly supported our conclusion that lines with the 21bp deletion had the lowest expression, but expression also tended to be reduced in lines with the DDI haplotype (Figure S4C). Together, 10 of the 150 lines that were fully genotyped carried one of the low expression haplotypes (DDD or DDI). After removing the lines with the 21bp deletion, there were four SNP in the 1,000bp region upstream that are perfectly associated with low gene expression and are therefore candidate *cis*-regulatory polymorphisms (Figure S5). In conclusion, there are multiple common *cis* regulatory polymorphisms segregating in natural populations that can largely abolish *Lectin-24A* expression.

As multiple variants cause the loss of *Lectin-24A* expression, we next examined whether loss-of-function variants are segregating in the protein-coding sequence of the gene. It has previously been reported that there are alleles of *Lectin-24A* containing premature stop codons in natural populations (*43*). As many more genomes have been sequenced since that analysis, we searched 1,039 published genomes from flies collected around the world for variants that are likely to result in null alleles of *Lectin-24A* (*44*). We identified a 165bp deletion in the protein coding sequence that resulted in a shift in the reading frame and a premature stop codon (p.Phe217_Glu273del*), and three additional point mutations that introduced premature stop codons either within or before the carbohydrate recognition domain of the Lectin-24A protein. Lines containing these premature stop codons were able to upregulate *Lectin-24A* following parasitoid wasp infection (Figure S6), suggesting that these variants cause the loss of gene function independently of the loss-of-expression mutations.

To understand the geographical distribution of putative loss-of-function alleles in *Lectin-24A*, we examined their frequency in 26 populations. Southern African populations have the highest frequency of loss-of-function alleles, with over half of alleles carrying a premature stop codon, and many alleles carrying more than one of such mutations (Figure 5B). Outside of southern Africa, premature stop codons are rare, but the 21bp deletion (c.-171_-151del) that abolished expression is widespread, reaching frequencies over 30% in some locations. We note that because there are additional loss-of-expression polymorphisms that we cannot identify from sequence alone (see above), these numbers underestimate the true frequency of loss-of-function alleles.

### Natural selection has driven the loss of *Lectin-24A*

Given the importance of *Lectin-24A* in defending flies against parasitoid wasps, the finding that non-functional alleles are common in nature is unexpected. We therefore explored the evolution of the gene in more detail. First, we examined whether non-functional alleles are the ancestral or derived states by aligning the *Lectin-24A* gene region with the homologous region from two closely related species—*D. simulans* and *D. sechellia*. Both species contain the 21bp sequence (AAATAAGGCTATCTGGGATCA; c.-171_-151del) that is required for the gene to be induced after infection. Furthermore, the coding sequence of the gene does not contain any premature stop codons. Therefore, in all cases the loss-of-function allele is the derived state.

The variable frequency of loss-of-function alleles in nature (Figure 5B) suggests that these alleles may be favored in some populations but not others. To explore how natural selection has acted on this gene in global fly populations, we used 1,039 published genome sequences (*44*). In line with previous analyses of a smaller dataset (*43*), multiple SNPs in *Lectin-24A* had very high levels of genetic differentiation, with several SNPs being among the 0.1% of variants in the genome with the greatest geographical variation in their frequency (Figure 6A). Pairwise comparisons between geographical regions showed that this pattern was driven by the southern African populations being highly differentiated from other geographic regions in the world (Figure 6B). Across the gene there are numerous variants that are near fixation across southern Africa but are rare elsewhere in the world (Figure S7). This was the result of a divergent haplotype of the gene that is common in southern Africa but rare elsewhere (Figure S8A, S8B). Two of the three premature stop codons and the coding sequence deletion (p.Phe217_Glu273del*) were at their highest frequency (16.7-56.9%) in southern Africa. The other premature stop codon (p.Leu81*) was not found in southern Africa, but was segregating at 4-5% in North America, Europe and North Africa and Central Africa.

**Figure 6.**
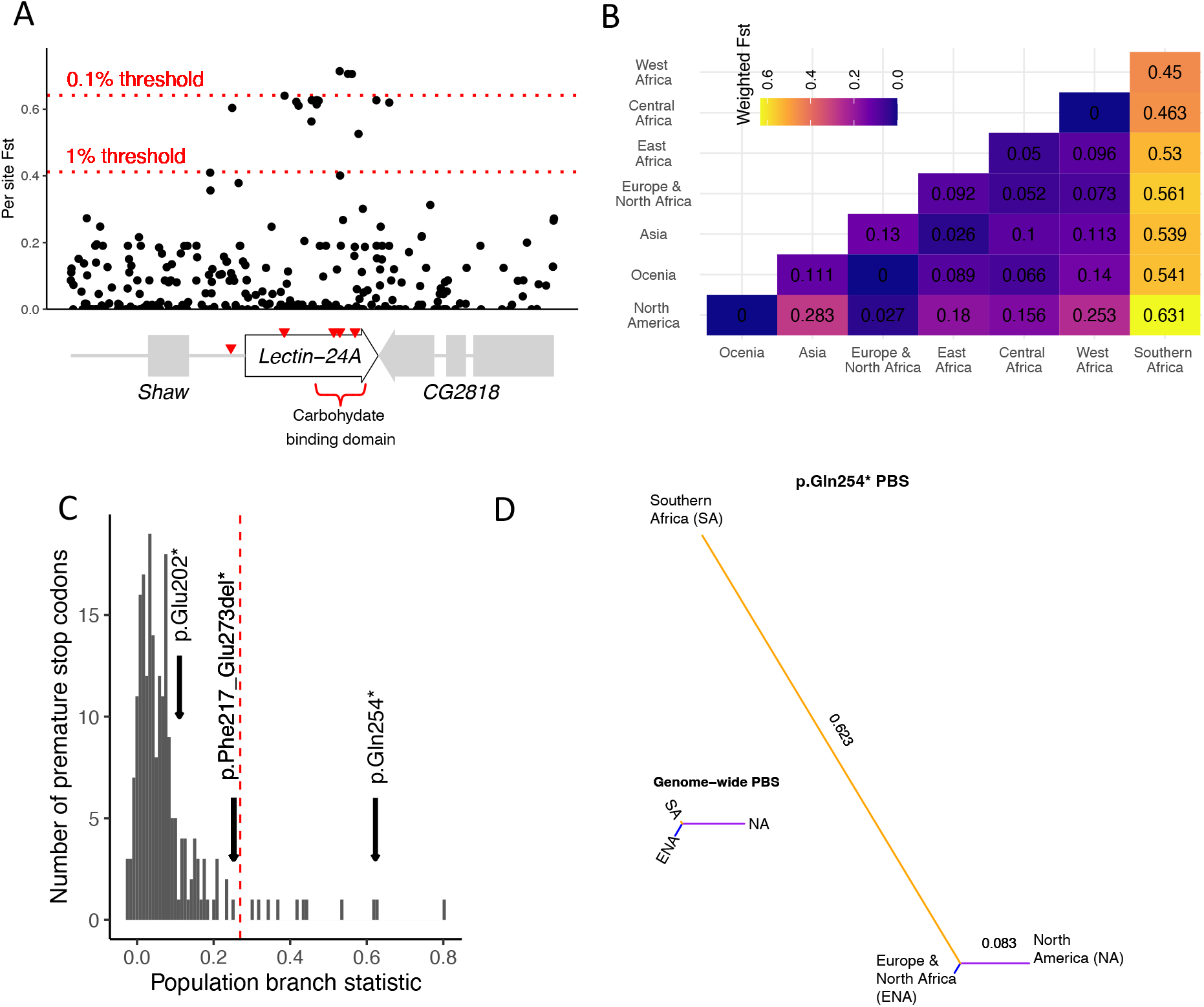
Evidence of local adaptation in *Lectin-24A*. (A) Per site Weir and Cockerham *F*_*ST*_ estimates for *Lectin-24A* and flanking regions (dm5; 2L:3715513–3719050) across 26 global fly populations. *F*_*ST*_ for the top 1% and 0.1% of autosomal SNPs is indicated by dotted red lines. The X axis shows the genes in the region. Red inverted triangles indicate loss-of-function variants. (B) Weir and Cockerham *F*_*ST*_ estimates for *Lectin-24A* protein coding sequence from pairwise comparisons between seven geographic regions. (C) Population branch statistic (PBS) for the three largest geographic regions in the *Drosophila* Genome Nexus. Histogram shows the PBS for the 211 premature stop codons found across the genome. The three *Lectin-24A* premature stop codons occurring within these populations are indicated with arrows. The red dotted line is the 95^th^ percentile. (D) A PBS tree for the polymorphic stop codon p.Gln254* and a PBS tree generated using pairwise *F*_*ST*_ across all 211 stop codons. Both trees are on the same scale.

As the loss of *Lectin-24A* has a large effect on susceptibility to infection, we tested whether natural selection has driven the premature stop codons to a high frequency in southern Africa using the population branch statistic (Figure 6C). This used pairwise *F*_*ST*_ estimates between the three regions with large numbers of published genomes (Europe & North Africa, North America or southern Africa) to generate a tree, with longer branches indicative of larger changes in allele frequency along that branch (*45*). To generate an empirical null distribution, we also calculated the population branch statistic for 211 other stop-gained mutations that occur at >5% frequency in at least one of the three regions (p.Leu81* was excluded as it <5% frequency in our samples). We found that p.Gln254* had an extremely high population branch statistic in southern Africa when compared to the genome-wide population branch statistic using all 211 stop codons, indicating that this loss-of-function variant is under positive selection in that region (Figure 6D).

## Discussion

In nature, *Leptopilina* wasps are capable of infecting and killing 90% of *Drosophila melanogaster* in some fruits in a single generation (*46*). Therefore, our discovery that numerous loss-of-function alleles are segregating in a gene that protects flies against these parasites is unexpected. Artificial selection experiments across multiple *Drosophila* species have shown that when populations evolve resistance to parasitoid infection, the fitness of uninfected flies is strongly reduced (*23,47*). Our population genetic analysis demonstrated that natural selection has favored null alleles of *Lectin-24A* in southern Africa, suggesting that this gene may contribute to these pleiotropic costs of parasite resistance.

Since maintaining resistance against parasitoid wasps is costly, loss-of-function mutations in resistance genes underlying this cost (i.e. susceptible alleles) may have a selective advantage in the absence of parasitoids. The ‘less is more’ view postulates that gene loss and pseudogenization can be beneficial, particularly following drastic shifts in environmental conditions (*48*). Adaptive loss-of-function has been implicated in the evolution of diverse organisms from bacteria evolving antibiotic resistance, yeast adapting to novel environmental challenges, mating system shifts in plants due to a breakdown of plant-pollinator interactions, evolution of herbivory in a *Drosophila* relative, and reduced risk to type 2 diabetes in humans (*49,50*). As natural selection specifically favored loss-of-function alleles of *Lectin-24A* in southern Africa, the balance of these costs and benefits appears to have shifted in different environments, resulting in the susceptible allele having an advantage. This may reflect differences in the parasite pressure. However, the cost of resistance is only apparent when food is scarce (*23,47*), so these geographical differences in selection may be due to the environment altering harmful pleiotropic effects of the resistant allele on some other trait.

*Lectin-24A* appears to be a hotspot of adaptive evolution in the *Drosophila* immune system (*43*). It has arisen recently in the common ancestor of *D. melanogaster* and *D. simulans* following gene duplication (*43*). It is one of the most rapidly evolving proteins in the genome, with most of the amino acid substitutions being fixed by natural selection (*43*). In *D. simulans* there has been a recent and strong selective sweep (*43*), while in *D. melanogaster* natural selection has caused exceptionally high levels of geographical variation in allele frequencies. Together, these observations suggest this gene may be a key player in the coevolution of *Drosophila* and parasitoids.

*Drosophila* kills parasitoids using a cellular immune response. However, parasitoid infection also triggers a strong transcriptional response in the fat body, resulting in the secretion of numerous humoral immune factors. The function of these molecules is largely unknown. *Lectin-24A* is massively upregulated following infection by *A. tabida* (*37,51*) and *L. boulardi* (*39*) but not following wounding or bacterial infection (*38*). We found that Lectin-24A localizes to the surface of the parasitoid egg before the attachment of hemocytes. It is possible that it is functioning as an opsonin, binding to molecules on the parasite surface to promote the attachment of hemocytes. An understanding of the molecular function of Lectin-24A may provide insights into why it evolves so fast, and why its presence appears to be costly to flies in some populations.

## Acknowledgements

We thank Kurt Drickamer for providing the C-type lectin carbohydrate recognition domain alignment. We thank John Pool for providing the SD and SP lines from the *Drosophila* Genome Nexus. We thank Dagmara Korona for sharing their pCFD5 and a plasmid containing the Flag-StrepII tag with us. We thank Angela Early for providing more detail on the genetic differentiation of *Lectin-24A* in the Global Diversity Lines. We thank Carlos Luque for assisting with *Drosophila* larval imaging.

## Funding

Natural Environment Research Council grant (NE/P00184X/1) awarded to Francis M. Jiggins and Alexandre B. Leitão. European Molecular Biology Organization fellowship (ALT-1556) awarded to Alexandre B. Leitão. Natural Sciences and Engineering Research Council of Canada fellowship (PDF-516634-2018) awarded to Ramesh Arunkumar. Shuyu O. Zhou was supported by the Gates Cambridge Trust. Juan Pascual-Gil was supported by an Erasmus+ traineeship and Belinda Clark was supported by the Balfour-Browne Fund from the University of Cambridge Zoology department.

## Author contributions

Conceptualization: FMJ, ABL, RA

Methodology: FMJ, RA, SOZ, JPD, ABL

Investigation: RA, SOZ, JPD, SB, SP, RJC, CYH, JPG, SO, BC

Visualization: RA, SOZ, FMJ, JPD, SB, SP

Funding acquisition: FMJ, ABL

Project administration: FMJ

Supervision: RA, FMJ, SOZ, ABL, JPD

Writing – original draft: RA, FMJ, SOZ, JPD

Writing – review & editing: RA, FMJ, SOZ, JPD, ABL, SB, SP, RJC, CYH, JPG, SO, BC

## Competing interests

Authors declare that they have no competing interests.

## Data and code availability

Miseq reads for cDNA and gDNA for ASE of *Lectin-24A* were deposited into the NCBI Sequence Read Archive under Bioproject PRJNA789229. The DGRP-437 *Lectin-24A* coding gene sequence and the Cas9 mutant sequence were deposited into GenBank: OM100576-OM100577. Scripts and processed data files are available in Github [DOI: 10.5281/zenodo.6543576].

## Notes

https://doi.org/10.5281/zenodo.6543576

### Competing Interest Statement

The authors have declared no competing interest.

## References

1. R. E. Howes et al., The global distribution of the Duffy blood group. Nat Commun 2, 266 (2011).

2. P. C. Sabeti et al., Genome-wide detection and characterization of positive selection in human populations. Nature 449, 913–918 (2007).

3. D. Burgner, S. E. Jamieson, J. M. Blackwell, Genetic susceptibility to infectious diseases: big is beautiful, but will bigger be even better? The Lancet Infectious Diseases 6, 653–663 (2006).

4. S. J. Chapman, A. V. Hill, Human genetic susceptibility to infectious disease. Nat Rev Genet 13, 175–188 (2012).

5. L. Salvaudon, T. Giraud, J. A. Shykoff, Genetic diversity in natural populations: a fundamental component of plant-microbe interactions. Curr Opin Plant Biol 11, 135–143 (2008).

6. A. L. Laine, J. J. Burdon, P. N. Dodds, P. H. Thrall, Spatial variation in disease resistance: from molecules to metapopulations. J Ecol 99, 96–112 (2011).

7. B. P. Lazzaro, B. K. Sceurman, A. G. Clark, Genetic basis of natural variation in D. melanogaster antibacterial immunity. Science 303, 1873–1876 (2004).

8. M. C. Tinsley, S. Blanford, F. M. Jiggins, Genetic variation in Drosophila melanogaster pathogen susceptibility. Parasitology 132, 767–773 (2006).

9. D. J. Obbard, G. Dudas, The genetics of host-virus coevolution in invertebrates. Curr Opin Virol 8, 73–78 (2014).

10. E. M. Duxbury et al., Host-pathogen coevolution increases genetic variation in susceptibility to infection. Elife 8, (2019).

11. M. A. Duffy et al., Ecological context influences epidemic size and parasite-driven evolution. Science 335, 1636–1638 (2012).

12. R. M. Penczykowski, A. L. Laine, B. Koskella, Understanding the ecology and evolution of host-parasite interactions across scales. Evol Appl 9, 37–52 (2016).

13. P. H. Thrall et al., Rapid genetic change underpins antagonistic coevolution in a natural host-pathogen metapopulation. Ecol Lett 15, 425–435 (2012).

14. E. R. Westra et al., Parasite Exposure Drives Selective Evolution of Constitutive versus Inducible Defense. Curr Biol 25, 1043–1049 (2015).

15. M. Boots, The evolution of resistance to a parasite is determined by resources. Am Nat 178, 214–220 (2011).

16. D. M. Gwynn, A. Callaghan, J. Gorham, K. F. Walters, M. D. Fellowes, Resistance is costly: trade-offs between immunity, fecundity and survival in the pea aphid. Proc Biol Sci 272, 1803–1808 (2005).

17. R. Medzhitov, D. S. Schneider, M. P. Soares, Disease tolerance as a defense strategy. Science 335, 936–941 (2012).

18. A. R. Burmeister et al., Pleiotropy complicates a trade-off between phage resistance and antibiotic resistance. Proc Natl Acad Sci U S A 117, 11207–11216 (2020).

19. M. C. Rigby, R. F. Hechinger, L. Stevens, Why should parasite resistance be costly? Trends in Parasitology 18, 116–120 (2002).

20. B. R. Levin, V. Perrot, N. Walker, Compensatory mutations, antibiotic resistance and the population genetics of adaptive evolution in bacteria. Genetics 154, 985–997 (2000).

21. J. Bjorkman, I. Nagaev, O. G. Berg, D. Hughes, D. I. Andersson, Effects of environment on compensatory mutations to ameliorate costs of antibiotic resistance. Science 287, 1479–1482 (2000).

22. R. C. Allen, J. Engelstadter, S. Bonhoeffer, B. A. McDonald, A. R. Hall, Reversing resistance: different routes and common themes across pathogens. Proc Biol Sci 284, (2017).

23. A. R. Kraaijeveld, H. C. Godfray, Trade-off between parasitoid resistance and larval competitive ability in Drosophila melanogaster. Nature 389, 278–280 (1997).

24. M. D. Fellowes, A. R. Kraaijeveld, H. C. Godfray, Trade-off associated with selection for increased ability to resist parasitoid attack in Drosophila melanogaster. Proc Biol Sci 265, 1553–1558 (1998).

25. M. D. E. Fellowes, A. R. Kraaijeveld, H. C. J. Godfray, Association between Feeding Rate and Parasitoid Resistance in Drosophila Melanogaster. Evolution 53, 1302–1305 (1999).

26. B. Lemaitre, J. Hoffmann, The Host Defense of <i>Drosophila melanogaster</i>. Annual Review of Immunology 25, 697–743 (2007).

27. A. R. Kraaijeveld, H. C. J. Godfray, Geographic Patterns in the Evolution of Resistance and Virulence in <i>Drosophila</i> and Its Parasitoids. The American Naturalist 153, (1999).

28. S. Dupas, Y. Carton, M. Poirie, Genetic dimension of the coevolution of virulence-resistance in Drosophila -- parasitoid wasp relationships. Heredity (Edinb) 90, 84–89 (2003).

29. H. A. Orr, S. Irving, The Genetics of Adaptation: The Genetic Basis of Resistance to Wasp Parasitism in Drosophila Melanogaster. Evolution 51, 1877–1885 (1997).

30. Y. Carton, A. J. Nappi, Drosophila cellular immunity against parasitoids. Parasitology Today 13, 218–227 (1997).

31. M. Poirie et al., Drosophila resistance genes to parasitoids: chromosomal location and linkage analysis. Proc Biol Sci 267, 1417–1421 (2000).

32. M. Hita et al., Mapping candidate genes for Drosophila melanogaster resistance to the parasitoid wasp Leptopilina boulardi. Genet Res 88, 81–91 (2006).

33. K. M. Jalvingh, P. L. Chang, S. V. Nuzhdin, B. Wertheim, Genomic changes under rapid evolution: selection for parasitoid resistance. Proc Biol Sci 281, 20132303 (2014).

34. C. Kim-Jo, J. L. Gatti, M. Poirie, Drosophila Cellular Immunity Against Parasitoid Wasps: A Complex and Time-Dependent Process. Front Physiol 10, 603 (2019).

35. C. J. Evans, T. Liu, U. Banerjee, Drosophila hematopoiesis: Markers and methods for molecular genetic analysis. Methods 68, 242–251 (2014).

36. A. B. Leitão et al. (NERC EDS Environmental Information Data Centre, 2022).

37. B. Wertheim et al., Genome-wide gene expression in response to parasitoid attack in Drosophila. Genome Biol 6, R94 (2005).

38. E. S. Keebaugh, T. A. Schlenke, Adaptive evolution of a novel Drosophila lectin induced by parasitic wasp attack. Mol Biol Evol 29, 565–577 (2012).

39. T. A. Schlenke, J. Morales, S. Govind, A. G. Clark, Contrasting infection strategies in generalist and specialist wasp parasitoids of Drosophila melanogaster. PLoS Pathog 3, 1486–1501 (2007).

40. G. D. Brown, J. A. Willment, L. Whitehead, C-type lectins in immunity and homeostasis. Nat Rev Immunol 18, 374–389 (2018).

41. K. Natarajan, N. Dimasi, J. Wang, R. A. Mariuzza, D. H. Margulies, Structure and function of natural killer cell receptors: multiple molecular solutions to self, nonself discrimination. Annu Rev Immunol 20, 853–885 (2002).

42. W. Huang et al., Natural variation in genome architecture among 205 Drosophila melanogaster Genetic Reference Panel lines. Genome Research 24, (2014).

43. A. M. Early et al., Survey of Global Genetic Diversity Within the <i>Drosophila</i> Immune System. Genetics 205, (2017).

44. J. B. Lack, J. D. Lange, A. D. Tang, R. B. Corbett-Detig, J. E. Pool, A Thousand Fly Genomes: An Expanded <i>Drosophila</i> Genome Nexus. Molecular Biology and Evolution 33, (2016).

45. X. Yi et al., Sequencing of 50 Human Exomes Reveals Adaptation to High Altitude. Science 329, (2010).

46. F. Fleury et al., Ecological and genetic interactions in Drosophila-parasitoids communities: a case study with D. melanogaster, D. simulans and their common Leptopilina parasitoids in south-eastern France. Genetica 120, 181–194 (2004).

47. J. E. McGonigle et al., Parallel and costly changes to cellular immunity underlie the evolution of parasitoid resistance in three Drosophila species. PLoS Pathogens 13, (2017).

48. M. V. Olson, When less is more: gene loss as an engine of evolutionary change. Am J Hum Genet 64, 18–23 (1999).

49. J. G. Monroe, J. K. McKay, D. Weigel, P. J. Flood, The population genomics of adaptive loss of function. Heredity (Edinb) 126, 383–395 (2021).

50. Y. C. Xu, Y. L. Guo, Less Is More, Natural Loss-of-Function Mutation Is a Strategy for Adaptation. Plant Commun 1, 100103 (2020).

51. L. Salazar-Jaramillo et al., Inter- and intra-species variation in genome-wide gene expression of Drosophila in response to parasitoid wasp attack. BMC Genomics 18, 331 (2017).

52. T. F. Mackay et al., The Drosophila melanogaster Genetic Reference Panel. Nature 482, 173–178 (2012).

53. S. Dupas, M. Brehelin, F. Frey, Y. Carton, Immune suppressive virus-like particles in a <i>Drosophila</i> parasitoid: significance of their intraspecific morphological variations. Parasitology 113, (1996).

54. A. B. Leitão et al., Independent effects on cellular and humoral immune responses underlie genotype-by-genotype interactions between Drosophila and parasitoids. PLoS Pathogens 15, 1–13 (2019).

55. D. Bates, M. Mächler, B. Bolker, S. Walker, Fitting Linear Mixed-Effects Models Using <b>lme4</b>. Journal of Statistical Software 67, (2015).

56. T. Hothorn, F. Bretz, P. Westfall, Simultaneous Inference in General Parametric Models. Biometrical Journal 50, (2008).

57. M. J. Wade, R. G. Winther, A. F. Agrawal, C. J. Goodnight, Alternative definitions of epistasis: dependence and interaction. Trends in Ecology & Evolution 16, (2001).

58. J. Thurmond et al., FlyBase 2.0: the next generation. Nucleic Acids Research 47, D759–D765 (2019).

59. K. W. Broman, H. Wu, S. Sen, G. A. Churchill, R/qtl: QTL mapping in experimental crosses. Bioinformatics 19, 889–890-889–890 (2003).

60. J. Varaldi, D. Lepetit, Deciphering the behaviour manipulation imposed by a virus on its parasitoid host: insights from a dual transcriptomic approach. Parasitology 145, (2018).

61. S. J. Gratz et al., Highly Specific and Efficient CRISPR/Cas9-Catalyzed Homology-Directed Repair in <i>Drosophila</i>. Genetics 196, (2014).

62. J. Bischof, R. K. Maeda, M. Hediger, F. Karch, K. Basler, An optimized transgenesis system for Drosophila using germ-line-specific phiC31 integrases. Proc Natl Acad Sci U S A 104, 3312–3317 (2007).

63. D. Korona et al., Characterisation of protein isoforms encoded by the Drosophila Glycogen Synthase Kinase 3 gene shaggy. PLoS One 15, e0236679 (2020).

64. T. Tokusumi, D. A. Shoue, Y. Tokusumi, J. R. Stoller, R. A. Schulz, New hemocyte-specific enhancer-reporter transgenes for the analysis of hematopoiesis in Drosophila. Genesis 47, 771–774 (2009).

65. H. Li, Aligning sequence reads, clone sequences and assembly contigs with BWA-MEM. 00, 1–3 (2013).

66. T. Picard, No Title. Broad Institute, GitHub repository, (2019).

67. H. Li, R. Durbin, Fast and accurate short read alignment with Burrows-Wheeler transform. Bioinformatics 25, 1754–1760 (2009).

68. A. McKenna et al., The Genome Analysis Toolkit: A MapReduce framework for analyzing next-generation DNA sequencing data. Genome Research 20, (2010).

69. B. Longdon, J. D. Hadfield, C. L. Webster, D. J. Obbard, F. M. Jiggins, Host Phylogeny Determines Viral Persistence and Replication in Novel Hosts. PLoS Pathogens 7, (2011).

70. A. M. Bolger, M. Lohse, B. Usadel, Trimmomatic: a flexible trimmer for Illumina sequence data. Bioinformatics 30, (2014).

71. R. A. Hoskins et al., Sequence Finishing and Mapping of Drosophila melanogaster Heterochromatin. Science 316, (2007).

72. V. Bansal, A statistical method for the detection of variants from next-generation resequencing of DNA pools. Bioinformatics 26, (2010).

73. P. Cingolani et al., A program for annotating and predicting the effects of single nucleotide polymorphisms, SnpEff. Fly 6, (2012).

74. K. Ye, M. H. Schulz, Q. Long, R. Apweiler, Z. Ning, Pindel: a pattern growth approach to detect break points of large deletions and medium sized insertions from paired-end short reads. Bioinformatics 25, (2009).

75. J. T. Robinson et al., Integrative genomics viewer. Nature Biotechnology 29, (2011).

76. D. Lüdecke, sjmisc: Data and Variable Transformation Functions. Journal of Open Source Software 3, (2018).

77. J. K. Grenier et al., Global Diversity Lines–A Five-Continent Reference Panel of Sequenced <i>Drosophila melanogaster</i> Strains. G3&#58; Genes|Genomes|Genetics 5, (2015).

78. F. Madeira et al., The EMBL-EBI search and sequence analysis tools APIs in 2019. Nucleic Acids Research 47, (2019).

79. P. Danecek et al., The variant call format and VCFtools. Bioinformatics 27, (2011).

80. T. Jombart, adegenet: a R package for the multivariate analysis of genetic markers. Bioinformatics 24, (2008).

81. J. Goudet, hierfstat, a package for r to compute and test hierarchical F-statistics. Molecular Ecology Notes 5, (2005).

82. V. Obenchain et al., VariantAnnotation: a Bioconductor package for exploration and annotation of genetic variants. Bioinformatics 30, (2014).

83. H. Li et al., The Sequence Alignment/Map format and SAMtools. Bioinformatics 25, (2009).

84. S. Kumar, G. Stecher, M. Li, C. Knyaz, K. Tamura, MEGA X: Molecular Evolutionary Genetics Analysis across Computing Platforms. Molecular Biology and Evolution 35, (2018).

85. K. P. Schliep, phangorn: phylogenetic analysis in R. Bioinformatics 27, (2011).

86. E. Paradis, J. Claude, K. Strimmer, APE: Analyses of Phylogenetics and Evolution in R language. Bioinformatics 20, (2004).

87. Z. Zhang, S. Schwartz, L. Wagner, W. Miller, A Greedy Algorithm for Aligning DNA Sequences. Journal of Computational Biology 7, (2000).

88. A.-S. Fiston-Lavier, N. D. Singh, M. Lipatov, D. A. Petrov, Drosophila melanogaster recombination rate calculator. Gene 463, (2010).

