## Supplementary material for "Recurrent loss of an immunity gene that protects *Drosophila* against a major natural parasite"

### Materials and Methods

*Drosophila melanogaster stocks maintenance*

We used the DGRP panel, which is a set of genome-sequenced inbred lines derived from a natural population in Raleigh, North Carolina, USA (*42, 52*). Based on preliminary data we selected two lines, DGRP-437 (Bloomington Drosophila Stock Center, BDSC#25194) and DGRP-892 (BDSC#28258) to serve as our resistant and susceptible lines in this study. Fly lines were maintained in glass vials on a cornmeal diet (per 1200ml water: 13g agar, 105g dextrose, 105g maize, 23g yeast, 35ml Nipagin 10% w/v). The vials were kept in a 25°C room with 70% of relative humidity and a light-dark cycle of 14 hour/10 hour. After a maximum of 20 days, adult flies were transferred into new clean vials.

*Leptopilina boulardi stock maintenance*

*Leptopilina boulardi* is an endoparasitoid wasp (Hymenoptera; Figitidae) infecting the larval stages of *Drosophila* (*27*). We used the low virulence *L. boulardi* strain G486 (Gif/Yvette, stock number 486) (*53*) created from an isofemale strain selected from a population of Brazzaville, Congo. We maintained wasps in the laboratory on an outcrossed population of *D. melanogaster* collected from Cambridge. To do so, *D. melanogaster* eggs laid overnight on agar plates in population cages were collected using phosphate buffed saline (PBS) and a bristle brush to ensure all eggs were removed from the agar plate and they were transferred into 15ml falcon tubes. Using a P1000 Gilson pipette with the end of the tips cut off, 500ul of the eggs were transferred into a 1.5ml microcentrifuge tube. Subsequently, a P20 Gilson pipette with the ends of the tip cut was used to transfer 6µl of eggs into plastic cornmeal food vials. After sex identification using the antennae size, 2-3 females and 1 male *L. boulardi* wasps were added to each vial. The vials were kept in a 25°C room with a relative humidity of 70% and a light-dark cycle of 14 hour/10 hour for at least 24 days. Once the adult wasps emerged, we administered $CO_2$ to anaesthetize them and put them into plastic cornmeal food vials supplemented with honey and kept at room temperature.

*Scoring encapsulation rates in larvae – resistance assay*

The female wasp lays its eggs inside the Drosophila larva's hemocoel through the ovipositor. If wasp larvae survive *Drosophila* immune defense, they will feed on its tissue and emerge from the pupae of the fly. If the immune response is successful, flies encapsulate wasp eggs, and this results in black melanized capsules easily visualized under a microscope.

We had used two different assay strategies in 2018 and 2019, which were largely similar except in how they controlled *D. melanogaster* larval density. For the resistance assays performed in 2018, 100-200 flies were maintained in population cages with apple agar media on 90 mm plates (per plate: 0.565g agar, 0.625g glucose, 6.25ml apple juice, 18.75ml water, 0.375ml Nipagin 10% w/v) covered with yeast (Sigma-Aldrich: YSC2) mixed in water. The plates were changed every morning and evening. Plates from the first day after egg transfer were discarded. Plates containing eggs laid overnight from the following three days were retained and we performed larval transfer. Eggs on the surface of the 90mm agar plates were washed with PBS. This solution was poured into a 15ml falcon tube, and the eggs were allowed to settle to the bottom. Using a cut P1000 tip, 500µL of solution saturated with eggs was transferred into a 1.5µL microcentrifuge tube. Then, five 30mm plates containing cornmeal were scratched in the middle. A cut P200 tip was used to transfer 13µL of solution from the microcentrifuge tube containing eggs to each 30mm cornmeal plate, in a line around the edge of the scratched area. The 30mm cornmeal plates were incubated at 25°C with a relative humidity of 70% and a light-dark cycle of 14 hour/10 hour. Two days after incubation, larvae were transferred from the plates into plastic vials containing cornmeal media using a pair of forceps under a Leica MZ6 stereomicroscope. From each plate, we attempted to make four vials containing 40 larvae each. We added three female wasps using a paintbrush to each vial after administering $CO_2$ to anaesthetize them. We incubated vials at 25°C for three hours to allow the wasps to infect the larvae and removed the wasps after that. Vials were kept at 25°C for another two days to allow them to develop.

We simplified the resistance assays in 2019 as we found we were able to attain a similar egg density by pipetting 5ul of eggs in PBS suspension from the 1.5µL microcentrifuge tube. Rather than using larger population cages, we placed ~50 flies in inverted 100ml laboratory plastic beakers with the opening covered with 30mm apple agar plates with yeast using the same recipe as above. Eggs from these plates collected into microcentrifuge tubes as above. We transferred 5ul of egg suspension per stock directly into plastic cornmeal food vials without explicitly controlling for number of larvae. The remaining steps were similar to the 2018 assays.

We dissected developed fly larvae to identify whether the wasp had survived or been encapsulated. To do so, we added 15% (w/v) sugar solution in $ddH_20$ and dissolved the cornmeal allowing the larvae to float to the top. We transferred individual larvae onto separate drops of $ddH_20$ on a clear plastic lid and dissected them using two pairs of forceps. We removed the cuticle of the larvae to check and allow for wasp larvae, if any, to settle to the bottom of the droplet of water. We scored flies as having encapsulated the wasp if the wasp larvae within the fly were all melanized and/or immotile. We score fly larvae as having failed to encapsulate if there is at least one non-melanized and motile wasp larvae in its body. We excluded fly larvae without any wasp as these were uninfected samples. We dissected 20 infected larvae from each vial. We used gloves during dissection and continuously cleaned the forceps during the dissection with ethanol.

*Characterizing melanization of resistant and susceptible parental lines*

We tested for difference in encapsulation rates between DGRP-437 and DGRP-892 using the method described above and analyzed the data with a generalized linear mixed model with a logit link function using the equation below:

$$\text{glmer}(\text{Encapsulation ratio} \sim \text{DGRP line} + (1|\text{Replica}), \text{family} = \text{"binomial"})$$

We also characterized the melanization response in the resistant and susceptible DGRP lines used for genetic mapping by injecting them with mineral oil containing homogenized parasitoid wasp. Twenty male *L. boulardi* and *A. tabida* were homogenized in 200µl of paraffin oil (Sigma-Aldrich M5904) using a pestle and injected into fly larvae using the protocol established in (*54*). Injected larvae were moved using forceps to cornmeal food vials where each vial contained 40 larvae. Vials were incubated in the 25°C room with a 14hr light-10hr dark cycle and 70% humidity for 10 – 48 hours depending on the scheduled dissection time. Prior to dissection, 15% w/v sugar solution was used to remove the larvae from food. The larvae were dissected on a PBS droplet that caused the injected oil droplets to float to the surface and the oil droplets were classed by their melanization state. The following binomial generalized linear model was built:

$$\text{glm}(\text{Encapsulation ratio} \sim \text{DGRP line} + \text{Time post injection}, \text{family} = \text{"binomial"})$$

The Wald $\chi^2$ statistic was used to assess significance of the fixed effects. A likelihood ratio test comparing the above model to a more complicated model containing an interaction term between line and time post injection, dissection date, injection date, dissection time and injection time was not significant, so the simpler model was used.

Wasp embryo development in the resistant and susceptible DGRP lines was observed at 18, 24 and 26 hours post *L. boulardi* infection. After performing the 2019 larval resistance assay protocol, the wasp embryo was isolated on a microscope slide and the *D. melanogaster* larval tissue was removed. A cover slide was placed on top of the reaction well and it was transferred into a humidity chamber. The images of the wasp embryo were taken on a Leica Microscope with an 8x1 objective.

A fly stock expressing the fluorescent reporters in hemocytes, $w^{1118}$; *Hml*Δ-GAL4 UAS-GFP; *Msn*-mCherry, was used to visualize hemocytes in resistance and susceptible DGRP lines. Female flies from the stock were crossed to male DGRP-437 and DGRP-892. The 2019 larval resistance assay protocol was used, and we assayed encapsulation in the $F_1$ progeny following infection with parasitoid wasp or injection of an oil droplet containing homogenized *L. boulardi*. Fisher's exact tests were used the test if the number of samples that melanized the wasp embryo or the oil differed. 24 hours post a separate set of infections and injections, hemocytes surrounding the wasp embryo were observed using a Leica DFC350 FX microscope and the Leica Applications Suite software (Leica Microsystems 2016).

*Dominance of resistance genotype*

We assessed the dominance of the resistance phenotype by crossing DGRP-437 with DGRP-892. We performed the crosses in two directions: virgin females of DGRP-437 were crossed to DGRP-892 males in one cross and vice-versa for the reciprocal cross. We used ~50 males and ~100 females for each cross and placed them in 135mm by 100mm (length by width) population cages. We tested for resistance to *L. boulardi* in the heterozygote progeny resulting from these crosses at the larval stage. We followed the 2018 protocol for the resistance assay and distinguished male and female larvae by the size of the developing gonads, seen as translucid region in posterior part of the fat body. We tested if parental genotypes (cross) and sex of progeny influenced encapsulation in the $F_1$s using binomial generalized linear mixed-effects model in the lme4 v1.1-23 R package (*55*). Wasp encapsulation was included as the binomial response variable. The following model was fit:

$$\text{glmer}(\text{Encapsulation} \sim \text{Cross} + \text{Sex} + (1|\text{Replica}), \text{family} = \text{"binomial"})$$

We performed Tukey's multiple comparison test on the fitted model using *glht* function from the multcomp v1.4-14 R package (*56*) and adjusted the resulting *p*-values using the Holm approach to compare the encapsulation levels in reciprocal crosses post-hoc. A likelihood ratio test was used to investigate if sex impacted encapsulation.

*Chromosome substitution lines assay*

We performed standard genetic crosses using DGRP-437 and DGRP-892 to generate lines with unique combinations of chromosomes from the susceptible and resistant parents to identify the chromosome(s) associated with encapsulation. We set up crosses to create six substitution lines carrying varying combinations of the X, II and III chromosomes from DGRP-437 and DGRP-892. Double balanced chromosome stocks used for the crosses, *f*/FM7 and $w^*$; *If*/CyO; MKRS/TM6B, were obtained from the Fly facility in the Department of Genetics, University of Cambridge.

To confirm the crosses were successful we genotyped the flies using EcoR1 cut sites primers designed by (*52*) to genotype the X, II and III chromosomes in the substitution lines. Primers X2, 2L1 and 3L3 were polymorphic between DGRP-437 and DGRP-892. After DNA extraction and PCR, we performed the EcoRI restriction digest. For the digest, we used 0.6µL EcoR1-HF (NEB: R3101), 3µL 10x NEBBuffer 2.1 (NEB: B7202S), 16.4µL nuclease-free water (Ambion: AM9930) and 10µL PCR product per sample. We performed the digest using a Veriti™ 96-Well Thermal Cycler with the following protocol: 1) 37°C for 1 hour and 30 minutes, 2) 65°C for 20 minutes. We ran the undigested and digested PCR products on a 2% agarose (Bioline: BIO-41025) gel in TBE with 4µL EtBr solution (Sigma-Aldrich: E1510). We used 6X loading dye (Termo Scientific: R0611) for loading the samples running it alongside Hyperladder 1kbp (Bioline: BIO-33053). The samples were run on a BioRad PowerPac™ HC submerged in TBE at 120mV for about 50 minutes. Then we took the image of the gel with a BioRad Gel Doc™ XR+ to score the genotype of each sample.

We assayed resistance levels in the larvae for chromosome substitution lines and the parental DGRP lines using a mixture of the 2018 and 2019 approaches described above (see section: resistance assay). We tested for differences in resistance by fitting a generalized binomial (logistic) linear mixed model by maximum likelihood in the lme4 v1.1-23 R package (*55*). In our model, wasp encapsulation was included as the binomial response variable. The chromosome substitution line was included as a fixed effect. We included the assay vial ('replica') and date of dissection date as a random effect to allow for overdispersion. The latter factor would account for the differences in assay methodology in 2018 and 2019. The fitted model was:

$$\text{glmer}(\text{Encapsulation} \sim \text{Line} + (1|\text{Replica}) + (1|\text{Dissection date}), \text{family} = \text{"binomial"})$$

Further, we performed the Tukey's multiple comparison test on the fitted model using *glht* function from the multcomp v1.4-14 R package (*56*) and adjusted the resulting *P*-values using the Holm approach to identify the significant comparisons. We also tested for epistasis between chromosomes II and III using a multiplicative model (*57*). We fitted a glmer including the two autosomes and their interaction as fixed effects using the formula:

$$\mathrm{glmer}(\mathrm{cbind}(\#\mathrm{resistant}, \#\mathrm{susceptible}) \sim \mathrm{chrII} * \mathrm{chrIII} + (1|\mathrm{Replica}), \mathrm{family} = \text{"binomial"})$$

We performed an analysis of variances employing $\chi^2$ statistic to assess the impact of the chromosomes and their interaction on encapsulation ratios.

*DNA extraction using Chelex resin*

We collected single flies in strip tubes, froze them at -18°C and performed DNA extractions subsequently. First, we prepared a 5% (w/v) Chelex solution (Sigma Aldrich: C7901) in $ddH_20$ and placed a magnetic bead within it. We autoclaved the mixture. We stirred the prepared Chelex solution on a magnetic stirrer (StarLab: N2400-3010) at 700rpm and added 150μL of solution using a cut P200 tip to each PCR tube as the solution was being stirred. Further, we added ~ six Zirconia 1.0 mm diameter beads (BioSpec: 11079110z) to each PCR tube. We then homogenized the flies using the TissueLyser II (Qiagen: 85300) for two minutes at 25Hz. We repeated the homogenization and checked that the flies were lysed at the end. We spun the tubes using a Beckman Coulter Optima™ MAX-XP tabletop ultracentrifuge at 1,000g for two minutes. Then, we added 1.5μL of proteinase K (Sigma Aldrich: P8044) at 20mg/mL in nuclease-free water to each sample and incubated the tubes at 56°C overnight. We spun the tubes again using the ultracentrifuge at 1,000 g for two minutes. We transferred ~100μL of supernatant into a new PCR plate taking care not to pipette any of the Chelex beads. We incubated the PCR plates at 94°C for 15 minutes to deactivate the proteinase K.

*Primer design and PCR for genotyping*

SNPs and indels that vary between DGRP-437 and DGRP-892 were used to genotype progeny resulting from crosses using the two parental DGRP stocks. We attained variant positions from the DGRP Freeze 2.0 website (http://dgrp2.gnets.ncsu.edu, last accessed May 2019). The DGRP Freeze 2.0 was mapped to version 5 of the genome but the recombination and physical maps from Flybase (FB2019_02) (*58*) are aligned to the version 6. Therefore, we converted the coordinates from version 6 to 5 using the *Coordinator Converter* function in Flybase. We searched for SNPs or indels 40 – 200bp long in each interval of interest. We then attained the sequence 1,000bp on either side of that indel. We used Primer3 (http://primer3.ut.ee, last accessed May 2019) to design primers to amplify the variant. We opted for products of sizes 300-1000 and only selected those that amplified the variant of interest.

In most cases, we performed PCR by mixing 0.5μL of 10μM dNTP (NEB: N0447), 2.5μL 10x ThermoPol® Reaction Buffer (NEB: B9004S), Taq DNA Polymerase (NEB: M0267X), 0.5μL of 10μM of forward primer, 0.5μL of 10μM of reverse primer, 18.875μL nuclease-free water (Ambion: AM9930) and 2 μL genomic DNA per sample. We performed the touchdown PCR reaction on the Veriti™ 96-Well Thermal Cycler with the following protocol: 1) 95°C for two minutes, 2) 10 cycles touchdown of 95°C for 30 seconds, 62°C to 52°C for 30 seconds where the temperature drops by 1°C every cycle, 68°C for seconds and 3) 25 cycles of 95°C for 30 seconds, 52°C for 15 seconds, 68°C for 30 seconds.

*QTL mapping of phenotyped larvae*

We performed QTL mapping to identify the genetic regions on the chromosome II associated with resistance to infection by the parasitic wasp. We crossed 100-200 female $X^{892}$; $II^{892}$; $III^{437}$ and 100-200 male $X^{892}$; $II^{437}$; $III^{437}$ to create $F_1$ individuals heterozygous for the second chromosome ($X^{892}$; $II^{437/892}$; $III^{437}$). We backcrossed 100-200 $F_1$ heterozygote males to 100-200 female $X^{892}$; $II^{892}$; $III^{437}$ in a single 210mm by 100mm population cage to generate our recombinant mapping population. We then parasitized larvae in this mapping population following the 2018 resistance assay protocol. Once an infected larva was dissected and scored, we placed the dissected fly larvae into PCR tubes kept on ice taking care not to sample any melanized capsule or wasp larvae. Once we collected 24 samples, we moved the PCR tubes to a -18°C freezer. Then, we extracted the DNA following the Chelex DNA extraction protocol described above and kept them at -18°C. We designed primers to target indel polymorphisms in DGRP lines 437 and 892 to genotype multiple positions within chromosome II (Table S3). We ran the PCR products on a 1.2% agarose gel with EtBr for 45-60 minutes until the bands were well resolved. We took image of the gels with a BioRad Gel Doc™ XR+ to score the genotype of each sample varying exposure time between 3 and 9 seconds.

We performed QTL mapping using the standard genetic map of *D. melanogaster* from Flybase. We only used samples that were fully typed for all ten markers. We used the R/QTL v1.46-2 R package (*59*) for QTL mapping. We first calculated genotyped probabilities between markers, assuming a genotyping error probability of 0.05 and step size of 1 cM using the Haley-Knott method, and then used linear regression to test whether these probabilities were associated with the phenotype. We also performed composite interval mapping using a window size of 10. We used the LOD drop of 1.5 from the maximal LOD score to establish the interval on the genetic map that is likely to contain the gene affecting the phenotype.

*High resolution QTL mapping with informative recombinants*

To increase the resolution of our QTL, we performed additional genotyping of adults that encapsulated the wasp within the identified interval of interest. We performed the same cross as before in cages with the goal of generating large numbers of flies. We set up the cross three times in separate 210mm by 100mm population cages. For the first cage, we followed the same approach as was done to score resistant and susceptible larvae in 2018, which included both egg and larval transfer (see section: resistance assay). In the second cage, we had 50-100 female $X^{892}$; $II^{892}$; $III^{437}$ and 50-100 male $X^{892}$; $II^{437}$; $III^{437}$ and in the third one we had 100-200 female $X^{892}$; $II^{892}$; $III^{437}$ and 100-200 male $X^{892}$; $II^{437}$; $III^{437}$. For the second and third cages, we collected eggs laid over a period of two weeks after each cage was first set up. We transferred 5µL eggs in PBS solution, collected from 90mm agar plates covered in yeast after cage set-up, directly to cornmeal plastic vials without yeast. In addition to using plates from the morning after an overnight egg lay, we transferred eggs collected on plates from the evening into plastic vials. On some dates, eggs lay occurred over a 24-hour period. For most vials, 2-4 days after egg transfer, we transferred one female wasp and placed it in the vials for 1-3 days. Vials were kept at 25°C with a relative humidity of 70% and a light-dark cycle of 14hr-10hr. We waited for 13 days post egg transfer for all flies to emerge. We checked for flies with melanized capsules after administering $CO_2$ to anaesthetize them. We discarded flies without any capsules from the infected vials. We collected ~1,900 flies with melanized capsules from infected vials and ~500 control flies from uninfected vials. Flies were collected in PCR strip tubes and frozen at -18°C.

As uninfected adults occasionally produced melanotic nodules that can be mistaken by melanotic capsules, we tested these flies for wasp DNA using quantitative PCR (qPCR). We designed primers from a randomly chosen location of the *L. boulardi* genome (scaffold_2167) obtained from NCBI (ASM312160v1) (*60*). The forward primer was AGCAGCGATTGAAACAGTTGT starting at 16,463bp and the reverse primer was TGATTGTTGAACACGTCGGA starting at 16,679bp. Alongside melanized *D. melanogaster* samples, we performed qPCR on DNA extracted from *L. boulardi* strain G468 with Chelex beads and uninfected *D. melanogaster* flies. For each reaction, we added 5µL SensiFAST SYBR® Hi-ROX mix (Bioline: BIO-92020), 2.75µL nuclease-free water, 2µL of DNA and 0.125 µL of each of 10 µM forward and 10 µM reverse primers. We performed the reactions with MicroAmp™ Fast Optical 96-Well Reaction Plate with Barcode (Applied Biosystems: 4346906) and covered the plates with MicroAmp™ Optical Adhesive Film (Applied Biosystems: 4311971). We performed qPCR using the StepOnePlus™ Real-Time PCR System v2.2.3 (Applied Biosystems: 4376600) with the following protocol: 1) 95°C for two minutes, 2) 40 cycles of 95°C for five seconds and 60 °C for 30 seconds. Melt curves were obtained immediately following the step and hold protocol: multiple cycles of 15 seconds at 95°C and 60°C to 95°C for one minute where the temperature increased by 0.3°C every cycle. We analyzed melt curves using the StepOne™ v2.3 software. Wasp DNA samples amplified had a peak melting temperature close to 77-78°C whereas DNA from uninfected *D. melanogaster* samples did not amplify (Figure S9). Sample DNA were extracted with Chelex resin using the approach detailed above (see section: DNA extraction using Chelex resin). Wasp infection status in the $F_2$s were determined by comparing the melt curves to those obtained for the wasp DNA samples. We only retained samples that amplified wasp DNA for our analysis.

We extracted DNA from the flies that had encapsulated the wasp and uninfected flies and genotyped them using the 3 cM and 27 cM indel primers (Table S3) based on the interval attained from QTL mapping of resistant and susceptible larvae. Flies that were not typed at both markers were excluded. Following that, we selected flies that had encapsulated the wasp and had a breakpoint between 3 cM and 27 cM. For these flies, we also genotyped the indel markers 7 cM, 10.3 cM, 12 cM and 17 cM. For markers with missing values, we imputed genotypes if the flanking genotypes were the same.

We did additional Sanger sequencing using SNP based primers (Table S3) on samples with breakpoints in the interval of interest. In this case, we only performed Sanger sequencing for individuals where the flanking indel markers indicated there was a recombination breakpoint. We first did PCR as detailed previously (see section: Primer design and PCR for genotyping). Then, 10µL of primer product was cleaned using 1µL Shrimp Alkaline Phosphatase (NEB:

M0371), 0.1µL Exonuclease I (NEB: M0293) and 3.9µL of nuclease-free water at 37°C for 1 hour and then the enzymes were inactivated at 80°C for 15 minutes. We then performed sequencing using the BigDye® Terminator v3.1 cycle sequencing kit (Applied Biosystems: 4337455). For each reaction, we added 1µL of BigDye® Terminator v3.1 Ready Reaction Mix, 2µL of 5X sequencing buffer, 2µL of cleaned PCR product, 4µL of nuclease-free water and 1µL of 3.3µM forward primer. We performed the sequencing reaction on the Veriti™ 96-Well Thermal Cycler with the following protocol: 1) 95°C for one minute and 2) 25 cycles touchdown of 95°C for 30 seconds, 50°C for 20 seconds and 60°C for four minutes. We sent the samples to Source Bioscience (https://www.sourcebioscience.com/, last accessed May 2019) for sequencing clean-up and Sanger sequencing. The sequenced chromatograms were analyzed using Geneious software v 11.0.3. Samples sequenced using the same primer were aligned. Low quality samples where multiple base calls were overlapping across the entire sequence were excluded. Heterozygous SNP variants were called when there were two clear base peaks at a given position. Following Sanger sequencing, we imputed missing genotypes based on flanking markers with the help of the zoo v1.8-8 R package. A $\chi^2$ drop to identify informative markers was selected was by simulating 1000 datasets based on the observed risk ratio estimated from non-recombinant flies and the observed recombination fraction and identifying the $\chi^2$ drop that defined a region that included the gene in 95% of simulations.

*CRISPR Cas9-mediated mutagenesis of Lectin-24A*

The necessity of *Lectin-24A* for resistance to parasitoid wasps was tested by using CRISPR Cas9-mediated mutagenesis to generate lines with mutations or deletions in *Lectin-24A* in the resistant background. Guide RNA (gRNA) constructs were designed to encode two separate gRNAs in duplicate using pCFD5-w as the backbone plasmid (pCFD5_w was a gift from Michael Boutros (Addgene plasmid # 112645); http://n2t.net/addgene:112645; RRID:Addgene_112645). gRNA for *Lectin-24A* were identified using the Target Finder (*61*) against dm6 using high stringency and only allowing NGG PAM sites. Two sequences on chromosome 2L were selected both with 0 predicted off-target effects. The first sequence was between 3717384 and 3717406 (GGTCTCCA|AAGATTCATGCATGG) and the second sequence was between 3717584 and 3717606 (CATACTCC|ATAACTGGCTTCAGG). The two gRNA were separated by 178bp and occurred close to the start of *Lectin-24A* coding region. Three fragments comprising tRNA, gRNA core and gRNA sequence were amplified using 10ng of pCFD5 as template in PCR reactions using the Lectin24a_PCR1,2,3 forward and reverse primers (using the protocol detailed in http://www.crisprflydesign.org/grna-expression-vectors/ and Table S4) and Q5 2xMaster Mix polymerase (NEB # M0492S). 600ng of pCFD5 was digested with BbsI restriction enzyme (NEB # R0539S). PCR amplified fragments and BbsI-digested pCFD5 vector were gel purified, quantified using Qubit HS DNA kit (Thermofisher # Q32854) and assembled using HiFi assembly master mix (NEB # E2621S) at 50°C for 1 hour with a 1:2:2:2 molar ratio of vector to the three fragments being cloned. Colonies were screened for correct sized (1.2 kbp) inserts using PCR with primers U63seqfwd and pCFDseqrev (Table S4), positive plasmids were purified and sequenced using the same primers. DNA for micro-injection of embryos was prepared using a plasmid midiprep kit (Qiagen # 12941).

To generate *Lectin-24A* mutants we first generated a transgenic fly line expressing the two gRNAs. The pCFD5-w LectinKO gRNA construct was microinjected into *D. melanogaster* line attP2Ar5 expressing a phiC31 integrase under the control of *vasa* promotor on chromosome 4 and harbouring an attP 2Ar5 landing site on the X chromosome (BDSC#24480) (*62*) by Bestgene Inc, USA. Mutations of *Lectin-24A* were obtained by expressing Cas9 controlled by the *vasa* promotor (BDSC#51323). Transformants were used in the crossing scheme outlined in Figure S10 to generate germline mutations in *Lectin-24A*. Individual males were screened by PCR for large deletions or insertions using primers that amplified the open reading frame of *Lectin-24A*. Those products not exhibiting noticeable changes in PCR product length were sanger sequenced for mutations at the site of each gRNA. Through this, we obtained a mutant with a 4bp insertion at 129bp after the start codon (TTAT) and a 2bp deletion and a 1bp insertion at 328bp after the start codon resulting in 97 truncated amino acid protein (Figure S11) that we named as *Lectin-24A*$^{\Delta 129}$. For the mutant and wild-type DGRP lines, larval encapsulation was scored using the methods above (see section: *Scoring encapsulation rates in larvae - resistance assay*) without controlling for larval density (the mutant lines produced very few larvae so there was little chance for overcrowding). 50-100 individuals were directly placed into a cornmeal vial for overnight egg lay. They adults were removed the next morning and larvae were infected with 2-3 parasitoid wasps at the 2/3 instar stage.

*Lectin-24A expression in DL2 cells*

In order to express Lectin-24A in *Drosophila* DL2 cells, it was cloned into the expression vector pMT-puro (gift from David Sabatini; Addgene plasmid #17923; http://n2t.net/addgene:17923; RRID:Addgene_17923) as a Lectin-24A-Flag-strepII tag fusion. The open reading frame of Lectin-24A was amplified from DGRP-437 genomic DNA using primers GGATCTAGATCGGGGTACACCACCATGTTTAGATTGTCAGTCTTAGTTCTGAACTTACTC and CTTTGTAGTCGATGCCATACTGGCATATGAATCTTTTTTCATAAG that added a consensus kozak sequence and removed the stop codon. The Flag-StrepII tag was amplified from a plasmid containing the 3xFLAG-StrepTagII-mVenus-StrepTagII cassette (plasmid provided by D. Korona) (*63*) using primers TATGGCATCGACTACAAAGACCATGACGGTGATTATAAAGAT and CTGATCAGCGGGTTTTTATTTTTCGAACTGCGGGTGGC. The expression vector pMT-puro was digested with the KpnI and PmeI restriction enzymes to remove the V5 and His-tags. The vector was then used in a HiFi assembly reaction with the two amplified fragments to form pMT-puro-Lectin-24A-Flag-Strep. pMT-puro-Lectin24-A-Flag-Strep plasmid was purified from *E.coli* using a Qiagen miniprep kit (Qiagen: 27106) and used to transfect *Drosophila* DL2 cells in a final volume of 2.2ml media using Effectene transfection reagent (Qiagen 301425) according to the manufacturer's instructions. 48 hours after transfection, protein expression was induced by adding $CuSO_4$ to a final concentration of 1mM. After incubating for 24 hours, cells were removed by centrifugation and StrepII-tagged Lectin-24A was precipitated from the media by incubating with 20ul washed Strep-Tactin XT beads (IBA life Sciences 2-4090-010) for 30 minutes. Beads were collected using a magnetic rack, then washed twice in 1ml cold PBS supplemented with protease inhibitor before they were boiled in sample buffer +/- 1:100 beta-mercaptoethanol.

*Visualising Lectin-24A localization*

The transgenic lines *Hml*Δ-GAL4 UAS-GFP (BDSC#30140) using the pan-hemocyte maker *Hml* was used to visualize hemocyte localization. *Lectin-24A* fused with C-terminal *Msn*-mCherry (*64*) was used to visualize its expression in *Drosophila* larvae. To generate *Msn*-mCherry tagged *Lectin-24A* constructs, pCFD5 was digested with XbaI and BglII to remove the dU6-3 promoter and gRNA scaffolds and the backbone vector was gel purified. A 3.4kbp genomic fragment encoding the open reading frame and a 2.5kbp *Lectin-24A* promoter was amplified using Q5 polymerase 2x master mix (NEB # M0492S) with a touch down 62-52 PCR program: 95$^{o}$C for 2 minutes followed by 10 cycles of 95$^{o}$C for 15s, 62$^{o}$C for 30s ramped down by 1$^{o}$C every cycle, 72$^{o}$C for 3 minutes followed by 25 cycles of 95$^{o}$C for 15s, 52$^{o}$C for 30s, 72$^{o}$C for 3 minutes. Primers CCATTTAGCCGATCAATTGACGAGTACTTCTACGCCGC and CTTTGTAGTCGATGCCATACTGGCATATGAATC that incorporated sequences complementary to the vector and mCherry at the 5' and 3' end of the fragment, respectively, were used to allow for Gibson assembly. The template in the PCR was genomic DNA isolated from either the DGRP-437 or DGRP-892 lines. Similarly, mCherry with a FLAG-tag was amplified using primers GTATGGCATCGACTACAAAGACGATGACGACAAG and TTTAACTGGCTACTCGTCCATGCCGCC. The 3'UTR of *Lectin-24A* was amplified using primers GGACGAGTAGCCAGTTAAAAACCAAAAAAAATC and CTGTTGCCGAGCACAATTGTCATTTCTTCATTTCGCCAC. Amplified fragments were gel purified, quantified using Qubit HS DNA assay kit (Thermofisher # Q32854) and used in a HiFi assembly reaction (NEB # E5520) with digested pCFD5 using an equimolar ratio of vector to fragments with a 1-hour incubation at 50$^{o}$C. Plasmids were sequenced with internal primers used to amplify the fragments. The vermillion marker was then excised by digesting plasmids with HindIII and replaced with the mini-white gene purified by digesting plasmid pWallium20 backbone plasmid (https://fgr.hms.harvard.edu/publications/transgenic-rnai-project-harvard-medical-school-resources-and-validation) with HindIII. The construct was injected into *D. melanogaster* line attP2Ar5 by the Fly Facility (Department of Genetics, University of Cambridge, UK) and transformants were selected by eye colour.

We extracted the mCherry-tagged Lectin-24A to ensure that it remained intact following the fusion. To do this, early third instar larvae from transgenic lines containing DGRP-437 Lectin-FLAG-mCherry or DGRP-892 Lectin-24A-FLAG-mCherry were infected with *L. boulardi* G486 or left uninfected for 24 hours. Larvae were harvested, washed in PBS, dried on filter paper, and homogenised in 20ul of lysis buffer containing 1% IGEPAL, 10mM Tris pH 7.5, 150mM NaCl supplemented 1 tablet to 50ml with protease inhibitor (Roche: 04 693 159 001) by grinding with a pestle. Samples were centrifuged at 17000g at 4$^{o}$C for 5 minutes to pellet debris and the supernatant was removed to a fresh tube. Protein was quantified using Bradford Assay. 20ug was added to 4ul of 4xSDS loading buffer supplemented with 1:25 beta-mercaptoethanol.

*Western blotting*

Presence of intact Lectin-24A protein, mCherry tagged Lectin-24A, and oligomerization was verified through western blotting. Samples were separated by SDS-PAGE using a NuPAGE 4-12% Bis-Tris Gel (Thermo Fisher: NP0322) including a lane containing PageRuler plus pre-stained protein ladder (Thermofisher: 26619). Protein was transferred to Amersham Protran

0.45 um Nitrocellulose blotting membrane (GE Healthcare: 10600008) and blocked in TBST with 5% milk for 30 minutes. Anti-FLAG M2 monoclonal antibody (Sigma: F1804) was diluted 1:5000 in 5% milk in TBST and the membrane was incubated with this over-night at $4^{o}$C. After washing, the membrane was incubated with IRDye 800CW Goat anti-mouse secondary (LiCor: 326 32210) (1:5000 dilution in TBST with 5% milk). To determine loading quantities or larval protein samples membranes were stripped and re-probed with anti-a-tubulin primary antibody (Sigma: T6199) diluted 1:10000 in 5% milk, washed and incubated with IRDye 800CW Goat anti-mouse secondary. Membranes were detected using Li-Cor Odyssey XF Imager for 10 minutes detection time for the 800nm wavelength and 2 minutes for the 700nm wavelength.

*Confocal microscopy imaging of wasp egg with tagged Lectin-24A*

We dissected wasp eggs from heterozygous $F_1$ larvae of the cross between *Lectin-24A-mCherry* females and *HmlΔ-GAL4 UAS-GFP* males at around 23 hours post-infection and fixed them in 4% formaldehyde directly on a microscope slide. The eggs were washed with PBS for three times, stained with DAPI for 5 minutes, then mounted in 50% glycerol (Sigma: G5516) with 0.35% n-Propyl gallate (Sigma: 02370) diluted in PBS and covered with a cover slip. The wasp eggs were then imaged using a Leica SP8 confocal microscope with 20x objective.

*Lectin-24A promoter indel haplotype characterization*

We Sanger sequenced the upstream and coding regions of *Lectin-24A* in 171 DGRP lines as indel variants in *Lectin-24A* were not identified in the genome sequences from DGRP Freeze 2.0. We amplified a 603bp fragment (dm6:2L:3717682-3718285) using the GGCGCCTCCTTCCACTATTT (forward) and TGGCTAACAAGGAGTAAGTTCAGA (reverse) primers. *Lectin-24A* coding sequence and neighboring regions were subject to Sanger sequencing as detailed above (see section: Fine mapping with resistant recombinant adults) in 171 DGRP lines. The primers were designed by Primer3 (http://primer3.ut.ee, last accessed July 2019) using the upstream and coding sequence of *Lectin-24A* obtained from Flybase (FB2019_02)(*58*). In some DGRP lines, heterozygous sites were encountered and, therefore, regions downstream of these sites were not properly sequenced. Sequences were aligned using Sequencer v.4.5 with the Dirty Data algorithm and default parameters. The c.-334_-333insACATTCAT indel alignments were manually curated for consistency with DGRP Freeze 2.0 (2L:3,718,040).

Having identified previously unknown indel variants, we identified the ancestral state by aligning the upstream indel variants and coding regions to closely related species. Orthologues of *Lectin-24A* were obtained from Ensembl Metazoa (accessed 19-11-2020) for *D. sechellia* (dsec_caf1\GM18119) and *D. simulans* (ASM75419v3\GD22727). These were the only two species containing one-to-one orthologues with > 75% identity. The gene region and 2,000bp upstream were obtained for each species. Then, the *D. sechellia* and *D. simulans* sequences and were aligned with Sanger sequences for DGRP-437 and DGRP-892. Geneious 11.0.3 was used for alignments using the native global alignment algorithm with free end gaps, 51% similarity cost matrix and gap open and extension penalties of 50 and 6, respectively. A total of five iterations were done with a guide tree used for alignments.

*Lectin-24A fluorescent reporter constructs*

*Lectin-24A* promoter reporter constructs were made using the pWallium20 backbone plasmid. An 8,494bp fragment of pWallium20 was created by restriction digest using HpaI and PstI enzmes (NEB # R0105S and R0140T) followed by gel purification and quantification using a Qubit HS DNA assay kit (Thermofisher # Q32854). The DGRP-437 (489bp upstream on translation start site) or DGRP-892 (453bp upstream on translation start site) *Lectin-24A* minimal promoter and was amplified using primers Lec500promF and lecpromvenusRv with Q5 2xMM PCR reagents (NEB # M0492S). The Venus open reading frame incorporating the ATG start codon of the *Lectin-24A* open reading frame was amplified using primers venusLecPromF and venusLecPromR using pDsRed-attP (Addgene: Plasmid #51019) as a template. The pWallium20 backbone and the two PCR fragments were then assembled using a HiFi assembly reaction (NEB # E5520) using an incubation at 50°C for 1 hour. Transformation plasmids were then purified by miniprep (Qiagen # 27104). Plasmid sequences were verified by sanger sequencing with primers Lec500PromRv, venusLecPromR and venusLecPromF. *Lectin-24A* promoter (LP) constructs containing different allelic combinations of the three upstream indels and six upstream SNPs (3717919_SNP, 3717922_SNP, 3717950_SNP, 3717956_SNP, 3718036_SNP, 3718037_SNP) that were polymorphic between DGRP-892 and DGRP-437. DGRP-892 alleles were substituted into the DGRP-437 LP sequence (437LP) by amplifying PCR fragments using primer combinations and template DNA as shown in Table S5. The fragments were then assembled with HpaI and PstI digested pWallium20 as for 437LP and 892LP. All constructs (Table S6) were injected into *D. melanogaster* line attP2Ar5 and transformants were selected by eye colour.

The regulatory activity of each lectin promoter reporter construct was evaluated by preparing samples of larval tissue lysate. At least eight independent samples were made for each promoter construct. Fifteen third instar larvae are collected for each sample, and tissuelysed in 100ul of PBS with about ten 1.0mm diameter zirconia/silica beads (Thistle Scientific # 11079110z) using Qiagen TissueLyser II (Qiagen # 85300) for 2 minutes at 30Hz. The samples were immediately spun down at 4000rpm for 10 minutes at 4°C, and 50ul of the supernatant for each sample were transferred into a well in a flat clear-bottom black polystyrene 96-well plate (Corning # 3603) for fluorescence reading with a SpectraMax iD3 Plate Reader (Molecular Devices) using the SoftMax Pro 7 software. Venus fluorescence intensity is measured with excitation at 485nm and emission at 535nm. The rest of the supernatant for each sample is used to measure the total protein level of the sample with a Bradford assay (Merck # B6916), following reagent protocol. The total protein level was used as a normalisation of Venus intensity. The whole larvae images in Figure 4C were taken with a Leica MZ16F Fluorescence Stereoscope fitted with a GXCAM HiChrome-S colour camera using the GXCAPTURE software captured at a 5ms exposure time. The GFP2 filter was used with an excitation of 480/40nm and emission measured at 510nm.

The assays took place over five days. We fitted a mixed effect linear model with relative Venus quantity as the response variable, the construct as a fixed effect and the experimentation date as a random factor. We performed the Tukey's multiple comparison test on the fitted model using *glht* function from the multcomp v1.4-14 R package (*56*) and adjusted the resulting *p*-values using the single-step method to identify the significant comparisons.

*Allele specific expression of Lectin-24A in DGRPs*

In order to understand the impact of upstream indels on expression, we selected multiple DGRP lines with varying combinations of the three upstream indels and tested if parasitoid infection upregulated *Lectin-24A* in those lines. 10 male flies from 20 DGRP lines with varying combinations of the upstream *Lectin-24A* promoter indels were crossed with 10 virgin females of DGRP-437. Females were allowed to lay eggs overnight in cornmeal vials with yeast. Adult flies were removed the next day and tipped into a new vial, which served as the second biological replicate. $F_1$ larvae in each vial developed for an additional 48 hours before they were infected with three female *L. boulardi* wasp strain G486 for three hours. 24 hours post infection, 10 larvae for each sample were homogenized in RTL buffer and RNA was extracted using an RNA easy miniprep kit (Qiagen: 74104), including optional on-column DNase treatment (Qiagen: 79254). cDNA was prepared using 1µl of template RNA with the GoScript Reverse Transcriptase system (Promega: A5003) primed with random hexamers, according to the manufacturer's instructions. For each $F_1$ cross, genomic DNA (gDNA) was also extracted from eight adult flies using the Blood and Tissue DNA extraction kit (Qiagen: 69504).

Sequencing libraries were prepared from each sample. PCR was performed (see section: Primer design and PCR for genotyping) on the gDNA for one of the biological replicates and cDNA from both biological replicates targeting the *Lectin-24A* gene region on chromosome 2L between 3,717,069 and 3,717,568 (dm6). *Lectin-24A* sequences obtained from Flybase (FB2020_03) were used to design primers TCGTCGGCAGCGTCAGATGTGTATAAGAGACAGACAAATGGAATACTTGTACGGAAAT (forward) and GTCTCGTGGGCTCGGAGATGTGTATAAGAGACAGGACGCCCAGGAAGTATGATT (reverse) that include adapter overhang sequences. Two technical replicates of each PCR reaction were performed. PCRs were performed using Q5 DNA polymerase (NEB: M0491S) in a total volume of 50µl using 2µl of template cDNA or gDNA. PCR amplicons were cleaned using 20µl of Ampure XP beads (Beckman: A63880) and eluted in 52.5µl of 10mM Tris, pH8. An index PCR reaction was carried out using 5µl of cleaned PCR as template, 25µl of 2 x Kapa HiFi hot start PCR mix (Roche: KK2601), 5µl of each of Truseq XT index 1 and 2 obtained from the Nextera XT DNA Library Prep Kit (Illumina: 15052165) and 10µl of water. After cleaning with 56µl of Ampure beads the libraries were eluted in 27.5µl of Tris pH 8.0 and analyzed/quantified by agarose gel electrophoresis. Libraries were pooled and sequenced on MiSeq nano 300 cycle kit, paired-end sequencing at the DNA sequencing facility at the University of Cambridge Biochemistry Department.

Following sequencing, Illumina BaseSpace Sequence Hub was used to convert Binary Base Call files into FASTQ files for all reads passing filtering. We mapped sequenced reads to a FASTA sequence containing the region of interest obtained from Flybase (FB2020_03) using bwa-mem (*65*) marking shorter split hits as secondary. We also mapped reads to references incorporating alternate alleles for eight SNPs segregating in the 21 DGRP lines that we tested to estimate read mapping bias (Table S7). Variant calls for the parent DGRP lines were obtained from DGRP Freeze 2.0. Aligned reads were reordered and read groups were added using Picard Toolkit v.2.8.1 (*66*) and the resulting files were sorted using samtools 1.9 (*67*). The aligned and processed reads were indexed with Picard Toolkit. We called variants and estimated allele depth in the $F_1$ gDNA and cDNA using GATK3.7-0 HaplotypeCaller (*68*). Reads were assigned to DGRP alleles using a SNP (dm6, 2L:3717102) where all tested DGRP lines had the reference allele and DGRP-437 had the alternate allele (Table S7). A Welch two sample t-test was used to test for differences in allelic expression in the $F_1$ hybrids generated from DGRP-437 and DGRP-892. Indel haplotypes for tested DGRP lines were obtained from the Sanger sequencing calls and association between indel haplotype and allele expression was tested using a $\chi^2$ test. Further, we tested for association between the counts of reference and alternate counts and indel haplotype for cDNA using a quasibinomial GLM.

*qPCR assays of Lectin-24A expression in DGRP lines*

We estimated gene expression using qPCR in separate group of 18 DGRP lines with varying indel haplotypes, partially overlapping with the ones referred to above for studying ASE. We used the 2019 infection assay (see section: resistance assay) and RNA extractions and cDNA protocols were the same as above. Primer3 (http://primer3.ut.ee, last accessed August 2020) was used to design the GATACTGCTTCCCCACCCTG (forward) and ATCTGATCCTTGCGGGCTTC (reverse) primers to amplify a 108bp fragment in the intron-less *Lectin-24A* coding region. RNA was incubated with DNase I (Qiagen RNase-Free DNase set: 79254) for 15 minutes per manufacturer protocol and qPCR was done as described for wasp fragment amplification (see section: Fine mapping with resistant recombinant adults). In addition, we included a 'no RT' (NRT) control in which the RNA was not reverse transcribed as a control for residual DNA contamination. Two replicates were performed per line and the average expression was taken. The expression of the housekeeping gene *RpL32* was estimated with the aid of primers obtained from (*69*) and ΔCT was calculated as $CT^{Lectin\text{-}24A}$–$CT^{RpL32}$ . - ΔCT is an estimate of the $\log_2$ relative *Lectin-24A* expression. A linear regression was done to test the significance of the association between indel haplotype and $\log_2$ relative *Lectin-24A* expression. A Tukey's honest significance test was done to compare $\log_2$ relative *Lectin-24A* expression between indel haplotypes.

A separate experiment was done to estimate fold change in DGRP-437 and DGRP-892 flies infected with parasitoid wasp or injected with wasp and oil droplets, as detailed in (*54*). We assessed expression at 6 and 18 hours post exposure to parasitoid wasps. The same protocol as above was used except that no DNase I treatment was done prior to qPCR and there were four replicates for each line and treatment combination. ΔCT from the NRT controls was used to set detection thresholds for expression for each sample. We did an analysis of variance on the model below to test the significance of the fixed effects on ΔCT.

$$\text{lm}(\Delta\text{CT} \sim \text{Line} + \text{ Time post wasp exposure} + \text{Exposure type, i. e. infection, injection})$$

*Variant identification in Drosophila Genome Nexus samples*

We used *Drosophila* Genome Nexus data generated and complied by (*44*) to investigate molecular evolution of *Lectin-24A*. First, fastq sequence data was obtained from NCBI SRA using SRA toolkit v2.9.6-1 (http://ncbi.github.io/sra-tools/). Then, Trimmomatic v.0.36 (*70*) was used to clip adaptor sequences and remove low quality bases from reads. We clipped adaptors sequences using reference file TruSeq3-PE.fa with the following parameters: seed mismatches = 2, palindrome clip threshold = 20, simple clip threshold = 10, minimum adapter length = 1 and keepbothreads = TRUE. For single end reads, we used the reference file TruSeq3-SE.fa. Further, we removed the last three bases from reads, filtered strings of low-quality bases found in sliding windows of sizes 4bp where quality dropped below 20 and ensured reads had a minimum size of 36bp. We mapped trimmed reads to the *D. melanogaster* reference (r5.13) (*71*) attained from Flybase (FB2019_02) (*58*) using bwa-mem (*65*) and reordered and added read group information specifying the sample and library names and Illumina planform using Picard Toolkit v.2.8.1 (*66*). Samtools 1.9 (*67*) was used to retain mapped reads with alignment quality above 20. For paired end data, we only retained properly paired and mapped reads. Reads were subsequently sorted by coordinate. Then, duplicate reads were marked and resulting alignment files were indexed using Picard Toolkit. Reads near indels were realigned and mis-encoded quality scores were fixed for some samples using GATK3.7-0 (*68*). CRISP v0.7 (*72*) was used to call variants in all samples jointly using a poolsize of 1. Further, the following default parameters were used: quality value offset = 33 for Sanger format, minimum base quality to consider a base for variant calling = 10, minimum read mapping quality to consider a read for variant calling = 20, minimum number of reads with alternate allele required for calling a variant = 4, log10 threshold on the contingency table *p-value* for calling position as variant = -3.5, log10 threshold on the quality values based *p-value* for calling position as variant = -5, maximum number of permutations for calculating contingency table *p-value* = 20000, filter reads with excessive number of mismatches and gaps compared to the reference sequence = TRUE, identify overlapping paired-end reads and treat as single read in the overlapping region = TRUE, reference allele bias for targeted sequencing data = 0.5, EM algorithm will be used for estimating pooled genotypes and calling variants = TRUE, PCR duplicates will be ignored = TRUE. The reference genome was split into 11 regions to perform variant calling in parallel. SnpEff v4.3t (*73*) was used to annotate variants using the dm5 database. *Lectin-24A* region SNPs were subset from the main variant call format (VCF). Preliminary investigations indicated a large deletion occurring in the *Lectin-24A* coding region in some samples. This prompted us to use pindel v0.2.4s (*74*) to systemically call deletions in the *Lectin-24A* region alignment files. Indeed, a 171bp region (TCCTGGGCGTCAACGAAAACACAAAAACCGGTGACTTTGTGTCTGCAGCCTCCGGAAAAAGTTGTT TGTATCACGAGTGGGGTCCTGGTGAACCCCATCACAATAATGACCAAGAGCGATGCGTTTCAATCTT GCGAAAACTCATGCATGTGGGCAATTGTACTTATGAAA) was substituted with a 6bp sequence (TAATCAT) near the 3′-end of *Lectin-24A* in some lines, which results in a premature stop codon following the first base in the substituted sequence. We identified samples as having the coding deletion if they were fully genotyped for regions flanking the coding indel but did not contain any variant calls in the region with the presumed deletion. We confirmed calls by using pindel to call the deletion and by manually checking alignment files for split reads. The deletion calls were manually checked using Integrative Genomics Viewer v2.9.2 (*75*) for split reads. The deletion calls were added for further analyses. In order to genotype samples for the three upstream indels of *Lectin-24A*, trimmed reads were mapped to a reference modified using GATK3.7-0 FastaAlternateReferenceMaker to include the insertions for all three indels and the above alignment processing steps were repeated. *Lectin-24A* region alignments were isolated from the main file after coordinate-based sorting and downstream processing steps and variant calling was performed.

Variants that were identified were masked and quality-checked using the R script dgn_lectin.R *Lectin-24A* region SNPs and the upstream indels were combined into a single file. Coding and noncoding region SNPs were named using protein and DNA based HGVS nomenclature, respectively, obtained from SnpEff annotations with the aid of the R package sjmisc (*76*). Variant names were modified to reflect ancestral and derived alleles rather than reference and alternate alleles. Indel names were manually given using HGVS rules and upstream SNP positions were modified to account for the ancestral upstream insertions, which are not found in the unmodified *D. melanogaster* reference. SRA run names were converted to sample names using the individual description provided by (*44*). For individuals with multiple SRA associations, genotype calls made in any of those sequencing files were retained unless they conflicted with each other. Multiallelic sites were only retained if all alleles had the same effect as predicted by SnpEff. The very recent migrant ZW184 was removed from population genetic analyses (see (*77*)). Regions identified as being affected by identity by decent, admixture or heterozygosity by (*44*) were masked from analyses. Following sample-specific site masking, individuals with more than 50% missing data for *Lectin-24A* region variants were removed from analyses. Only the 26 populations that each contained more than three samples were retained for the analyses. Further, only variants that had an allele frequency >0.05 in two or more populations and with genotype quality greater than or equal to 10 were retained.

The consistency and reproducibility of the variant calls were confirmed by comparing to the constituent variant call datasets for subsets of the *Drosophila* Genome Nexus through Sanger sequencing. There was an exact match of variant calls using the approach detailed above and the calls published in DGRP Freeze2.0 (*42, 52*) and the Global Diversity Lines (*77*) (Table S8). Further, Sanger sequencing (using approach detailed in section: Fine mapping with resistant recombinant adults) of a subset of lines from the SD and SP populations from Southern Africa, received through the courtesy of John Pool, indicated an exact match for key upstream indels and loss-of-function variant calls including the 165bp deletion segregating in some lines (Table S8). The following forward (GGCGCCTCCTTGGACTATTT) and reverse (CTGTTGCCGAGCACAATTGTCATTTCTTCATTTCGCCAC) primers were used for amplifying the fragment for sequencing. We performed qPCR for *Lectin-24A* expression following infection and no infection of 14 lines from Southern Africa containing one or more of the presumed loss-of-function mutations in the *Lectin-24A* coding sequence. Our goal was to test if any loss of expression mutations occur alongside the loss-of-function mutations, nullifying any effect of the latter. The method used in section '*qPCR assays of Lectin-24A expression in DGRPs*' was repeated for qPCR assay.

*Drosophila Genome Nexus population genetics analyses*

Ancestral and derived status for variants in the 200 kbp surrounding *Lectin-24A* was obtained from dm5 using samtools. Using species-specific BLAST from Flybase (FB2019_02) (*58*), the orthologous regions in *D. sechellia* and *D. simulans* were obtained. Sequences from the three species were then aligned using MAFFT as implemented on the EBI server (*78*). Following multi-species alignment, the allele that also occurred either in *D. sechellia* or *D. simulans* was designated as the ancestral state. However, if both the reference and alternate alleles or neither of them were found in the other two species, an ancestral state classification was not given, and these sites were removed from analyses.

VCFtools v0.1.15 (*79*) was used to calculate per-site Weir and Cockerham $F_{ST}$ and weighted pairwise $F_{ST}$ along the entire *Lectin-24A* coding region between geographic regions. Gene diagrams for the per site $F_{ST}$ plot was generated using the R package gggenes v0.4.1 (https://wilkox.org/gggenes/index.html). In order to calculate the significance threshold, per site $F_{ST}$ was calculated for autosomal SNPs. The same variant processing, masking and filtering steps as done for *Lectin-24A* region-specific analyses were applied for the background SNPs. However, no window-level masking for uncalled genotypes was applied and 1,039 individuals from the eight geographic regions were used for calculating the $F_{ST}$ for background SNPs. $F_{ST}$ was only calculated when there were at least 22 populations each containing at least four samples genotyped for a given site. The top 1% and 0.1% $F_{ST}$ percentiles were calculated using 8,440,512 autosomal SNPs.

For subsequent *Lectin-24A* specific analyses, only populations that contained four or more individuals where at least 50% of the *Lectin-24* region was genotyped were used resulting in 672 individuals from 26 populations being retained for the analyzed. *Lectin-24* coding region $F_{ST}$ was obtained using VCFtools. The R packages adegenet v2.1.3 (*80*) and hierfstat v0.5-7 (*81*) were used to calculate population-level allele frequencies. Pairwise linkage disequilibrium between sites was estimated using the R package LDheatmap v1.0-4 with input VCFs read and processed into a genotype matrix using the R package VariantAnnotation v1.36 (*82*).

A neighbor-joining was constructed to infer the relatedness between samples. Processed genotype calls were formatted into a VCF. An indexed *D. melanogaster* reference was used alongside bcftools v1.11-21 (*83*) consensus to attain a reconstructed coding region alignment for all analyzed samples. A neighbor joining tree was constructed using the reverse complement of the alignment with the software MEGAX (*84*). The tree was constructed using a Jukes Cantor substitution model. The best fitting substation model was assessed using the R package phangorn v2.5.5 (*85*). The R package APE v5.4-1 (*86*) was used to read and process alignment files. For tree construction, uniform rates were applied for all sites and the pairwise deletion option was used for dealing with gaps and missing data. Trees were further processed in R using the package ggtree v2.4.1.

*Population branch statistic*

We estimated the population branch statistic to test to positive selection on premature stop codons in *Lectin-24A*. To provide a null distribution of this statistic, we also estimated the population branch statistic for other premature stop codons elsewhere in the genome. First, we identified the ancestral state of stop gained mutations by extracting 100bp on either side of the mutation using samtools. These query sequences were used to perform a nucleotide BLAST using the discontiguous megaBLAST (*87*) with the subject organism restricted to either *D. sechellia* or *D. simulans*. Alleles for sites containing premature stop codons were designated as the ancestral state if they occurred in either *D. sechellia* or *D. simulans*. Ancestral status could not be identified for sites where both the reference and alternate alleles were present in *D. sechellia* and *D. simulans*, and these sites were excluded from further analyses. Only premature stop codons designated as being the derived state in *D. melanogaster* were used in our analyses. The gene transfer file for *D. melanogaster* reference (*71*) was used for calculating the distance of the premature stop codon from the actual stop codon. Premature stop codons occurring with 10bp of the actual stop codon were excluded from analyses as they are less likely to have a significant impact on protein function. When premature stop codons occurred within exons used by multiple transcripts, the distance to the closest stop codon was calculated. Further, premature stop codons occurring in low recombining regions (<1 cM/Mbp) were excluded from analyses as it is difficult to ascertain if selection is acting on those stop codons or a nearby linked variant. Recombination rates for regions where premature stop codons occur were obtained from (*88*). Allele frequencies for stop codons in the three geographic regions (227 samples from North America, 132 samples from Europe and North Africa and 317 samples from Southern Africa) with the largest number of published genomes out of the 1,039 available in the *Drosophila* Genome Nexus were calculated. Only polymorphic sites where the premature stop codon had greater than 5% frequency in at least one of the three regions were used for subsequent analyses, resulting in the identification 211 premature stop codons in *Drosophila* Genome Nexus autosomes. Pairwise $F_{ST}$ for the premature stop codons in these three regions was calculated using VCFtools and we calculated the population branch statistic for each stop codon using the equation from (*45*). Population branch statistic tress were plotted using the R package APE v5.4-1 (*86*).

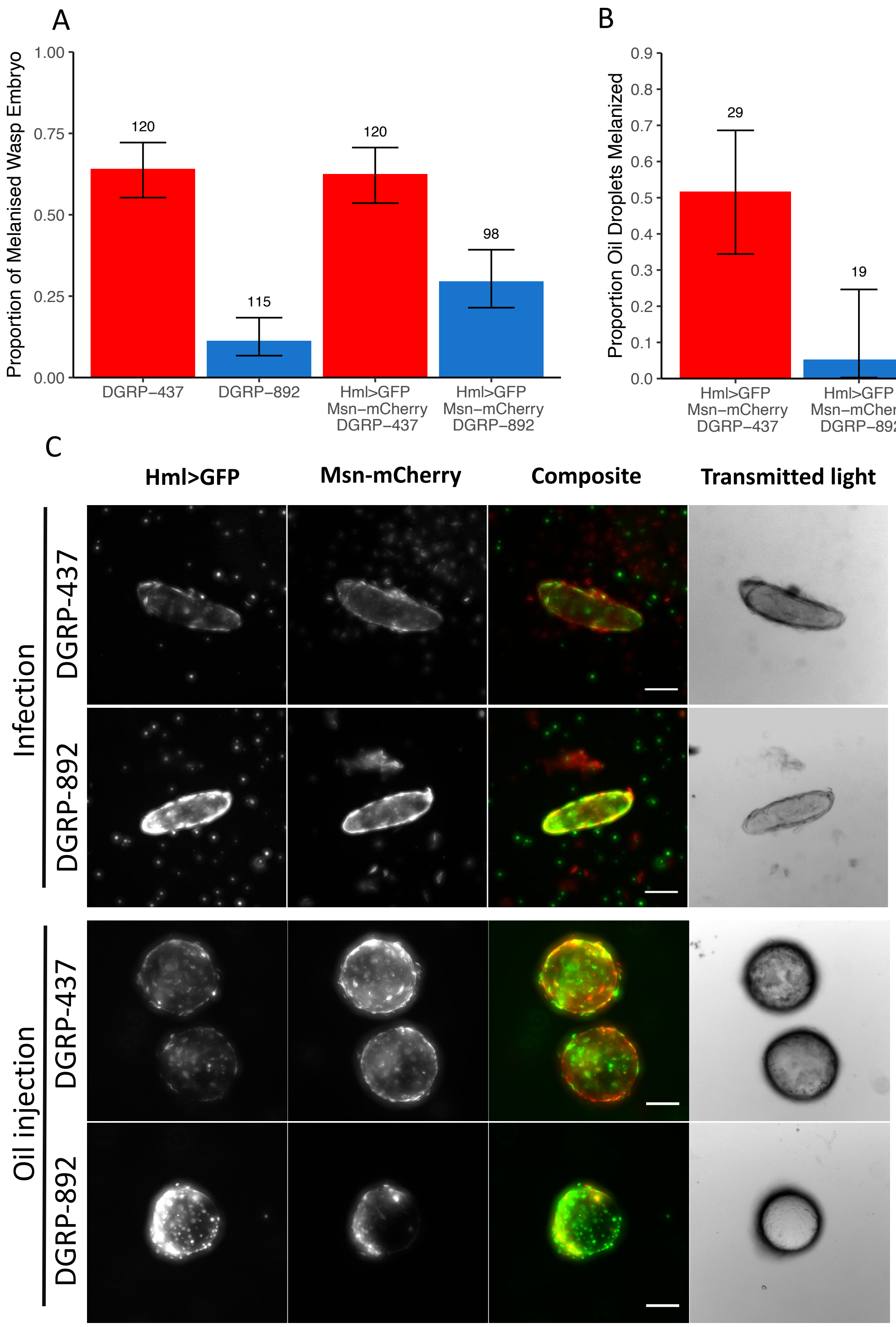


**Figure S1. Recruitment of plasmatocytes and lamellocytes following wasp infection in resistant and susceptible lines.** (A) Proportion of melanized *L. boulardi* wasp embryos 24

hours post infection (hpi) in the $F_1$ progeny resulting from crosses between lines carrying the HmlΔ-GAL4 UAS-GFP and Msn-mCherry constructs and DGRP-437 or DGRP-892. DGRP-437 and DGRP-892 are included as controls. (B) Proportion of oil droplets containing homogenized male *L. boulardi* that were melanized 24 hpi in the $F_1$ progeny resulting from the same cross. In (A) and (B) error bars show the 95% confidence interval. (C) Images taken with red and green channels show the hemocytes that have attached to the wasp embryo or the oil droplet containing homogenized wasp 24 hours post infection or injection of the $F_1$ larvae. The pan-hemocyte driver HmlΔ-GAL4 UAS-GFP and lamellocyte-specific Msn-mCherry were used to visualize plasmatocytes and lamellocytes respectively. Scale bars correspond to 100μm.

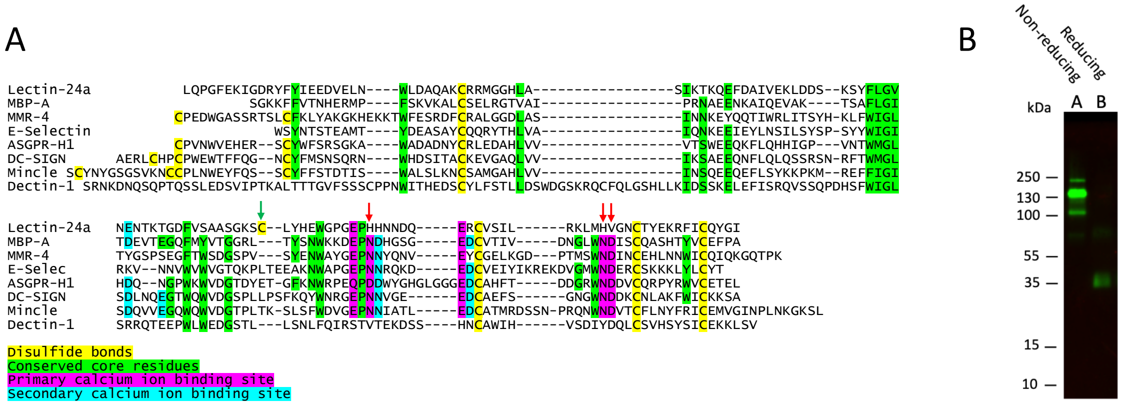


**Figure S2. The carbohydrate binding domain of Lectin-24A and tetramer formation.** (A) Alignment of the carbohydrate recognition domain from Lectin-24A and other C-type lectins. Key conserved residues that define the domain architecture are highlighted in yellow (cysteines) and green (other residues). The presence of these residues confirms that the domain will likely have a C-type lectin fold. Other colors highlight residues needed to form $Ca^{2+}$ binding sites, including the primary $Ca^{2+}$ binding site that forms the carbohydrate-binding site. The red arrow highlights three essential residues in the primary $Ca^{2+}$ binding site that have been lost from Lectin-24A. Note these have also been lost in Dectin-1, which has a C-type lectin fold yet binds sugar independently of $Ca^{2+}$. The green arrow shows an additional cysteine residue (4 additional cysteines are found outside the carbohydrate recognition domain). (B) Lectin-24A secreted from *Drosophila* cells forms b-mercaptoethanol (2-ME) sensitive tetramers. Samples under reducing (with 2-ME) conditions exhibit the apparent 36kDa polypeptide size predicted from translation of the open reading frame of *Lectin-24A* with the FLAG and strep tags. Samples under non-reducing conditions (without 2-ME) exhibit a main polypeptide band of 140kDa.

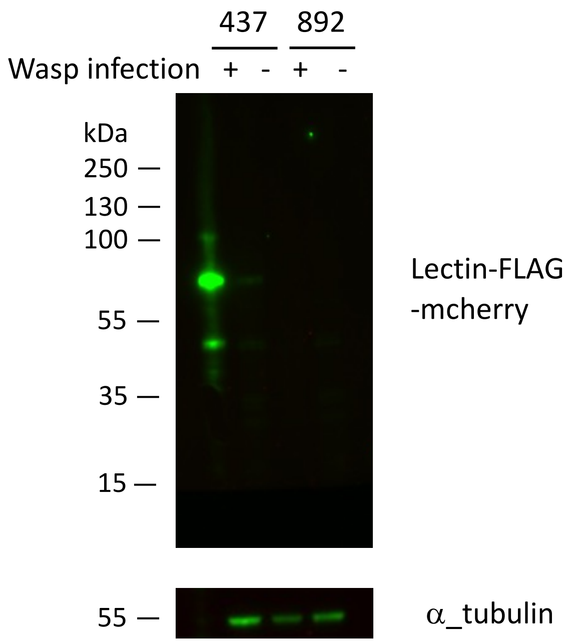


**Figure S3. Transgenic FLAG-mCherry-tagged Lectin-24A expressed in *Drosophila* larvae is upregulated after wasp infection.** A 2.5kb promoter fragment and the open reading frame of *Lectin-24A* from DGRP-437 or DGRP-892 were cloned to incorporate a FLAG and mCherry tag at the C-terminus. Transgenic flies were generated, and larvae were infected or left uninfected with *L. boulardi* G486. Protein extracts from these larvae were analyzed by Western blotting using an anti-FLAG tag antibody to detect tagged Lectin-24A. Blots were stripped and re-probed with anti α-tubulin antibody to show total protein loading. Tagged DGRP-437 Lectin-24A exhibits a predominant band of approximately 70kDa compared with the expected size of 60kDa. FLAG-mCherry-Lectin-24A is upregulated upon infection (437 + compared to 437 -).

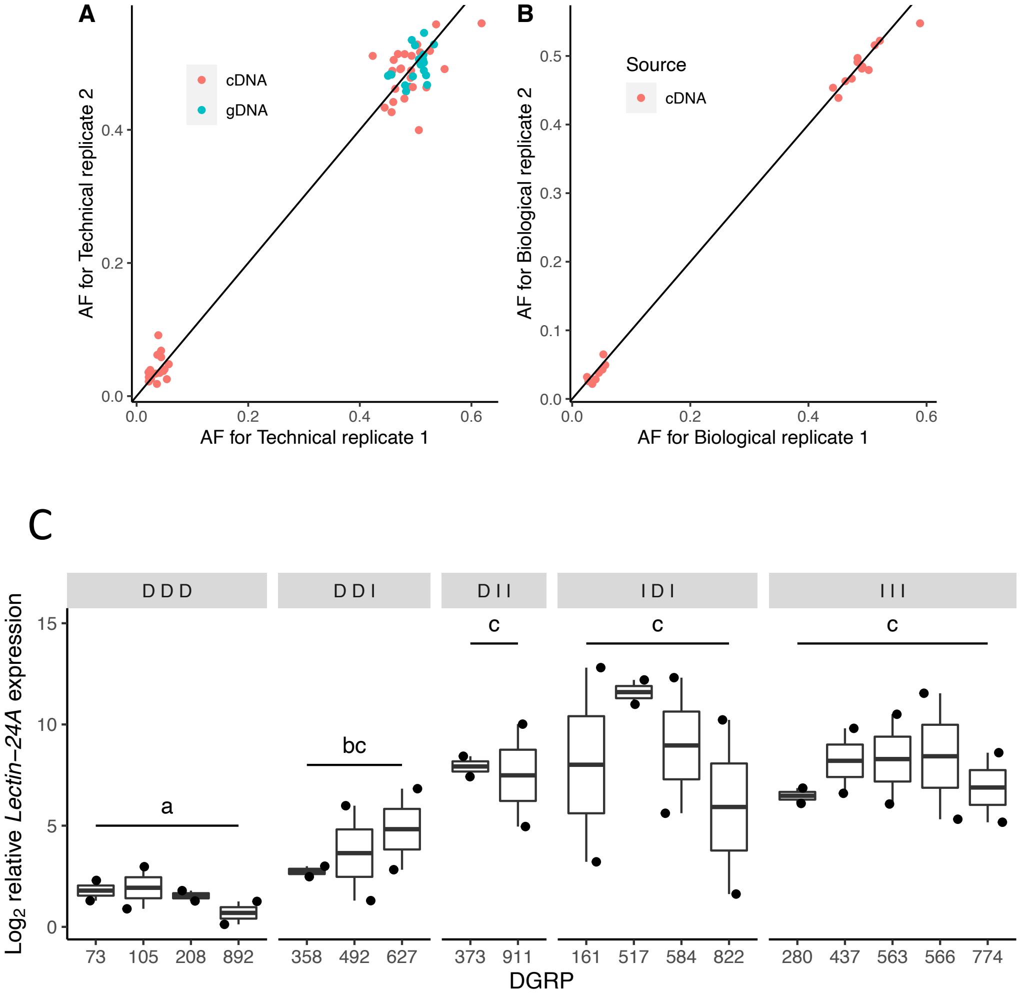


**Figure S4. Genetic variation in in the expression of *Lectin 24A*.** (A and B) Comparison of replicate measurements of *Lectin-24A* allele-specific expression. Panel A compares two technical replicates on the same nucleic acid sample, and Panel B compares two biological replicate measurements on each inbred line. Each line was crossed to the high expression line DGRP-437, *Lectin-24A* was amplified by PCR from genomic DNA (gDNA) or mRNA (cDNA), and then Illumina sequenced. The axes represent the frequency of reads derived from the test line (AF) as opposed to the DGRP-437 reference. There were no biological replicates of the genomic DNA (gDNA) controls. The read counts from the two technical replicates were summed for each biological replicate. (C) Expression of *Lectin-24A* in inbred lines from the DGRP panel 24 hours post infection. Lines were selected to include different indel haplotypes upstream of the gene. The indel haplotype is ordered as c.-439_-433del, c.-334_-333insACATTCAT and 21bp indel (c.-171_-151del), and the deletion ‘D’ or insertion ‘I’ state for each indel is depicted. Each point is an independent biological replicate and is the mean of two biological replicates. The haplotypes were compared with Tukey’s honest significant difference test, $p<0.05$ between groups, $p>0.05$ within groups. Expression is relative to the reference gene *RpL32.*

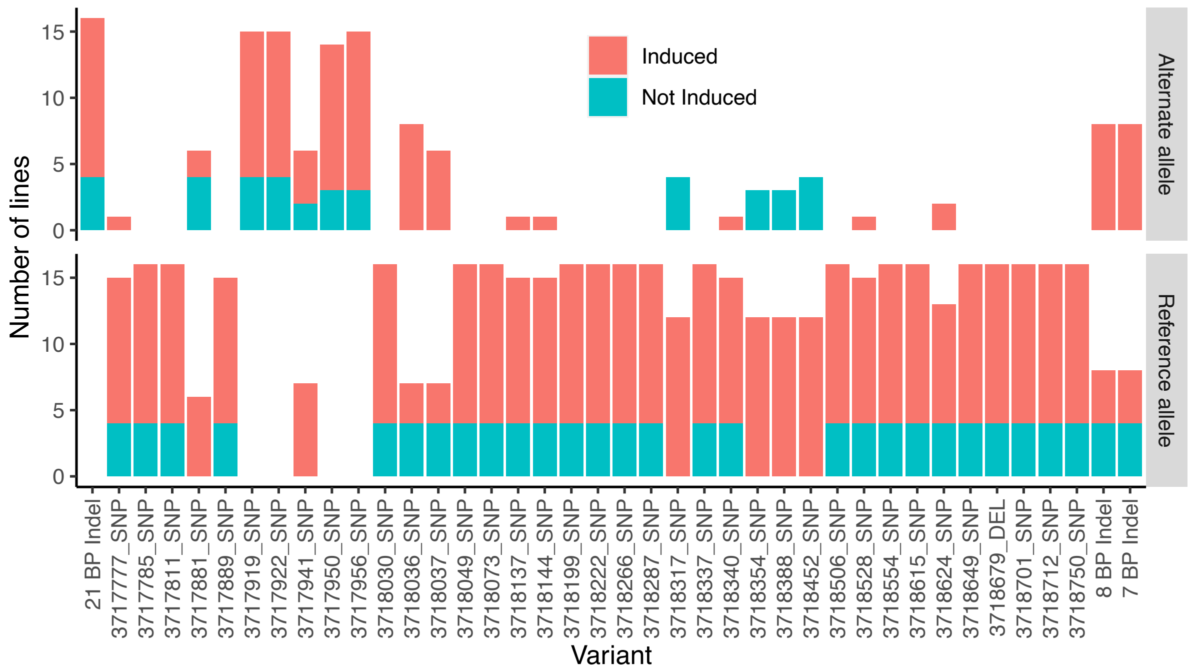


**Figure S5. Association of alleles for variants 1,000bp upstream of the *Lectin-24A* start codon with capability to induce expression following parasitoid infection.** Sixteen DGRP lines which all carried the insertion allele for the 21bp indel that permits expression were studied. The sixteen alleles were classified as to whether they were induced after infection based on allele-specific expression data. Only biallelic sites shown.

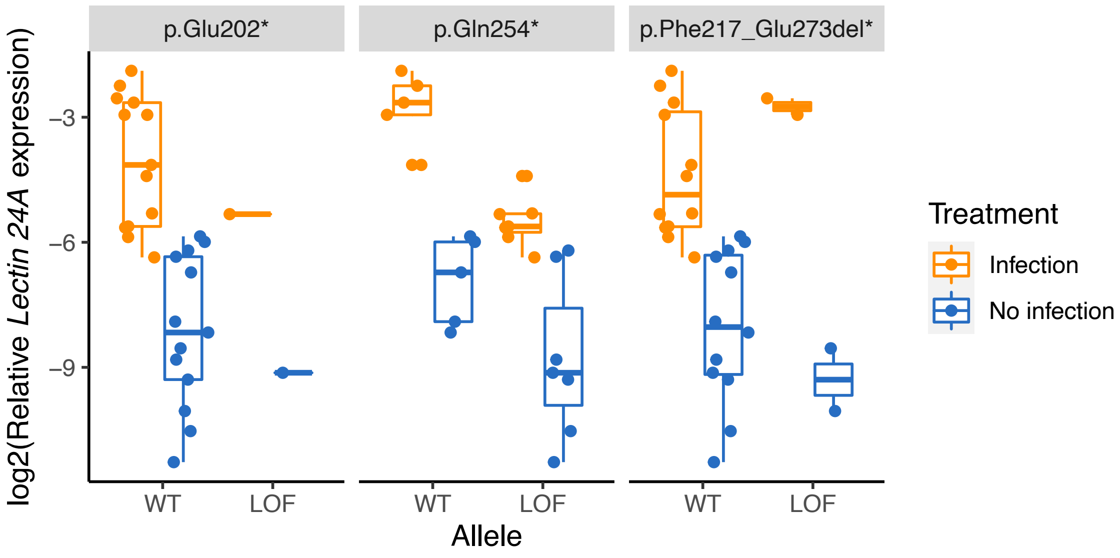

**Figure S6. Expression of *Lectin-24A* in Southern African lines with premature stop codons.** *Lectin-24A* expression is calculated relative to *RpL32*. LOF lines carry the stop codon and WT lines do not. Expression was measured 24 hours post infection with parasitoid wasps.

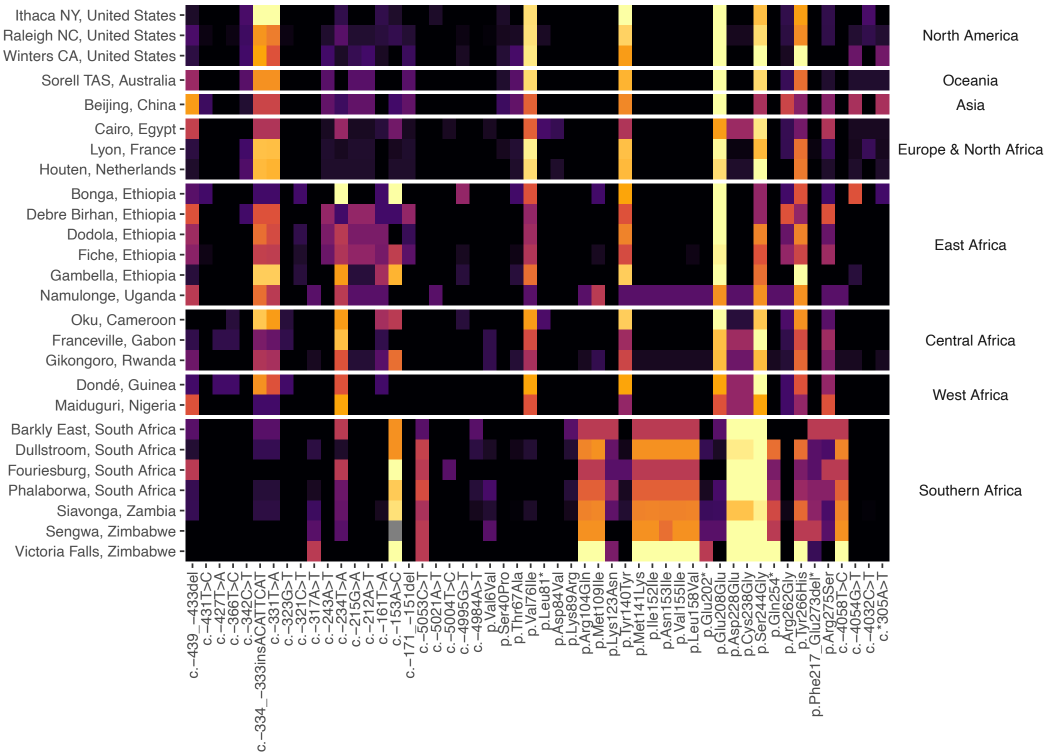


**Figure S7. Geographic variation in the derived allele frequency of variants in *Lectin-24A*.** Includes variants found in 412bp upstream of the start codon (inclusive of three promotor indels) and 67bp downstream of the stop codon. Populations (left) are grouped into regions (right). Data is from 672 genomes in the *Drosophila* Genome Nexus that were genotyped for >50% of the *Lectin-24A* gene region.

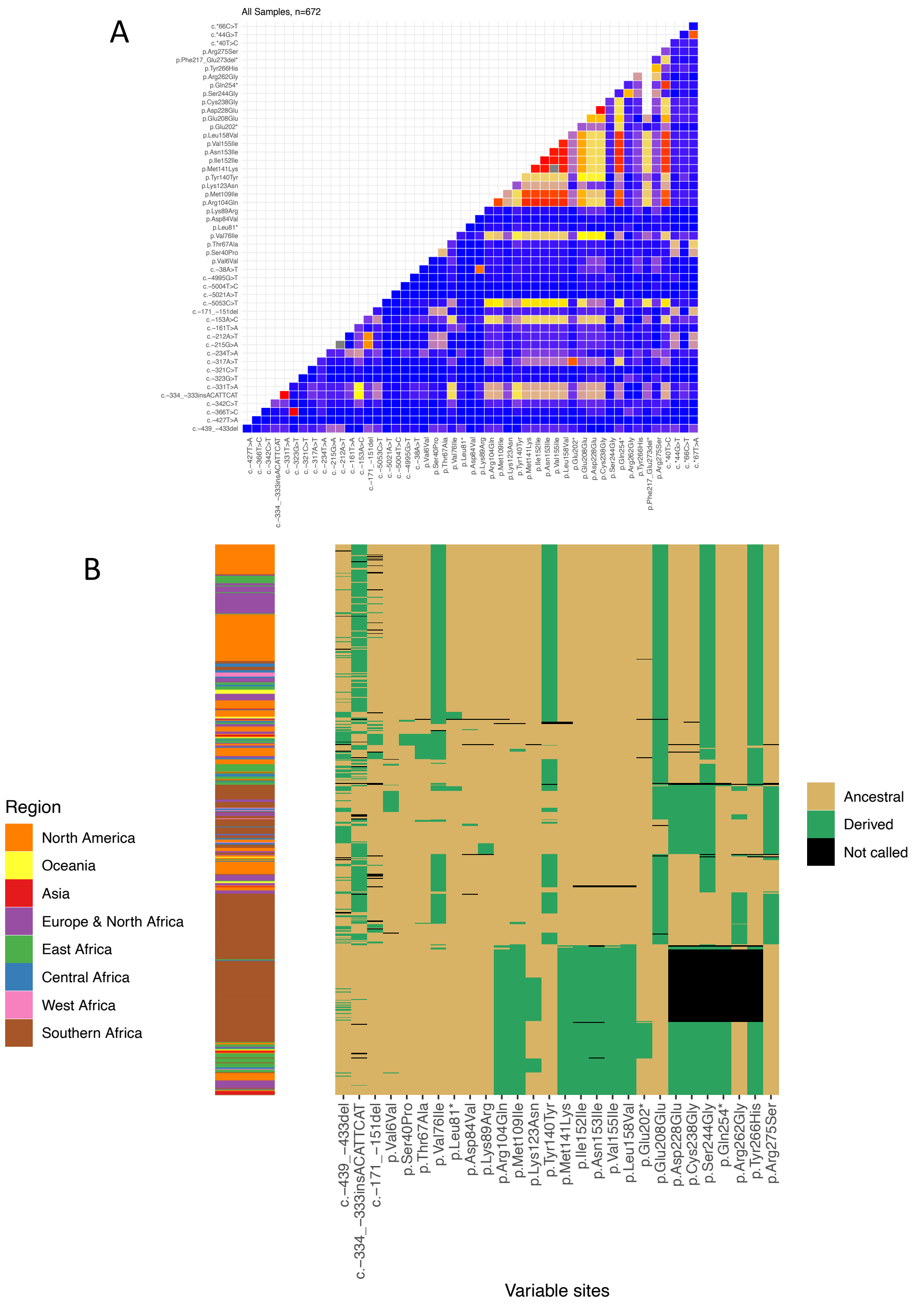


**Figure S8. Haplotype Structure in *Lectin-24A*.** (A) Linkage disequilibrium between *Lectin-24A* coding sequence and variants found in 412bp upstream of the start codon (inclusive of three promotor indels) and 67bp downstream of the stop codon. LD across all 672 samples of the *Drosophila* Genome Nexus that were genotyped for >50% of the *Lectin-24A* region are shown. (B) Genotypes of upstream indels, non-synonymous and stop polymorphisms in 672 *Lectin-24A* sequences. Samples were clustered using a neighbor-joining tree and color-coded by the region from where they were obtained. Upstream indels were not used for tree construction but are shown in the genotype matrix. Note that samples with blocks of variants that are not called in the 3’ end of the *Lectin-24A* coding sequence have the 165bp deletion p.Phe217_Glu273del*. In both panels, data are from the *Drosophila* Genome Nexus.

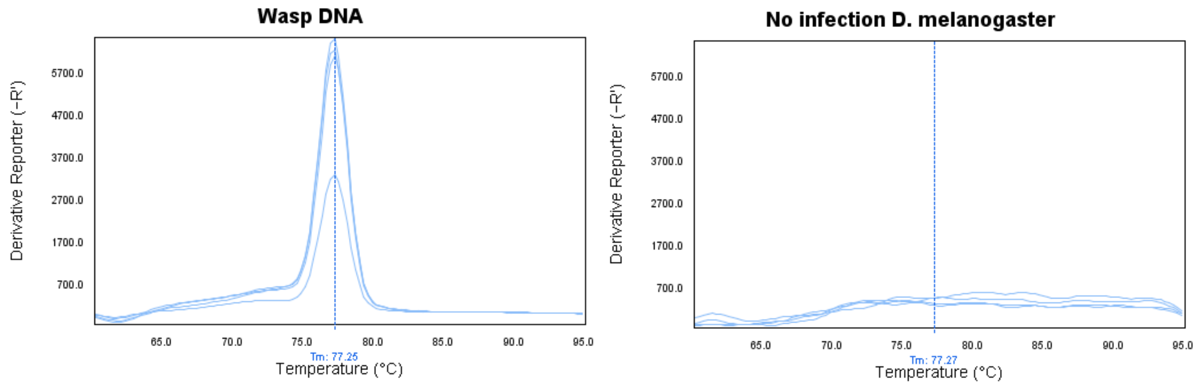


**Figure S9. Amplification of parasitoid wasp DNA and DNA from uninfected DGRP-437/DGRP-892 hybrids using wasp-specific primers.** Four technical replicates for each DNA were included. The peak primer melting temperature (Tm) under wasp DNA amplification is indicated by a vertical line.

```
>Wild-type Lectin-24A ORF
ATGTTTAGATTGTCAGTCTTAGTTCTGAACTTACTCCTTGTTAGCCATGAATTCTCGGCAGGAACCGCGAAAATTGAAAT
TCAGCCATTACCTGCCCTATGCAACGGATACTGCTTCTCCACCCTGAAG----CCAGTTATGGAGTATGTTGCTATCCAC
CAGGACAAATGGAATACTTGTACGGAAATATTAGCGAACGAAACCCGCAAGGATCAGATCCAGTTGAATATCCAGCTGGA
TGCCTTGAAAGCAGACGTTTCCAACATAAAGGCATCGCAGCTGTCAAAGGATGAGAAGCTGGACAGGATGGAGCGAGAGC
AGTTTGCCATGCATGAATCTTTGGAGACCATCAATCGGTATCTTACAGTGAAACTGGACAGAACGAAATTGCAGCTTGAG
GCGATCAAAAACACAATGGATTACATGAAGGCCCAAATGGATGGCTATTTCTCAGCCATAAATGGAGTTCAATGCCTACA
GCCAGGATTTGA-GAAGATAGGCGATAGATATTTTTACATCGAAGAAGATGTTGAGCTAAATTGGCTGGATGCTCAGGCC
AAATGTCGTCGAATGGGAGGTCACCTAGCCTCTATAAAAACTAAACAGGAGTTTGACGCAATCGTAGAGAAACTCGATGA
TTCGAAATCATACTTCCTGGGCGTCAACGAAAACACAAAAACCGGTGACTTTGTGTCTGCAGCCTCCGGAAAAAGTTGTT
TGTATCACGAGTGGGGTCCTGGTGAACCCCATCACAATAATGACCAAGAGCGATGCGTTTCAATCTTGCGAAAACTCATG
CATGTGGGCAATTGTACTTATGAAAAAGATTCATATGCCAGTATGGCATC
>Mutant Lectin-24A
-----------------------------------------------TGNATTCTCGGCAGGAACCGCGAAAATTGAAAT
TCAGCCATTACCTGCCCTATGCAACGGATACTGCTTCTCCACCCTGAAGTTATCCAGTTATGGAGTATGTTGCTATCCAC
CAGGACAAATGGAATACTTGTACGGAAATATTAGCGAACGAAACCCGCAAGGATCAGATCCAGTTGAATATCCAGCTGGA
TGCCTTGAAAGCAGACGTTTCCAACATAAAGGCATCGCAGCTGTCAAAGGATGAGAAGCTGGACAGGATGGAGCGAGAGC
AGTTTGCCATGC--CAATCTTTGGAGACCATCAATCGGTATCTTACAGTGAAACTGGACAGAACGAAATTGCAGCTTGAG
GCGATCAAAAACACAATGGATTACATGAAGGCCCAAATGGATGGCTATTTCTCAGCCATAAATGGAGTTCAATGCCTACA
GCCAGGATTTNAAAAAAAATAGGCAATNAAAATTTTTANNTCCAAAAAAAAAGTTTANCTAAATTGGCTGGATGNTCAGGCA
AAA-----------------------------------------------------------------------------
--------------------------------------------------------------------------------
--------------------------------------------------------------------------------
--------------------------------------------------
```

**Figure S10. Frameshift mutations introduced in the resistant *Lectin-24A* allele.** A 4bp insertion and a 2bp deletion (indicated in yellow), which were 129bp and 328bp downstream of the start codon, respectively, were introduced via Cas9 mediated mutagenesis. Their locations correspond approximately to the target region of the guide RNAs. The first 4bp insertion results in a frameshift and causes a premature stop codon (indicated in red) that results in a truncated 97 amino acid protein missing the entire carbohydrate binding domain.

**Main Cross**

| | | | | | | |
|---|---|---|---|---|---|---|
| f / FM7 | II / II | III / III | 1 x | w / y | If / CyO | MKRS / TM6B |
| attp[gRNA] / attp[gRNA] | II / II | III / III | 2 x | FM7 / y | If / II | MKRS / III |
| attp[gRNA] / FM7 | II / If | III / MKRS | 3 x | 437 / y | 437 / 437 | 437 / 437 |
| Cas9 / FM7 | II / CyO | III / TM6B | 4 x | attp[gRNA] / y | If / 437 | MKRS / 437 |
| Mutagenesis | | | | | | |
| attp[gRNA] / Cas9 | 437 / CyO | 437 / TM6B | 5 x | w / y | If / CyO | MKRS / TM6B |
| | | | | Single male | | |
| 892 / 892 | If / CyO | MKRS / TM6B | 6 x | X[Mess] / y | 437[mutant] / CyO | 437 / TM6B |
| 892 / 892 | If / CyO-GFP | MKRS / TM6B | 7 x | 892 / y | 437[mutant] / CyO | 437 / TM6B |
| 892 / 892 | 437[mutant] / CyO-GFP | 437 / TM6B | 8 x | 892 / y | 437[mutant] / CyO-GFP | 437 / TM6B |

Stocks

| | | | | | |
|---|---|---|---|---|---|
| f / FM7 | II / II | III / III | 892 / 892 | If / CyO | MKRS / TM6B |
| w / y | If / CyO | MKRS / TM6B | 892 / 892 | If / CyO-GFP | MKRS / TM6B |
| attp[gRNA] / attp[gRNA] | II / II | III / III | Cas9 / Cas9 | II / II | III / III |
| 437 / 437 | 437 / 437 | 437 / 437 | | | |

**Figure S11. Crosses to generate germline mutants for *Lectin-24A* in the resistant ($X^{892}$; $II^{437}$; $III^{437}$) background.** The stock attP2Ar5 containing two guide RNAs, which were integrated at an attP site on the X chromosome, and the Cas9 containing stock (BDSC#51323) were used for mutating *Lectin-24A* alleles. GFP tagged CyO was eventually removed from the mutated lines.

**Table S1. Table of 107 genes under the chromosome II quantitative trait locus between 10 - 11.6 cM (3.43-4.03 Mbp).** Gene descriptions were obtained from Flybase (FB2019_02) (*58*). Cell in orange are non-coding RNAs. Cells in yellow indicate genes occurring within marker positions flanking the peak at 10.3 cM.

| Chr | Pos start | Pos end | Sense | ID | Name | Alias |
|---|---|---|---|---|---|---|
| 2L | 3404460 | 3426719 | - | FBgn0031530 | Pgant2 | Polypeptide N-Acetylgalactosaminyltransferase 2 |
| 2L | 3445992 | 3447589 | + | FBgn0031533 | CG2772 | FBgn0063009,BcDNA:RE45077 |
| 2L | 3449683 | 3452106 | + | FBgn0031534 | Snx1 | Sorting nexin 1 |
| 2L | 3452466 | 3453755 | + | FBgn0031535 | CG12795 | SO:0000010,SO:0000087,GO:0008270 |
| 2L | 3453697 | 3457064 | - | FBgn0031536 | Cog3 | Cog3 |
| 2L | 3457302 | 3460367 | + | FBgn0266670 | Sec5 | Secretory 5 |
| 2L | 3460322 | 3461952 | - | FBgn0031538 | CG3246 | SO:0000010,SO:0000087,GO:0008289 |
| 2L | 3463290 | 3463400 | - | FBgn0263526 | mir-4971 | mir-4971 stem loop |
| 2L | 3462218 | 3466112 | - | FBgn0005616 | msl-2 | male-specific lethal 2 |
| 2L | 3466157 | 3467334 | + | FBgn0021967 | ND-PDSW | NADH dehydrogenase (ubiquinone) PDSW subunit |
| 2L | 3467342 | 3470004 | - | FBgn0031540 | Pif1 | Pif1 ortholog |
| 2L | 3470182 | 3472364 | + | FBgn0051776 | CG31776 | FBgn0031541,FBgn0063621,CG8845a,CG8845,dpp GalNAcT10 |

| | | | | | | |
|---|---|---|---|---|---|---|
| 2<br>L | 3472<br>603 | 347<br>533<br>8 | + | FBgn00<br>51956 | Pgant4 | Polypeptide N-Acetylgalactosaminyltransferase 4 |
| 2<br>L | 3475<br>365 | 347<br>605<br>8 | + | FBgn02<br>66912 | lncRNA:C<br>R45372 | long non-coding RNA:CR45372 |
| 2<br>L | 3476<br>343 | 347<br>764<br>8 | + | FBgn02<br>64549 | lncRNA:C<br>R43928 | long non-coding RNA:CR43928 |
| 2<br>L | 3478<br>434 | 347<br>961<br>2 | + | FBgn02<br>61560 | Thor | Thor |
| 2<br>L | 3480<br>755 | 349<br>107<br>1 | - | FBgn00<br>31542 | CG15414 | SO:0000010,SO:0000087 |
| 2<br>L | 3504<br>032 | 350<br>418<br>2 | - | FBgn02<br>86737 | snoRNA:<br>Or-CD16 | snoRNA:Orphan CD box 16 |
| 2<br>L | 3493<br>986 | 350<br>811<br>9 | - | FBgn00<br>14396 | tim | timeless |
| 2<br>L | 3508<br>598 | 350<br>903<br>6 | + | FBgn02<br>66837 | lncRNA:C<br>R45299 | long non-coding RNA:CR45299 |
| 2<br>L | 3509<br>099 | 350<br>944<br>0 | + | FBgn02<br>66838 | asRNA:C<br>R45300 | antisense RNA:CR45300 |
| 2<br>L | 3509<br>058 | 351<br>010<br>2 | - | FBgn00<br>51954 | CG31954 | FBgn0031543,SP107,CG3229 |
| 2<br>L | 3510<br>328 | 351<br>465<br>6 | - | FBgn02<br>84253 | LeuRS | Leucyl-tRNA synthetase |
| 2<br>L | 3514<br>872 | 351<br>726<br>4 | + | FBgn00<br>31544 | CG17593 | SO:0000010,SO:0000087,GO:0005737,GO:001250<br>5,GO:0005509,GO:0005783 |
| 2<br>L | 3517<br>273 | 351<br>965<br>1 | - | FBgn00<br>31545 | CG3213 | bs30c03.y1 |
| 2<br>L | 3519<br>677 | 352<br>255<br>5 | + | FBgn00<br>31546 | CG8851 | SO:0000010,SO:0000087 |
| 2<br>L | 3522<br>521 | 352<br>405<br>6 | - | FBgn00<br>31547 | Sr-CIV | Scavenger receptor class C%2C type IV |
| 2<br>L | 3525<br>131 | 352<br>758<br>8 | + | FBgn00<br>31548 | CG8852 | CT9284 |
| 2<br>L | 3527<br>387 | 353<br>031<br>7 | - | FBgn00<br>31549 | Spindly | Spindly |

| | | | | | | |
|---|---|---|---|---|---|---|
| 2 L | 3530 789 | 353 267 8 | + | FBgn00 31550 | IFT57 | Intraflagellar transport 57 |
| 2 L | 3537 844 | 353 833 3 | + | FBgn02 65367 | lncRNA:C R44308 | long non-coding RNA:CR44308 |
| 2 L | 3539 247 | 354 842 3 | + | FBgn00 24244 | drm | drumstick |
| 2 L | 3562 038 | 356 311 8 | + | FBgn02 65368 | lncRNA:C R44309 | long non-coding RNA:CR44309 |
| 2 L | 3563 308 | 356 354 0 | - | FBgn02 65960 | lncRNA:C R44747 | long non-coding RNA:CR44747 |
| 2 L | 3575 350 | 357 580 6 | + | FBgn02 64370 | lncRNA:C R43822 | long non-coding RNA:CR43822 |
| 2 L | 3576 210 | 357 662 3 | + | FBgn02 64363 | CG43815 | SO:0000010,SO:0000087,GO:0008150,GO:000367 4,GO:0005575 |
| 2 L | 3578 013 | 358 076 9 | - | FBgn00 04892 | sob | sister of odd and bowl |
| 2 L | 3587 068 | 358 746 9 | - | FBgn02 84410 | lncRNA:T S2 | long noncoding RNA: testis-specific 2 |
| 2 L | 3588 485 | 359 030 1 | - | FBgn02 64371 | lncRNA:C R43823 | long non-coding RNA:CR43823 |
| 2 L | 3604 224 | 360 675 6 | - | FBgn00 02985 | odd | odd skipped |
| 2 L | 3616 296 | 361 761 9 | + | FBgn02 65369 | lncRNA:C R44310 | long non-coding RNA:CR44310 |
| 2 L | 3619 097 | 362 157 3 | + | FBgn00 15663 | Ugt36A1 | UDP-glycosyltransferase family 36 member A1 |
| 2 L | 3621 110 | 362 170 4 | - | FBgn00 31554 | CG15418 | SO:0000010,SO:0000087,GO:0005615,GO:000486 7 |
| 2 L | 3636 967 | 363 707 3 | - | FBgn02 63523 | mir-4972 | mir-4972 stem loop |
| 2 L | 3622 074 | 365 695 3 | - | FBgn00 00721 | for | foraging |
| 2 L | 3662 745 | 369 211 4 | + | FBgn00 85369 | Drgx | Dorsal root ganglia homeobox |

| | | | | | | |
|---|---|---|---|---|---|---|
| 2 L | 3667977 | 3668551 | - | FBgn0259951 | Sfp24Ba | Seminal fluid protein 24Ba |
| 2 L | 3668691 | 3669134 | - | FBgn0259953 | Sfp24Bd | Seminal fluid protein 24Bd |
| 2 L | 3669261 | 3669775 | - | FBgn0259952 | Sfp24Bb | Seminal fluid protein 24Bb |
| 2 L | 3670131 | 3670581 | - | FBgn0261054 | Sfp24Bc | Seminal fluid protein 24Bc |
| 2 L | 3692123 | 3692587 | - | FBgn0031558 | CG16704 | FBgn0047025,CT37177,BcDNA:RH31195 |
| 2 L | 3692699 | 3693182 | - | FBgn0031559 | CG3513 | SO:0000010,SO:0000087,GO:0004867 |
| 2 L | 3693184 | 3693905 | - | FBgn0051779 | Acp24A4 | Accessory gland protein 24A4 |
| 2 L | 3694105 | 3694418 | - | FBgn0263096 | CR43364 | SO:0000087,SO:0000042 |
| 2 L | 3694431 | 3695020 | - | FBgn0262721 | CG43165 | SO:0000010,SO:0000087,GO:0005615,GO:0004867 |
| 2 L | 3695032 | 3695411 | - | FBgn0031560 | CG16713 | CT37195 |
| 2 L | 3696218 | 3696658 | - | FBgn0031561 | IM33 | Immune induced molecule 33 |
| 2 L | 3697242 | 3697872 | - | FBgn0031562 | CG3604 | FBgn0063121,BcDNA:LP04037 |
| 2 L | 3698451 | 3699092 | - | FBgn0031563 | CG10031 | SO:0000010,SO:0000087,GO:0008150,GO:0004867,GO:0005615 |
| 2 L | 3699464 | 3699826 | + | FBgn0259954 | CG42464 | SO:0000010,SO:0000087,GO:0004867 |
| 2 L | 3700195 | 3700658 | + | FBgn0259955 | CG42465 | SO:0000010,SO:0000087,GO:0004867 |
| 2 L | 3700674 | 3701092 | + | FBgn0259956 | Sfp24C1 | Seminal fluid protein 24C1 |
| 2 L | 3701290 | 3701699 | + | FBgn0259957 | CG42467 | SO:0000010,SO:0000087,GO:0004867 |

| | | | | | | |
|---|---|---|---|---|---|---|
| 2 L | 3701826 | 3702248 | + | FBgn0262621 | CG43145 | lincRNA.26 |
| 2 L | 3702614 | 3702947 | + | FBgn0265370 | lncRNA:CR44311 | long non-coding RNA:CR44311 |
| 2 L | 3703659 | 3704253 | + | FBgn0031564 | CG2816 | SO:0000010,SO:0000087,GO:0004867 |
| 2 L | 3704106 | 3705488 | - | FBgn0051778 | CG31778 | FBgn0031565,FBgn0047241,FBgn0063619,BcDNA:HL03084 |
| 2 L | 3706551 | 3707063 | - | FBgn0051777 | CG31777 | FBgn0031565,FBgn0063620,BEST:RH19272 |
| 2 L | 3707230 | 3711748 | - | FBgn0015376 | cutlet | cutlet |
| 2 L | 3711881 | 3713105 | - | FBgn0051955 | CG31955 | FBgn0031565 |
| 2 L | 3713360 | 3716810 | + | FBgn0031566 | CG2818 | SO:0000010,SO:0000087,GO:2001070,GO:0046475,GO:0047389 |
| 2 L | 3716800 | 3717773 | - | FBgn0040104 | lectin-24A | lectin-24A |
| 2 L | 3718185 | 3729145 | + | FBgn0003386 | Shaw | Shaker cognate w |
| 2 L | 3730466 | 3754617 | + | FBgn0031568 | CG10019 | FBgn0043805,GH25970,BEST:GH25970 |
| 2 L | 3757256 | 3769525 | + | FBgn0263846 | CG43707 | CG31959,FBgn0051959,FBgn0031569,CG31772,FBgn0051772,CG31772,CG31959,CG10020 |
| 2 L | 3765494 | 3765863 | + | FBgn0263842 | CG43703 | SO:0000010,SO:0000087,GO:0004867 |
| 2 L | 3767621 | 3767688 | + | FBgn0262392 | mir-1004 | mir-1004 stem loop |
| 2 L | 3769926 | 3770454 | - | FBgn0263843 | lncRNA:CR43704 | long non-coding RNA:CR43704 |
| 2 L | 3771700 | 3784121 | + | FBgn0004893 | bowl | brother of odd with entrails limited |
| 2 L | 3784947 | 3785622 | + | FBgn0051960 | CG31960 | FBgn0031570,CG10022,CBP1 |

| | | | | | | |
|---|---|---|---|---|---|---|
| 2 L | 3785738 | 3786343 | + | FBgn0051958 | CR31958 | FBgn0031570,CG10022,CG31958 |
| 2 L | 3787167 | 3802726 | + | FBgn0031571 | bark | bark beetle |
| 2 L | 3803545 | 3810671 | + | FBgn0021800 | Reph | Regulator of eph expression |
| 2 L | 3810801 | 3813410 | - | FBgn0031573 | CG3407 | cg3407 |
| 2 L | 3815438 | 3815948 | + | FBgn0264885 | lncRNA:CR44076 | long non-coding RNA:CR44076 |
| 2 L | 3817724 | 3818534 | + | FBgn0264886 | lncRNA:CR44077 | long non-coding RNA:CR44077 |
| 2 L | 3825675 | 3827099 | + | FBgn0003430 | slp1 | sloppy paired 1 |
| 2 L | 3836840 | 3839185 | + | FBgn0004567 | slp2 | sloppy paired 2 |
| 2 L | 3845130 | 3845716 | - | FBgn0266318 | lncRNA:CR44983 | long non-coding RNA:CR44983 |
| 2 L | 3851951 | 3852604 | + | FBgn0267007 | lncRNA:CR45451 | long non-coding RNA:CR45451 |
| 2 L | 3854288 | 3854994 | - | FBgn0267005 | lncRNA:CR45449 | long non-coding RNA:CR45449 |
| 2 L | 3856478 | 3857172 | - | FBgn0267006 | lncRNA:CR45450 | long non-coding RNA:CR45450 |
| 2 L | 3861405 | 3862097 | - | FBgn0264887 | lncRNA:CR44078 | long non-coding RNA:CR44078 |
| 2 L | 3862668 | 3867507 | + | FBgn0031574 | TTLL4B | Tubulin tyrosine ligase-like 4B |
| 2 L | 3867697 | 3871425 | + | FBgn0031575 | Cep97 | Centrosomal protein 97kDa |
| 2 L | 3871271 | 3872458 | - | FBgn0051957 | CG31957 | SO:0000010,SO:0000087,GO:0006413,GO:0003743 |
| 2 L | 3872646 | 3902860 | - | FBgn0000256 | capu | cappuccino |

| | | | | | | |
|---|---|---|---|---|---|---|
| 2L | 3911796 | 3914288 | - | FBgn0051773 | CG31773 | FBgn0031577,CG3390 |
| 2L | 3902928 | 3991713 | - | FBgn0051774 | fred | friend of echinoid |
| 2L | 4000018 | 4000806 | + | FBgn0264341 | CG43797 | SO:0000010,SO:0000087 |
| 2L | 4001084 | 4001620 | + | FBgn0266883 | lncRNA:CR45344 | long non-coding RNA:CR45344 |
| 2L | 4004080 | 4004480 | + | FBgn0085205 | CG34176 | SO:0000010 |
| 2L | 4004984 | 4005670 | + | FBgn0031579 | CG15422 | SO:0000010,SO:0000087 |
| 2L | 4006041 | 4006606 | + | FBgn0031580 | CG15423 | SO:0000010,SO:0000087 |
| 2L | 4007690 | 4007984 | + | FBgn0264295 | CG43773 | FBgn0031581,CG10039 |
| 2L | 4008653 | 4009090 | + | FBgn0264296 | CG43774 | FBgn0031581,CG10039 |
| 2L | 4031377 | 4115749 | + | FBgn0000547 | ed | echinoid |

**Table S2. Sanger sequencing calls for the three upstream promoter indels in 171 DGRP lines.** The indels were sequenced in the order: c.-439_-433del, c.-334_-333insACATTCAT and 21bp indel (c.-171_-151del). When a heterozygous indel was encountered, indicating that the line was not fully inbred, the remaining indels were not sequenced and, therefore, indicated with NA. I: insertion allele and D: deletion allele. The c.-334_-333insACATTCAT call from DGRP Freeze2.0 (*42, 52*) and a SNP that occurs within the 21bp insertion are also indicated.

| Line | resistance 486 | c.-439_-433del | c.-334_-333insACATTCAT | c.-171_-151del | comment | c.-334_-333insACATTCAT dgrp2 call | snp at pos 3 in c.-171_-151del |
|---|---|---|---|---|---|---|---|
| 21 | 0.08 | I | I | NA | NA | I | NA |
| 26 | 0.02 | D | D | NA | NA | NA | NA |
| 31 | 0.14 | I | I | NA | NA | I | NA |
| 32 | 0.03 | I | D | NA | NA | D | NA |
| 38 | 0.01 | I | I | I | NA | I | G |

| | | | | | | | |
|---|---|---|---|---|---|---|---|
| 40 | 0.55 | I | I | I | NA | I | A |
| 41 | NA | I | I | NA | NA | I | NA |
| 42 | 0.03 | I | I | I | NA | I | G |
| 45 | 0.03 | I | I | I | NA | I | A |
| 48 | 0.03 | I | D | I | NA | D | G |
| 57 | 0.02 | I | I | I | NA | I | G |
| 59 | 0.06 | I | I | I | NA | I | G |
| 69 | 0.02 | D | D | D | NA | D | NA |
| 73 | 0.14 | D | D | D | NA | D | NA |
| 75 | 0.04 | I | I | I | NA | I | G |
| 85 | 0.06 | I | NA | NA | het 8bp | NA | NA |
| 88 | 0.14 | NA | NA | NA | het 7bp | NA | NA |
| 91 | 0.08 | I | I | I | NA | I | G |
| 93 | 0 | I | D | I | NA | D | G |
| 100 | 0.05 | NA | NA | NA | het 7bp | NA | NA |
| 101 | 0.04 | I | NA | NA | het 8bp | NA | NA |
| 105 | 0.12 | D | D | D | NA | D | NA |
| 109 | 0 | I | I | I | NA | I | G |
| 129 | 0.04 | I | I | I | NA | I | A |
| 136 | 0.01 | I | I | I | NA | I | G |
| 138 | 0.03 | D | D | D | NA | D | NA |
| 142 | 0.04 | I | I | I | NA | I | G |
| 153 | 0.01 | I | I | I | NA | I | G |
| 161 | 0.05 | I | D | I | NA | D | G |
| 176 | 0.1 | I | I | I | NA | I | A |
| 177 | 0.01 | I | I | I | NA | I | G |
| 181 | 0.03 | D | D | D | NA | D | NA |
| 189 | 0.01 | I | I | I | NA | NA | A |
| 195 | 0.03 | I | I | I | NA | I | A |
| 208 | 0.05 | D | D | D | NA | D | NA |
| 217 | 0.02 | I | I | I | NA | I | A |
| 228 | 0.02 | D | D | D | NA | D | NA |
| 229 | 0.05 | I | I | I | NA | I | A |
| 235 | 0.15 | I | I | I | NA | I | A |
| 237 | 0.01 | I | NA | NA | het 8bp | NA | NA |
| 256 | 0.09 | I | I | I | NA | I | G |
| 280 | 0.21 | I | I | I | NA | I | G |
| 287 | 0.54 | I | I | I | NA | I | G |
| 301 | 0.12 | NA | NA | NA | het 7bp | NA | NA |
| 303 | 0.08 | I | I | I | NA | I | G |
| 304 | 0 | I | I | I | NA | I | G |
| 306 | 0.11 | I | I | I | NA | I | A |

| | | | | | | | |
|---|---|---|---|---|---|---|---|
| 309 | 0.01 | I | I | I | NA | I | G |
| 313 | 0.03 | I | I | I | NA | I | A |
| 317 | 0.23 | I | I | I | NA | I | G |
| 318 | 0.07 | I | I | I | NA | I | G |
| 319 | 0.03 | I | I | I | NA | I | A |
| 320 | 0.08 | I | I | I | NA | I | A |
| 321 | 0.06 | I | I | I | NA | I | A |
| 324 | 0.19 | I | I | I | NA | I | G |
| 332 | 0.04 | I | I | I | NA | I | A |
| 336 | 0.11 | I | I | I | NA | I | G |
| 338 | 0.15 | I | NA | NA | het 8bp | NA | NA |
| 340 | 0.01 | I | I | I | NA | I | G |
| 348 | 0.15 | D | D | NA | NA | D | NA |
| 350 | 0.1 | D | D | I | NA | D | G |
| 352 | 0.09 | I | NA | NA | het 8bp | NA | NA |
| 354 | 0.01 | I | D | I | NA | D | G |
| 355 | 0.04 | I | D | I | NA | D | G |
| 356 | 0.07 | I | I | I | NA | I | A |
| 357 | 0.02 | I | I | I | NA | I | A |
| 358 | 0.06 | D | D | I | NA | D | G |
| 360 | 0 | I | I | I | NA | I | A |
| 361 | 0.06 | I | I | I | NA | NA | A |
| 362 | 0.05 | I | D | I | NA | D | G |
| 365 | 0.28 | I | I | I | NA | I | G |
| 370 | 0.01 | I | I | I | NA | I | A |
| 371 | NA | I | I | I | NA | I | A |
| 373 | 0 | D | I | I | NA | I | G |
| 374 | 0.1 | I | I | I | NA | I | G |
| 375 | 0.11 | D | D | D | NA | NA | NA |
| 377 | NA | I | NA | NA | het 8bp | NA | NA |
| 379 | 0.1 | I | I | I | NA | I | G |
| 380 | 0.09 | I | I | I | NA | I | G |
| 381 | 0.01 | NA | D | I | NA | D | G |
| 382 | 0.07 | I | I | I | NA | I | G |
| 385 | 0.08 | I | I | I | NA | I | A |
| 386 | 0.03 | D | D | I | NA | NA | G |
| 390 | 0.03 | I | D | I | NA | D | G |
| 391 | 0.21 | I | I | I | NA | I | G |
| 392 | 0.02 | I | I | I | NA | I | A |
| 395 | 0.03 | I | I | I | NA | I | A |
| 397 | 0.07 | I | I | I | NA | I | A |
| 399 | 0.15 | I | I | I | NA | I | A |

| | | | | | | | |
|---|---|---|---|---|---|---|---|
| 405 | 0.02 | I | NA | NA | het 8bp | NA | NA |
| 406 | 0.02 | D | D | I | NA | D | G |
| 409 | 0.04 | D | I | I | NA | I | G |
| 426 | 0.09 | NA | NA | NA | het 7bp | NA | NA |
| 427 | 0 | I | I | I | NA | I | A |
| 437 | 0.39 | I | I | I | NA | I | A |
| 439 | 0.15 | I | I | I | NA | I | G |
| 440 | 0.12 | I | I | I | NA | I | G |
| 441 | 0.02 | I | I | I | NA | I | G |
| 443 | 0 | D | D | I | NA | NA | G |
| 461 | 0.02 | D | I | I | NA | I | G |
| 486 | 0.08 | D | I | I | NA | I | G |
| 491 | 0.02 | I | I | I | NA | I | G |
| 492 | 0.04 | D | D | I | NA | D | G |
| 502 | 0.08 | I | NA | I | het 8bp | NA | G |
| 508 | 0.02 | I | I | I | NA | I | G |
| 509 | 0.05 | D | I | I | NA | I | A |
| 517 | 0.13 | I | D | I | NA | D | G |
| 528 | 0.16 | I | NA | NA | het 8bp | NA | NA |
| 530 | 0.31 | I | I | I | NA | I | G |
| 531 | 0.05 | I | I | I | NA | I | G |
| 535 | 0.05 | I | I | I | NA | I | A |
| 551 | 0.11 | I | I | I | NA | I | G |
| 555 | 0.08 | I | I | I | NA | I | G |
| 559 | 0.14 | D | D | D | NA | D | NA |
| 563 | 0.21 | I | I | I | NA | I | A |
| 566 | 0.48 | I | I | I | NA | I | A |
| 584 | 0.14 | I | D | I | NA | NA | G |
| 589 | 0.48 | I | I | I | NA | I | G |
| 595 | 0 | I | D | I | NA | D | G |
| 596 | 0.03 | I | I | I | NA | I | A |
| 627 | 0.07 | D | D | I | NA | D | G |
| 703 | 0.03 | I | I | I | NA | I | A |
| 705 | 0.08 | I | I | I | NA | I | G |
| 712 | 0.04 | I | D | I | NA | D | G |
| 714 | 0.01 | D | I | I | NA | I | G |
| 716 | 0.05 | I | I | I | NA | I | G |
| 721 | 0.03 | I | I | I | NA | I | G |
| 732 | 0.06 | I | I | I | NA | I | G |
| 737 | 0.04 | I | I | I | NA | I | G |
| 738 | NA | I | I | I | NA | I | G |
| 748 | 0.07 | D | D | I | NA | D | G |

| | | | | | | | |
|---|---|---|---|---|---|---|---|
| 757 | NA | I | I | I | NA | I | G |
| 774 | 0.11 | I | I | I | NA | I | G |
| 776 | 0.06 | I | I | I | NA | I | G |
| 783 | 0.15 | I | I | I | NA | I | G |
| 786 | 0.03 | I | I | I | NA | I | G |
| 787 | 0.07 | I | I | I | NA | I | A |
| 790 | 0.15 | I | I | I | NA | I | G |
| 796 | 0.07 | I | D | I | NA | NA | G |
| 802 | 0.05 | I | I | I | NA | I | A |
| 804 | 0.06 | I | I | I | NA | I | G |
| 808 | 0.05 | I | I | I | NA | I | G |
| 810 | 0.24 | I | I | I | NA | I | G |
| 812 | 0.27 | I | NA | NA | het 8bp | NA | NA |
| 818 | 0.07 | I | I | I | NA | I | A |
| 819 | 0.05 | I | I | I | NA | I | A |
| 820 | 0.29 | D | I | I | NA | I | G |
| 821 | 0.07 | I | I | I | NA | I | A |
| 822 | 0.11 | I | D | I | NA | NA | G |
| 832 | 0.78 | I | D | I | NA | D | G |
| 843 | 0.18 | I | I | I | NA | I | A |
| 849 | 0.14 | I | I | I | NA | NA | A |
| 850 | 0.11 | I | I | I | NA | I | A |
| 852 | 0.17 | I | I | I | NA | I | A |
| 853 | 0.06 | I | I | I | NA | I | A |
| 855 | 0.04 | I | I | I | NA | I | G |
| 857 | 0.02 | I | I | I | NA | I | G |
| 859 | 0.13 | I | I | I | NA | I | A |
| 861 | 0.06 | I | I | I | NA | I | A |
| 879 | 0.07 | I | I | I | NA | I | A |
| 882 | 0.02 | I | I | I | NA | I | A |
| 884 | 0.11 | I | I | I | NA | I | G |
| 890 | 0.06 | I | I | I | NA | I | A |
| 892 | 0.03 | D | D | D | NA | D | NA |
| 894 | 0.03 | I | D | I | NA | D | G |
| 897 | 0.31 | I | I | I | NA | I | G |
| 900 | 0.03 | I | I | I | NA | I | G |
| 907 | 0.08 | I | I | I | NA | I | G |
| 908 | 0.06 | I | I | I | NA | I | A |
| 911 | 0.04 | D | I | I | NA | I | G |
| 913 | 0.11 | I | I | I | NA | I | G |

**Table S3. Primers to amplify indel and SNP primers for quantitative trait locus (QTL) mapping of the second chromosome of *Drosophila melanogaster*.** Genetic and physical positions were obtained from version 6 of *D. melanogaster* genome from Flybase (FB2019_02) (*58*). As genetic positions were only identified to the closest integer on Flybase, positions in between were assigned based on their respective physical location. Indel size variation and number of SNPs differentiating DGRP-437 and DGRP-892 were based on DGRP Freeze 2.0 (*42, 52*).

| Location (Chr-cM) | Forward primer | Reverse primer | Forward primer start (bp) | Reverse primer start (bp) | Type | 437size (bp) | 892 size (bp) | # SNP | Cross |
|---|---|---|---|---|---|---|---|---|---|
| 2L-3 | CATTACACTTGCAGCCGGAA | GGTGTACCGAATGAGATTGCG | 2007685 | 2008065 | Indel | 380 | 325 | 8 | larvae |
| 2L-5 | ACACCAAATACGCCAAGCAA | AGGCTGAGGTATGCGTTCTT | 2243388 | 2243634 | Indel | 247 | 292 | 0 | larvae |
| 2L-7 | CGCAATCAGAGCCAGCATTT | CCAACCGGTCCAAAGTTCAG | 2804176 | 2804735 | Indel | 524 | 523 | 8 | larvae, adult |
| 2L-10.3 | CGTCGATGGCAATATCTCCG | CCTATCGCCACTGTATCCCA | 3522926 | 3523290 | Indel | 364 | 304 | 6 | larvae, adult |
| 2L-12 | CTGGACAGCTGGATCTCGAT | TCGCTCTGTCGCTCTGTTAT | 4250761 | 4251130 | Indel | 369 | 416 | 5 | larvae, adult |
| 2L-17 | ATGGTGGAGGCGCTTAAGT | ACTCTGTCACTGTTGCTGTTG | 5511730 | 5512284 | Indel | 490 | 545 | 4 | larvae, adult |
| 2L-27 | TTCCACGGTAGCAAAATGGC | CCCCGTCTCCCTCTTTCTTT | 7564467 | 7564772 | Indel | 305 | 195 | 0 | larvae |
| 2L-34 | AGGATTTGAGGGGAGTGCAA | GAGCAGAGGTAGAGGCAGAG | 8762316 | 8762746 | Indel | 430 | 484 | 8 | larvae |
| 2L-54 | AATTGCACACGGAGGACAAC | GCCAATTAGAGCTCCACTGC | 19518131 | 19518468 | Indel | 416 | 337 | 3 | larvae |
| 2R-67 | TTTGATGGATGAGGAGCCGT | CTGATGCAGTGTTTACCCCG | 12748432 | 12748868 | Indel | 479 | 436 | 20 | larvae |
| 2R-104 | ATCGACTCAAGTGGCTGTCA | GCACGATCAAGTTTCACCGA | 23666654 | 23667009 | Indel | 355 | 406 | 4 | larvae |
| 2L-8 | TGTCGTGGTTTTACAAGCCG | TGGAGACGCTCGATCTAACC | 3056629 | 3057129 | SNP | 501 | 500 | 10 | adult |

| | | | | | | | | | |
|---|---|---|---|---|---|---|---|---|---|
| 2L-8.5 | GCTCTGGTTG GGTAGGGAG | AGTGCGGAT GGTCAGTGT AA | 321428 0 | 3214818 | S N P | 538 | 538 | 6 | adult |
| 2L-9 | GCAGTATTCG CGGCATTTTG | TGAATTCGCA GCGTTGAAC A | 332511 7 | 3325661 | S N P | 544 | 544 | 1 3 | adult |
| 2L-10 | TGTGGGGAC GGGGATTTAT T | GTAAGCGTG AGAGGGAGA CA | 342561 0 | 3426126 | S N P | 516 | 516 | 8 | adult |
| 2L-10.7 | GCCAGGATTT GAGAAGATA GGC | TCTTCATTTC GCCACTTAGC T | 371661 8 | 3717252 | S N P | 634 | 634 | 5 | adult |
| 2L-11 | TCAGTACCAC CGCTCCAAAT | TCGTTTCGGC AATAGTTAAT GGT | 380211 6 | 3802665 | S N P | 549 | 549 | 1 | adult |
| 2L-11.3 | GTAAATTCCG CGACTGGGT C | AAAATGTTG GTAGGGGCA GC | 393000 5 | 3930570 | S N P | 565 | 565 | 1 7 | adult |
| 2L-11.6 | GAGGAGAGC AGCAACGAC TA | CCAATTGACA GGAGCCAAC A | 403272 9 | 4033264 | S N P | 535 | 535 | 1 6 | adult |

**Table S4. Primers used for generating pCFD5-w LectinKO for integrating two guide RNAs targetting *Lectin-24A*.** Sequence in bold refereces to the insert and nonbold is the backbone vector.

| Name | Sequence |
|---|---|
| Lectin24aKO_PCR1_Fw | GCGGCCCGGGTTCGATTCCCGGCCGATGCA**GGTCTCCAAAGATTCATGCA**GTTTTAGAGCTAGAAATAGCAAG |
| Lectin24aKO_PCR1_Rv | **GAAGCCAGTTATGGAGTATG**TGCACCAGCCGGGAATCGAACCC |
| Lectin24aKO_PCR2_Fw | **CATACTCCATAACTGGCTTC**GTTTTAGAGCTAGAAATAGCAAG |
| Lectin24aKO_PCR2_Rv | **TGCATGAATCTTTGGAGACC**TGCACCAGCCGGGAATCGAACCC |
| Lectin24aKO_PCR3_Fw | **GGTCTCCAAAGATTCATGCA**GTTTTAGAGCTAGAAATAGCAAG |
| Lectin24aKO_PCR3_Rv | ATTTTAACTTGCTATTTCTAGCTCTAAAAC**GAAGCCAGTTATGGAGTATG**TGCACCAGCCGGGAATCGAACCC |
| U63seqfwd | ACGTTTTATAACTTATGCCCCTAAG |
| pCFDseqrev | GCACAATTGTCTAGAATGCATAC |

**Table S5. Primers used to generate fluorescent reporter constructs carrying different alleles for the 3 indels and 6 SNPs polymorphic between DGRP-892 and DGRP-437 occurring in the 1,000bp region upstream of *Lectin-24A*.**

| Primer | Sequence |
|---|---|

| | |
|---|---|
| Lec500bppromF | GCTATACGAAGTTATCTGCAGGAATCCGAACAAACCGAACC |
| Lec500bppromR | CCTTGCTCACCATGATTTTATTTATTTAGATGCGTTTAATATCAACAGCTTTG |
| venusLecPromF | TAAAATCATGGTGAGCAAGGGCGAGGAG |
| venusLecPromR | ATTATAAGCTGCAATAAACAAGTTTTACTTGTACAGCTCGTCCATGCC |
| lecpromD21frag1R | TACTTAAAAAACTATTCAGTAAAAAATAGTTGCTATGTATACAATATTGTGAAATATT |
| lecpromD21frag2F | TTTTTACTGAATAGTTTTTTAAGTAGAAACACAAGACTGAAGAAAATAAAGCAATCAAG |
| 1_2insmutfrag1R | GCACATTGGCGCCATACTCAAGTGAATGTCAGAGC |
| 1_2insmutfrag2F | GGCGCCAATGTGCAACACTGCGTATGAGTGATG |
| snpmutfrag1R | CTTAAATATTATCAAAACGAATTAAAAGTAATTTTTTAAATAATTCTTTAAATATGAAATTTG |
| Snpmutfrag2F | CTTTTAATTCGTTTTGATAATATTTAAGTAATTCACAATATTGTATACATAGC |
| lecPrdelta7Rv | CTGCGCATGCACTCGATTCTGAGATGAAAAACAAATC |
| LecPrdelta8Fw | ATCTCAGAATCGAGTGCATGCGCAGTGTCAC |
| LecPrdelta8Rv | ACATCACTCATACGCAGTGTTGCACATTGGC |
| LecPr21bpFw | GTGCAACACTGCGTATGAGTGATGTAGAAAGC |
| LecPr2snpF | CATTCATACTTGAGTATGGCGCCAATGTGC |
| LecPr2snpRv | TTGGCGCCATACTCAAGTATGAATGTGAATGTCAGAG |
| Truedelta8Fw | GATGTTGATTTGTTTTTCATCTCAGAATCGAGTG |
| truedelta8Rv | TTGGCGCCATACTTGAGTGAATGTCAGAGCAAAAG |
| 21bp2snpFw | GACATTCACTCAAGTATGGCGCCAATGTGC |
| 7bptrue8Rv | CTGAGATGAAAAACAAATCAACATCCCTCCAC |

**Table S6. *Lectin-24A* promoter (LP) constructs carrying different alleles for the 3 indels and 6 SNPs polymorphic between DGRP-892 and DGRP-437 occurring in the 500bp region upstream of *Lectin-24A*.** Δ refers to the removal of the insertion allele from the DGRP-437 *Lectin-24A* upstream sequence (437LP). For SNPs, DGRP-892 alleles were substituted with the DGRP-437 alleles in 437LP.

| Construct | Primers used | Template |
|---|---|---|
| 437LP | Lec500bppromF/Lec500bppromR | DGRP-437 |
| | venusLecPromF/venusLecPromR | venus |
| 892LP | Lec500bppromF/Lec500bppromR | DGRP-892 |
| | venusLecPromF/venusLecPromR | venus |
| Δc.-171_-151del | Lec500bppromF/lecpromD21frag1R | DGRP-437 |
| | lecpromD21frag2F/lec500bppromR | DGRP-437 |
| | venusLecPromF/venusLecPromR | venus |
| Δc.-439_-433del, Δ333insACATTCAT | Lec500bppromF/1_2insmutfrag1R | DGRP-892 |
| | 1_2insmutfrag2F/lec500bppromR | DGRP-437 |
| | venusLecPromF/venusLecPromR | venus |
| 3717919_SNP, 3717922_SNP, | Lec500bppromF/snpmutfrag1R | DGRP-437 |
| | Snpmutfrag2F/lec500promR | DGRP-437 |
| | venusLecPromF/venusLecPromR | venus |

| | | |
|---|---|---|
| 3717950_SNP, 3717956_SNP | | |
| Δc.-439_-433del | Lec500promF/lecPrdelta7Rv | DGRP-892 |
| | LecPrdelta8Fw/Lec500PromR | DGRP-437 |
| | venusLecPromF/venusLecPromR | venus |
| Δ333insACATTCAT, 3718036_SNP, 3718037_SNP | LecPrdelta8Fw/LecPrdelta8Rv | DGRP-892 |
| | LecPr21bpFw/Lec500PromR | DGRP-437 |
| | Lec500PromF/LecPrdelta7Rv | DGRP-437 |
| | venusLecPromF/venusLecPromR | venus |
| 3718036_SNP, 3718037_SNP | LecPr2snpF/LecProm500Rv | DGRP-437 |
| | LecProm500Fw/LecPr2snpRv | DGRP-437 |
| | venuslecPromF/venusLecPromR | venus |
| Δ333insACATTCAT | LecProm500Fw/7bpRvtrue8 | DGRP-437 |
| | Truedelta8Fw/truedelta8Rv | DGRP-892 |
| | 21bp2snpFw/LecProm500Rv | DGRP-437 |
| | venuslecPromF/venusLecPromR | venus |

**Table S7. SNPs segregating in the 21 DGRP lines used for testing *Lectin-24A* allele expression.** Calls for eight SNPs in the amplified region between 3717069 and 3717568 on chromosome 2L were obtained from DGRP Freeze2.0 (*42, 52*). 2L:3717102 was the polymorphic site used for estimating allele-specific expression.

| **Line** | **2L_3717102** | **2L_3717224** | **2L_3717262** | **2L_3717291** | **2L_3717309** | **2L_3717487** | **2L_3717503** | **2L_3717530** |
|---|---|---|---|---|---|---|---|---|
| Ref allele | C | A | T | A | A | A | T | C |
| Alt allele | T | T | C | G | G | T | C | T |
| 105 | 0 | 0 | 0 | 0 | 0 | 0 | 0 | 0 |
| 136 | 0 | 0 | 0 | 0 | 2 | 0 | 0 | 2 |
| 161 | 0 | 0 | 0 | 0 | 2 | 0 | 0 | 2 |
| 208 | 0 | 0 | 0 | 0 | 0 | 0 | 0 | 0 |
| 217 | 0 | 0 | 0 | 0 | 2 | 0 | 0 | 2 |
| 228 | 0 | 0 | 0 | 0 | 0 | 0 | 0 | 0 |
| 280 | 0 | 0 | 0 | 0 | 2 | 0 | 0 | 2 |
| 350 | 0 | 0 | 0 | 0 | 2 | 2 | 0 | 2 |
| 386 | 0 | 0 | 0 | 0 | 2 | 2 | 0 | 2 |
| 406 | 0 | 0 | 0 | 0 | 2 | 2 | 0 | 2 |
| 409 | 0 | 0 | 0 | 0 | 2 | 0 | 0 | 2 |
| 427 | 0 | 0 | 0 | 0 | 2 | 0 | 0 | 2 |
| 437 | 2 | 0 | 0 | 0 | 2 | 0 | 0 | 2 |
| 486 | 0 | 0 | 0 | 0 | 2 | 0 | 2 | 2 |
| 509 | 0 | 0 | 0 | 0 | 0 | 0 | 0 | 2 |
| 517 | 0 | 2 | 0 | 0 | 2 | 0 | 0 | 2 |

| | | | | | | | | |
|---|---|---|---|---|---|---|---|---|
| 584 | 0 | 0 | 0 | 0 | 2 | 0 | 0 | 2 |
| 627 | 0 | 0 | 0 | 0 | 2 | 2 | 0 | 2 |
| 787 | 0 | 0 | 0 | 0 | 2 | 0 | 0 | 2 |
| 820 | 0 | 0 | 0 | 0 | 2 | 0 | 0 | 2 |
| 822 | 0 | 0 | 2 | 2 | 2 | 0 | 2 | 2 |
| 892 | 0 | 0 | 0 | 0 | 0 | 0 | 0 | 0 |

**Table S8. Comparison of *Lectin-24A* variant calls using the method used in this paper versus those made by DGRP Freeze2.0 (*42, 52*), Global Diversity Lines (*77*) and Sanger sequencing.** Green cells indicate a match between datasets being compared and orange cells indicate a mismatch.

| DGRP Freeze2.0 - SNPs | | | |
|---|---|---|---|
| Variant | Allele_DGN | Reference_allele_DGRP_Freeze2.0 | Alternate_allele_DGRP_Freeze2.0 |
| p.Arg275Ser | Reference | 130 | 0 |
| p.Arg275Ser | Alternate | 0 | 16 |
| p.Tyr266His | Reference | 111 | 0 |
| p.Tyr266His | Alternate | 0 | 40 |
| p.Arg262Gly | Reference | 133 | 0 |
| p.Arg262Gly | Alternate | 0 | 18 |
| p.Ser244Gly | Reference | 140 | 0 |
| p.Ser244Gly | Alternate | 0 | 12 |
| p.Cys238Gly | Reference | 143 | 0 |
| p.Cys238Gly | Alternate | 0 | 6 |
| p.Asp228Glu | Reference | 144 | 0 |
| p.Asp228Glu | Alternate | 0 | 7 |
| p.Glu208Glu | Reference | 150 | 0 |
| p.Glu208Glu | Alternate | 0 | 0 |
| p.Tyr140Tyr | Reference | 14 | 0 |
| p.Tyr140Tyr | Alternate | 0 | 136 |
| p.Asp84Val | Reference | 151 | 0 |
| p.Asp84Val | Alternate | 0 | 0 |
| p.Leu81* | Reference | 142 | 0 |
| p.Leu81* | Alternate | 0 | 9 |
| p.Val76Ile | Reference | 141 | 0 |
| p.Val76Ile | Alternate | 0 | 10 |
| p.Thr67Ala | Reference | 9 | 0 |
| p.Thr67Ala | Alternate | 0 | 143 |
| p.Ser40Pro | Reference | 7 | 0 |
| p.Ser40Pro | Alternate | 0 | 144 |
| p.Val6Val | Reference | 151 | 0 |

| | | | |
|---|---|---|---|
| p.Val6Val | Alternate | 0 | 1 |
| c.-4995G>T | Reference | 148 | 0 |
| c.-4995G>T | Alternate | 0 | 3 |
| c.-153A>C | Reference | 102 | 0 |
| c.-153A>C | Alternate | 0 | 18 |
| c.-161T>A | Reference | 124 | 0 |
| c.-161T>A | Alternate | 0 | 16 |
| c.-212A>T | Reference | 13 | 0 |
| c.-212A>T | Alternate | 0 | 132 |
| c.-215G>A | Reference | 13 | 0 |
| c.-215G>A | Alternate | 0 | 132 |
| c.-234T>A | Reference | 102 | 0 |
| c.-234T>A | Alternate | 0 | 34 |
| c.-243A>T | Reference | 11 | 0 |
| c.-243A>T | Alternate | 0 | 133 |
| c.-323G>T | Reference | 148 | 0 |
| c.-323G>T | Alternate | 0 | 3 |
| c.-342C>T | Reference | 121 | 0 |
| c.-342C>T | Alternate | 0 | 24 |
| c.-366T>C | Reference | 149 | 0 |
| c.-366T>C | Alternate | 0 | 3 |
| **DGRP Sanger - Indels** | | | |
| **Upstream_Indel** | **Allele_DGN** | **Ancestral_allele_Sanger** | **Derived_allele_Sanger** |
| c.-439_-433del | Insertion | 101 | 0 |
| c.-439_-433del | Deletion | 0 | 22 |
| c.-334_-333insACATTCAT | Deletion | 27 | 0 |
| c.-334_-333insACATTCAT | Insertion | 0 | 97 |
| c.-171_-151del | Insertion | 100 | 0 |
| c.-171_-151del | Deletion | 0 | 10 |
| **GDL - SNPs** | | | |
| **Variant** | **Allele_DGN** | **Reference_allele_GDL** | **Alternate_allele_GDL** |
| c.*67T>A | Reference | 7 | 0 |
| c.*67T>A | Alternate | 0 | 49 |
| c.*66C>T | Reference | 52 | 0 |
| c.*66C>T | Alternate | 0 | 4 |
| c.*44G>T | Reference | 6 | 0 |
| c.*44G>T | Alternate | 0 | 49 |

| | | | |
|---|---|---|---|
| c.*40T>C | Reference | 50 | 0 |
| c.*40T>C | Alternate | 0 | 6 |
| p.Arg275Ser | Reference | 48 | 0 |
| p.Arg275Ser | Alternate | 0 | 8 |
| p.Tyr266His | Reference | 37 | 0 |
| p.Tyr266His | Alternate | 0 | 18 |
| p.Arg262Gly | Reference | 45 | 0 |
| p.Arg262Gly | Alternate | 0 | 10 |
| p.Gln254* | Reference | 50 | 0 |
| p.Gln254* | Alternate | 0 | 5 |
| p.Ser244Gly | Reference | 46 | 0 |
| p.Ser244Gly | Alternate | 0 | 8 |
| p.Cys238Gly | Reference | 49 | 0 |
| p.Cys238Gly | Alternate | 0 | 6 |
| p.Asp228Glu | Reference | 49 | 0 |
| p.Asp228Glu | Alternate | 0 | 6 |
| p.Glu208Glu | Reference | 49 | 0 |
| p.Glu208Glu | Alternate | 0 | 7 |
| p.Glu202* | Reference | 53 | 0 |
| p.Glu202* | Alternate | 0 | 3 |
| p.Leu158Val | Reference | 50 | 0 |
| p.Leu158Val | Alternate | 0 | 6 |
| p.Val155Ile | Reference | 50 | 0 |
| p.Val155Ile | Alternate | 0 | 6 |
| p.Asn153Ile | Reference | 50 | 0 |
| p.Asn153Ile | Alternate | 0 | 6 |
| p.Ile152Ile | Reference | 50 | 0 |
| p.Ile152Ile | Alternate | 0 | 6 |
| p.Met141Lys | Reference | 50 | 0 |
| p.Met141Lys | Alternate | 0 | 6 |
| p.Tyr140Tyr | Reference | 13 | 0 |
| p.Tyr140Tyr | Alternate | 0 | 43 |
| p.Lys123Asn | Reference | 53 | 0 |
| p.Lys123Asn | Alternate | 0 | 2 |
| p.Met109Ile | Reference | 50 | 0 |
| p.Met109Ile | Alternate | 0 | 6 |
| p.Arg104Gln | Reference | 50 | 0 |
| p.Arg104Gln | Alternate | 0 | 6 |
| p.Lys89Arg | Reference | 56 | 0 |
| p.Lys89Arg | Alternate | 0 | 0 |
| p.Asp84Val | Reference | 55 | 0 |
| p.Asp84Val | Alternate | 0 | 1 |

| | | | |
|---|---|---|---|
| p.Leu81* | Reference | 56 | 0 |
| p.Leu81* | Alternate | 0 | 0 |
| p.Val76Ile | Reference | 43 | 0 |
| p.Val76Ile | Alternate | 0 | 13 |
| p.Thr67Ala | Reference | 7 | 0 |
| p.Thr67Ala | Alternate | 0 | 49 |
| p.Ser40Pro | Reference | 4 | 0 |
| p.Ser40Pro | Alternate | 0 | 51 |
| p.Val6Val | Reference | 56 | 0 |
| p.Val6Val | Alternate | 0 | 0 |
| **SD_SP Sanger - Indels + SNPs** | | | |
| **Variant** | **Allele_DGN** | **Ancestral_allele_SD_SP_Sanger** | **Derived_allele_SD_SP_Sanger** |
| c.-439_-433del | Insertion | 11 | 0 |
| c.-439_-433del | Deletion | 0 | 3 |
| c.-334_-333insACATTCAT | Deletion | 13 | 0 |
| c.-334_-333insACATTCAT | Insertion | 0 | 1 |
| c.-171_-151del | Insertion | 14 | 0 |
| c.-171_-151del | Deletion | 0 | 0 |
| p.Leu81* | Wild-type | 14 | 0 |
| p.Leu81* | Stop gained | 0 | 0 |
| p.Glu202* | Wild-type | 13 | 0 |
| p.Glu202* | Stop gained | 0 | 1 |
| p.Gln254* | Wild-type | 4 | 0 |
| p.Gln254* | Stop gained | 0 | 7 |
| p.Phe217_Glu273del* | Insertion | 11 | 0 |
| p.Phe217_Glu273del* | Deletion | 0 | 3 |
